# Perfect adaptation achieved by transport limitations governs the inorganic phosphate response in S. cerevisiae

**DOI:** 10.1101/2022.05.26.493535

**Authors:** Hon Ming Yip, Shiyu Cheng, Evan J. Olson, Michael Crone, Sebastian J. Maerkl

## Abstract

Cells cope with and adapt to ever-changing environmental conditions. Sophisticated regulatory networks allow cells to adjust to these fluctuating environments. One such archetypal system is the *S. cerevisiae* Pho regulon. When external inorganic phosphate (*P_i_*) concentration is low, the Pho regulon activates, expressing genes that scavenge external and internal *P_i_*. However, the precise mechanism controlling this regulon remains elusive. We conducted a systems analysis of the Pho regulon on the single cell level under well-controlled environmental conditions. This analysis identified a robust, perfectly adapted Pho regulon state in intermediate *P_i_* conditions, and we discovered a hitherto unknown intermediate nuclear localization state of the transcriptional master regulator Pho4p. The existence of an intermediate nuclear Pho4p state unifies and resolves outstanding incongruities associated with the Pho regulon, explains the observed programmatic states of the Pho regulon, and improves our general understanding of how nature evolves and controls sophisticated gene regulatory networks. We further propose that robustness and perfect adaptation are not achieved through complex network-centric control, but by simple transport biophysics. The ubiquity of multi-transporter systems suggests that similar mechanisms could govern the function of other regulatory networks as well.

## Introduction

Cellular regulatory complexity evolved to allow cells to adapt to fluctuating environmental conditions. Extracellular nutrient availability changes over time and cells cope with these fluctuations by adapting their gene expression patterns. Inorganic phosphate (*P_i_*) is one such essential nutrient for which organisms evolved a sophisticated regulatory network [1]. In the budding yeast *S. cerevisiae*, *P_i_* is sensed internally by the cyclin-dependent kinase complex Pho80p and Pho85p. Pho80p/Pho85p are located in the nucleus and transduce *P_i_* levels to their target, the transcriptional master regulator Pho4p [2, 3, 4] through phosphorylation [5, 6]. The phosphorylation state of Pho4p in turn determines whether Pho4p is localized to the nucleus or the cytoplasm. When Pho4p is fully phosphorylated it is localized to the cytoplasm and inactive, whereas it becomes nuclear localized when it is dephosphorylated. Nuclear localization of Pho4p leads to the expression of Pho regulon genes including phosphate transporters, phosphatases, and vacuolar storage regulators. Expression of these genes increases phosphate import and maintains intracellular *P_i_* levels.

This raised the question of how the Pho regulon can activate without entering a futile oscillation cycle in which a *P_i_* flux increase due to Pho-regulon activation leads to an increase in internal *P_i_*, in turn raising *P_i_* levels again above the critical activation threshold and turning the system off, just to be activated once again when the Pho-regulon gene products have been diluted out. The current Pho-regulon model therefore stipulates and necessitates the existence of one or more feedback loops that act to decrease internal *P_i_* concentrations. These negative feedback loops are thought to be Spl2p which binds and inactivates low-affinity transporters [7], and Vtc1-4p which are thought to decrease internal *P_i_* by sequestering and storing *P_i_* in the vacuole [8].

The current model assumes that *P_i_* influx rate depends on extracellular *P_i_* concentration, which in turn means that internal *P_i_* levels vary over a continuum of concentrations [8, 9]. Varying internal *P_i_* concentrations requires a sophisticated control mechanism that constantly adjusts the relative strength of the negative feedback loop [10], which is feasible and has been described in other systems such as osmoregulation [11] and tunable gene expression in the Gal system [12]. Our current understanding of Pho4p also suggests that nuclear Pho4p concentration should track internal *P_i_* concentration in order to achieve the required gene expression tuning [10, 13]. But so far, no differences in nuclear-localized Pho4p levels have been observed and no intermediate states have been identified. It is also counter-intuitive that the model requires the existence of feedback loops that counteract the function of the Pho regulon, which is to increase *P_i_* flux, not decrease it. And finally, one of the main feedback loops suggested to be Vtc1-4p should become inactive under prolonged intermediate phosphate starvation, as *P_i_* sequestration into the vacuole has to cease once vacuoles have reached capacity. Many questions therefore remain in regards to the mechanistic details of how the Pho regulon functions.

We observed that the Pho regulon can attain three clearly defined states as opposed to a continuum of states: an “off” state, a “plateau” state, and a fully activated “on” state. In the off state, when extracellular *P_i_* concentrations are high, all Pho regulon genes are off. The plateau state is a previously observed but hitherto unexplained state in which the entire population of cells are partially activating the Pho regulon [7]. We call this the plateau state because we found that this state is robust to differences in extracellular *P_i_* concentration. The off and plateau states are linked by a bistable region [7]. The third and final state is the fully activated Pho regulon and is observed in conditions of very low or depleted extracelullar *P_i_*. We were also able to quantify nuclear Pho4p localization in these states. Pho4p is highly nuclear localized in the on state, as previously described. But we discovered that Pho4p is partially localized to the nucleus in the plateau state, and that the nuclear concentration of Pho4p in the plateau state is also robust and insensitive to differences in extracellular *P_i_* concentration ([*eP_i_*]). The existence of an intermediate Pho4 nuclear state resolves current incongruities and readily explains the observed Pho-regulon behavior. The observation of perfect adaptation and robustness in the plateau state led us to suggest that *S. cerevisiae* uses a different control mechanism than previously thought. Departing from the generally adopted regulatory network-centric dogma, we suggest that the Pho regulon is governed by simple transporter biophysics.

## Results

### Strain libraries and microfluidic devices

To conduct a systems-level analysis we chose 23 genes known or presumed to be regulated by Pho4p (Fig. 1A, Supplementary Table S1). The promoters of these 23 target genes were cloned upstream of yeast optimized GFP (yoGFP) and integrated into the *LYS2* locus (Fig. 1B, Supplementary Figure S1). We created an “LF” library in which all cassettes use the same generic terminator ADH1te, and an “NN” combined readout / deletion library by replacing these genes in their native loci with yoGFP. To enable characterization of Pho4p localization, we generated a yomScarlet-i - Pho4p fusion in the endogenous locus to preserve native expression levels. We did not observe large differences in target gene expression between the native Pho4p and the N-terminally tagged version, indicating that the fusion protein did not considerably alter Pho4p function (Supplementary Fig. S2).

**Figure 1:**
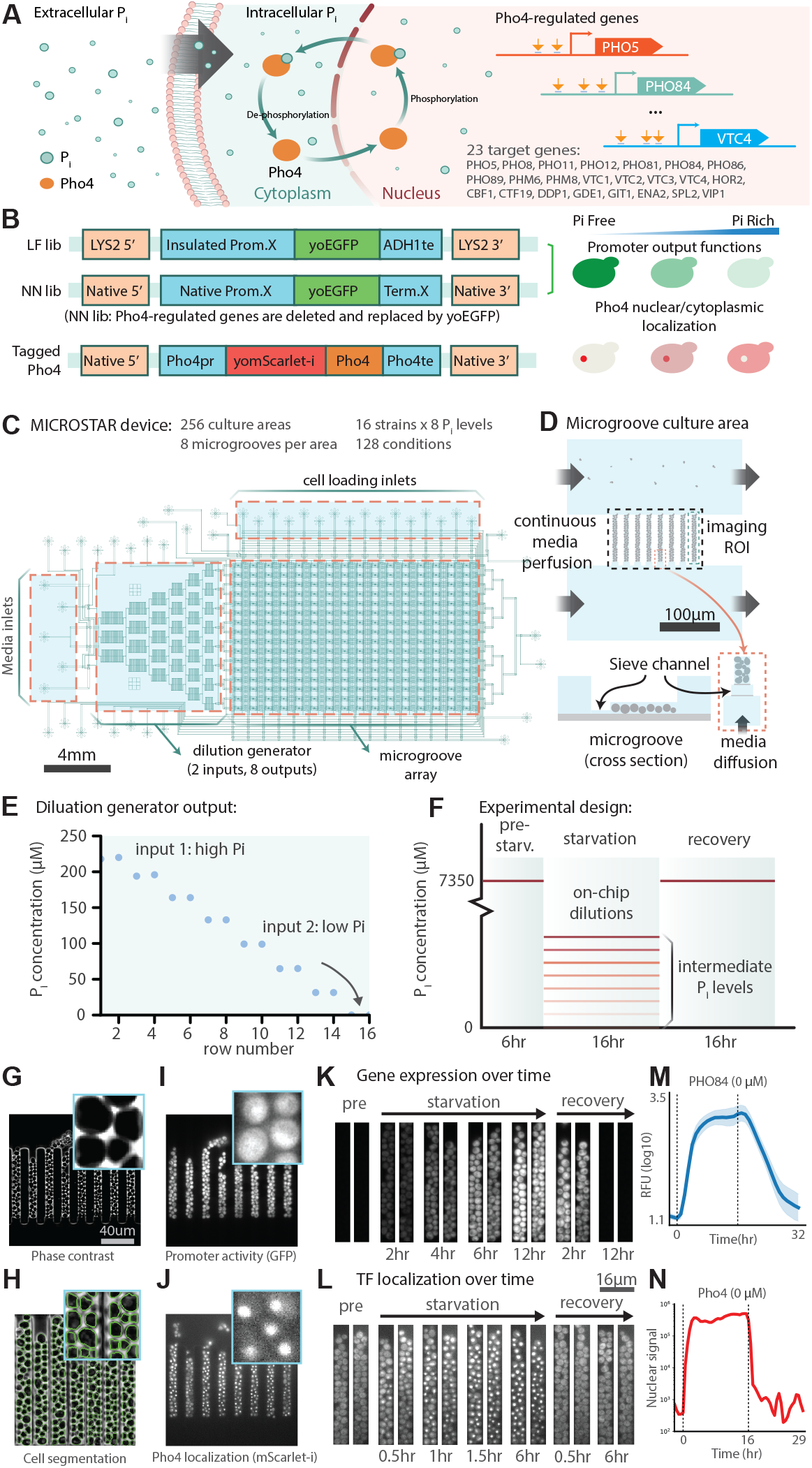
Overview of the *S. cerevisiae* Pho regulon, strain libraries, and MICROSTAR microfluidic device. **A,** *P_i_* is sensed internally by *S. cerevisiae,* leading to phsophorylation of the master regulator Pho4p, which in turn determines whether Pho4p is nuclear localized, or not. When nuclear localized, Pho4p can bind to binding sites (indicated by orange arrows) in its target promoters leading to expression of the Pho regulon genes. **B,** In this study we generated two libraries: LF and NN. The LF library has 23 of the Pho regulon promoters cloned in front of yoGFP and all cassettes are terminated by a common terminator. The NN library has the native genes replaced with yoGFP leading to a combined deletion / reporter library. We also generated a yomScarlet-i-Pho4p strain to track Pho4p localization on the single cell level. **C,** We used two high-throughput microfluidic devices in this study. A 1,152 chamber MCA device (Supplementary Fig. S3) and a MCIROSTAR device with enhanced control over the growth micro-environment. **D,** The MICROSTAR cultures cells in narrow grooves, with the bottom of each groove connected to a media supply channel via a narrow groove. **E,** Both the MCA and MICROSTAR devices are equipped with an on-chip dilution generator to automatically generate 10 and 8 *P_i_* media conditions, respectively. **F,** Automated media switching enables complex experiments: growing strains under rich media conditions, followed by a switch to different *P_i_* starvation conditions, and a switch back to rich conditions. **G-N,** Using time-lapse imaging with high spatio-temporal resolutions returns information on promoter activation and Pho4p nuclear localization.

For large-scale characterization of all strains we employed a previously developed high-throughput microchemostat array (MCA), that allows characterization of 1,152 yeast strains in parallel [14, 15]. We added a microfluidic dilution generator (DG) to automatically generate different extracellular *P_i_* concentrations (Supplementary Figs. S3,S4) [16]. In order to achieve more precise control over growth and environmental conditions we engineered a MICROSTAR (MICROfluidic *S. cerevisiae* Trapping ARray) device, on which yeast cultures are confined to narrow, independent culturing grooves roughly two cell diameters in width (Fig. 1C-D, Supplementary Figs. S5, S6). This design is similar to the “mother-machine” for bacterial cultures [17], but with the added feature that cells are perfused from both ends of the groove, while permitting cell exit only from the top. The MICROSTAR device allows 16 strains to be analyzed under eight media conditions for 128 experiments per device. The DG generates eight media conditions including the high and low media inputs (Figure 1E, Supplementary Figs. S7-9). Media conditions can be changed on the fly so that a standard experiment initially cultures cells under nominal, rich *P_i_* conditions for six hours, followed by a switch to eight different *P_i_* concentrations for 16 hours, followed by a 16 hour recovery period in rich conditions (Fig. 1F). The devices are imaged on an automated microscope with a time resolution of 20-30 minutes (Fig. 1G-L). We can therefore follow promoter activity (Fig. 1M), or assess Pho4p nuclear localization (Fig. 1N).

### Systems analysis of the Pho regulon

To obtain a systems-level overview of the possible states the Pho regulon can attain as a function of [*eP_i_*], we characterized the activity of all 23 promoter strains across both libraries, LF and NN, on the MCA device (Supplementary Figs. S10, S11) and analyzed all LF strains on the MICROSTAR device (Fig. 2, Supplementary Figs. S12, S13).

**Figure 2:**
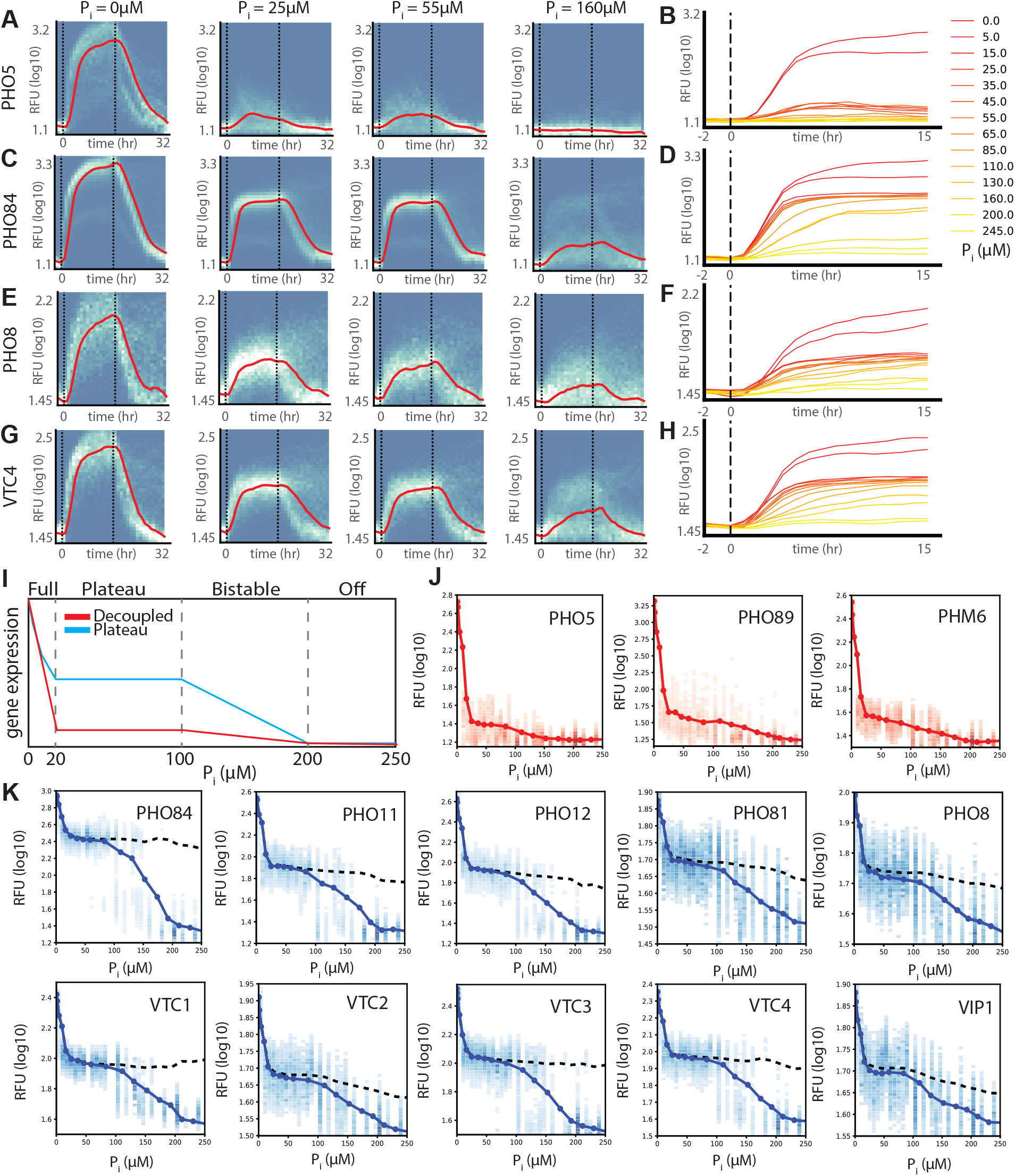
Systems analysis of the Pho regulon on the single-cell level as a function of [*eP_i_*]. **A-B**, *PHO5,* **C-D,** *PHO84*, **E-F,** *PHO8,* and **G-H,** *VTC4* promoter activity measured on a MICROSTAR device over time and different [*eP_i_*]. **A, C, E, G,** Vertical dashed lines indicate when media switches occurred. Graphs show single cell promoter activity distributions as a heat map, and moving median intensities are indicated by red lines. **B, D, F, H,** Median promoter activity levels for all *P_i_* concentrations measured. **I,** Pho regulon promoters can be classified into “decoupled” and “plateau” promoters. **J,** Decoupled promoters: *PHO5, PHO89,* and *PHM6.* **K,** Plateau promoters: *PHO8, PHO11, PHO12, PHO81, PHO84, VTC1-4,* and *VIP1.* Single cell intensity distributions are plotted as heat maps, and solid lines show the median value. Dotted lines indicate median values above a certain threshold to capture only activated cells. For each strain we analyzed over 250,000 cells (Supplementary Figs. S14,S15), across 4-9 devices (Supplementary Table S2)

We were able to map the precise output function of the Pho regulon under defined steadystate conditions. This showed that, rather than a plethora of possible promoter classes and output states, Pho-regulated promoters fall into two classes (Supplementary Fig. S16). One class are the previously described decoupled promoters including *PHO5* (Fig. 2A,B) [18]. Decoupled promoters show no activity in conditions above 200 *μ*M *eP_i_*, very slight activation in intermediate *eP_i_* ranges of 20-200 *μ*M, and very high activation below 20 *μ*M *eP_i_*. The second, and major, class of promoters to which *PHO84* belongs, we call “plateau” promoters (Fig. 2C-H). Plateau promoters remain inactive above 200 *μ*M *eP_i_*, but enter an intermediate activation state in the range of 20-200 *μ*M *eP_i_*. Strikingly, these promoters exhibit a surprisingly invariable expression level over an order of magnitude range of [*eP_i_*]. In other words, these promoters exhibit perfect adaptation and robust control. When [*eP_i_*] fell below 20 *μ*M, expression of these promoters increased further. It should be noted that even decoupled promoters display a low-level plateau behaviour.

The Pho regulon can therefore attain three distinct states or programs (Fig. 2I, Supplementary Fig. S17). When [*eP_i_*] is above 200 *μ*M the entire Pho regulon remains in the off state. In a relatively narrow range of 100-200 *μ*M *eP_i_* the Pho regulon exhibits bistability, where cells can be either in the off or plateau state [7]. In the range of 20 - 100 *μ*M *eP_i_* the Pho regulon exhibits a high degree of robustness in that promoter activity is largely independent of [ePi]. This plateau region functionally extends all the way to the upper end of the bistable region at 200 *μ*M *eP_i_*. At [*eP_i_*] below 20 *μ*M the Pho regulon is fully activated, with decoupled promoters turning on and plateau promoters further increasing in activity above their plateau levels.

The existence of a plateau region is puzzling. Pho4p is known to be localized to either the cytoplasm or nucleus and no intermediate states of Pho4p nuclear localization have been described thus far. It is therefore not clear how promoter decoupling can be achieved at intermediate phosphate conditions, nor is it at all understood how the system achieves a robust, perfectly adapted state.

### Pho4p nuclear localization

We attempted to quantify Pho4p localization under steady-state conditions over the entire functionally relevant *eP_i_* range (Fig. 3). Nuclear localization of Pho4p was clearly visible under full starvation conditions. In intermediate *eP_i_* conditions, nuclear localization of Pho4p was not as obvious, but we were able to observe a detectable level of Pho4p nuclear localization (Fig. 3A).

**Figure 3:**
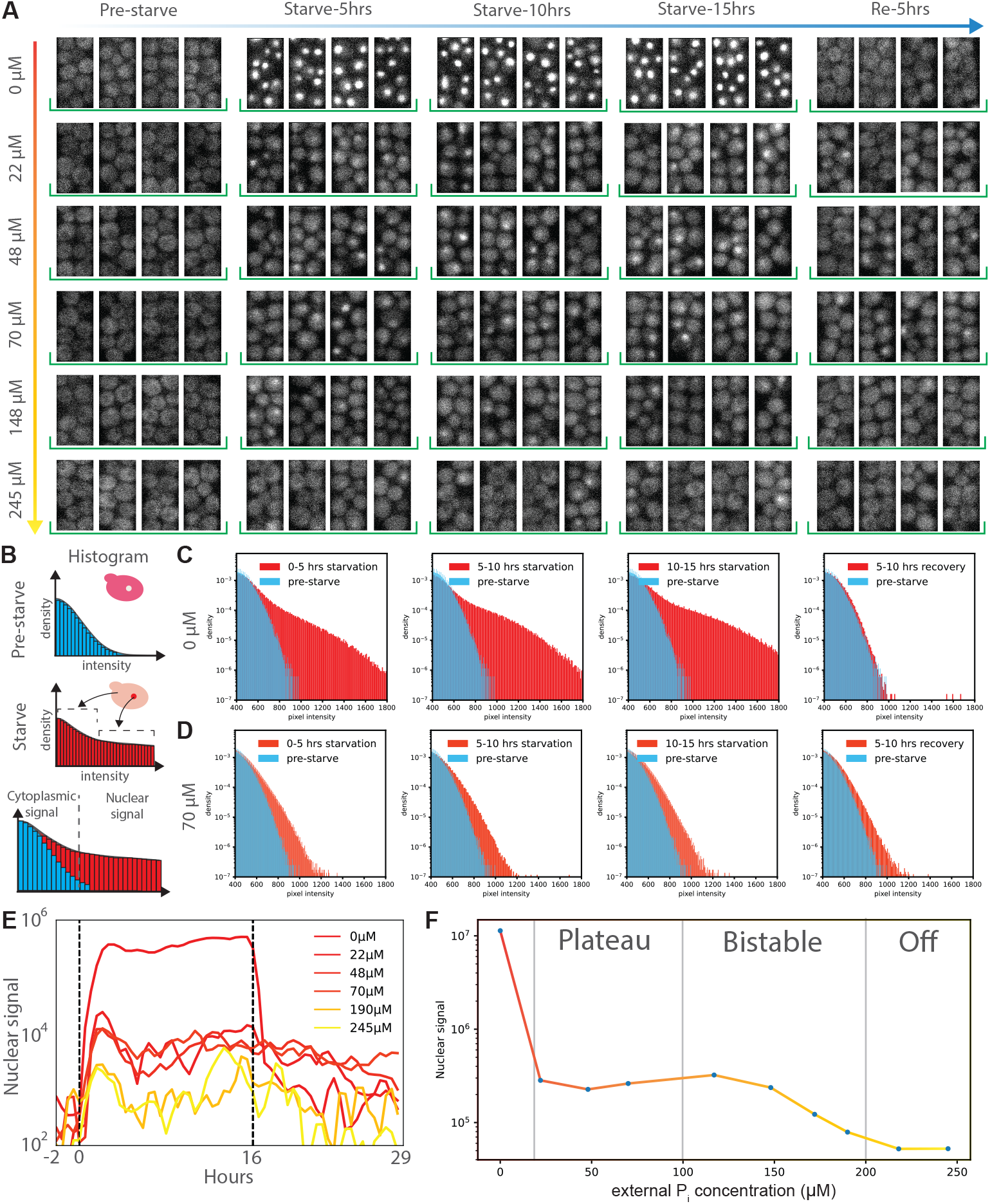
Pho4p nuclear localization as a function of [*eP_i_*]. **A,** Time-lapse image series showing single-cell Pho4p nuclear localization. No Pho4p is visibly nuclear localized pre-starvation. Upon shift to lower [*eP_i_*], nuclear localization is clearly visible for 0 *μ*M *eP_i_*. In the range of 22 - 245 *μ*M *eP_i_* low levels of nuclear Pho4p are visible. **B,** Schematic explanation of the image histogram based analysis. When Pho4p becomes nuclear localized it also becomes locally more concentrated, giving rise to higher intensity pixels. **C-D,** Image histograms showing pixel intensity distributions for 0 *μ*M and 77 *μ*M *eP_i_*. In every plot the blue histogram shows the pixel intensity distribution prior to starvation for reference. The red histograms show either the intensity distribution during starvation or during recovery as indicated. **E,** Pho4p nuclear localization plotted over time for different [*eP_i_*]s. **F,** Pho4p nuclear localization at steady state under different [*eP_i_*]. For each concentration bin we analyzed between 8-30 culturing chambers, 11 - 51 grooves, across 1-4 devices (Supplementary Fig. S18)

Although nuclear Pho4p signal was rather low at intermediate conditions, we attempted to quantify Pho4p localization. We used a simple, but effective analysis method based on the fact that when Pho4p is localized to the nucleus it also becomes locally more concentrated (Fig. 3B).

Localization of Pho4p to the nucleus should thus be reflected in an increased number of higher intensity pixels observable in image histograms. For low [*eP_i_*] the transition of Pho4p to the nucleus is visually obvious and results in very pronounced histogram tails (Fig. 3C). This histogram method is sufficiently sensitive to detect nuclear Pho4p at intermediate and high [*eP_i_*] (Fig. 3C, Supplementary Fig. S19). For example, at 70 *μ*M *eP_i_*, a clear difference in Pho4p localization can be seen between the pre-starvation state and the starvation state (Fig. 3D).

In full starvation conditions Pho4p localized to the nucleus to a large degree (Fig. 3E, Supplementary Fig. S20). Intermediate conditions gave rise to low but measurable Pho4p nuclear localization, while rich *P_i_* conditions showed very slight or no Pho4p nuclear localization. Strikingly, Pho4p nuclear localization in intermediate conditions also appeared to be robust and invariant to [*eP_i_*]. We thus plotted Pho4p nuclear localization at steady state as a function of [*eP_i_*] (Fig. 3F), showing that Pho4p is only partly nuclear localized in the plateau region. Complete nuclear localization only occurs in the on state of the system under low [*eP_i_*]. Partial nuclear Pho4p localization in the plateau state is furthermore largely constant and insensitive to [*eP_i_*].

The output of the Pho regulon therefore appears to be correlated with, and likely determined by, Pho4p nuclear concentration. In the off state, no, or very small quantities of Pho4p are nuclear localized, leading to a nuclear concentration that is insufficient to activate any Pho regulon promoters. In the plateau state, Pho4p is partially nuclear localized, leading to a nuclear Pho4p concentration that is sufficient to activate plateau promoters but remains below the threshold to fully activate decoupled promoters. Under low *eP_i_* conditions, Pho4p is largely nuclear, giving rise to a high nuclear concentration that is sufficient to activate all promoter classes.

### Transport-centric model

Given these insights we set out to derive a model that could explain the observed phenotypes. Current models assume that *P_i_* influx rate varies with [*eP_i_*]. At the same time, we and others have not observed measurable growth rate decreases at intermediate [*eP_i_*]. We posit that *P_i_* consumption or assimilation rate has to be proportional to growth rate, which means that at constant growth rate, *P_i_* consumption rate is also constant. Under these conditions: a *P_i_* influx rate that varies with [*eP_i_*], and a constant *P_i_* consumption rate; internal *P_i_* concentrations ([*iP_i_*]) should track [*eP_i_*]. A continuum of [*iP_i_*] makes it difficult to explain the observed nuclear localization behaviour of Pho4p and general behaviour of the Pho regulon, aside from appearing to be a suboptimal control strategy.

We therefore asked whether, instead of a control mechanism established by a finely tuned network architecture, the Pho regulon could be governed by an entirely different mechanism. This led us to look more closely at the transporter system, which consists of constitutively expressed low-affinity transporters (Pho90p and Pho87p), and high-affinity transporters (Pho84p and Pho89p) that are expressed at intermediate and low [*eP_i_*]. The Pho regulon expresses Spl2p at intermediate and low [*eP_i_*], which is thought to bind to low-affinity transporters turning them off. The existence of the dual transporter system has been suggested to provide a competitive advantage, albeit with the same supposition that *P_i_* influx varies with [*eP_i_*] [9]. More recently Bosdriesz and colleagues put forth a transport carrier model as opposed to the standard Michaelis-Menten model [19]. They argue that the carrier model explains the existence of different affinity transporters, as transporters of different affinities will achieve optimal transport rates at different external substrate concentrations.

Another important consequence of the carrier model that has not been described thus far is the fact that it predicts that [*iP_i_*] are robust and insensitive to [*eP_i_*]. This can be easily seen when plotting the import rate (J) as a function of both internal and external substrate concentration (Fig. 4A). The import rate function J collapses on a single curve when external substrate concentrations are sufficiently high. The system is also stable, as the *P_i_* import rate J will always tend towards matching the cellular *P_i_* consumption rate. Choosing a constant import rate J, one can then plot internal versus external *P_i_* concentration which shows that internal substrate concentration is stable and robust to [*eP_i_*], and below a critical threshold, [*iP_i_*] will tend to zero (Fig. 4B). A Michaelis-Menten model on the other hand stipulates that above the critical threshold [*iP_i_*] will always track [ePi] (Fig. 4C-D).

**Figure 4:**
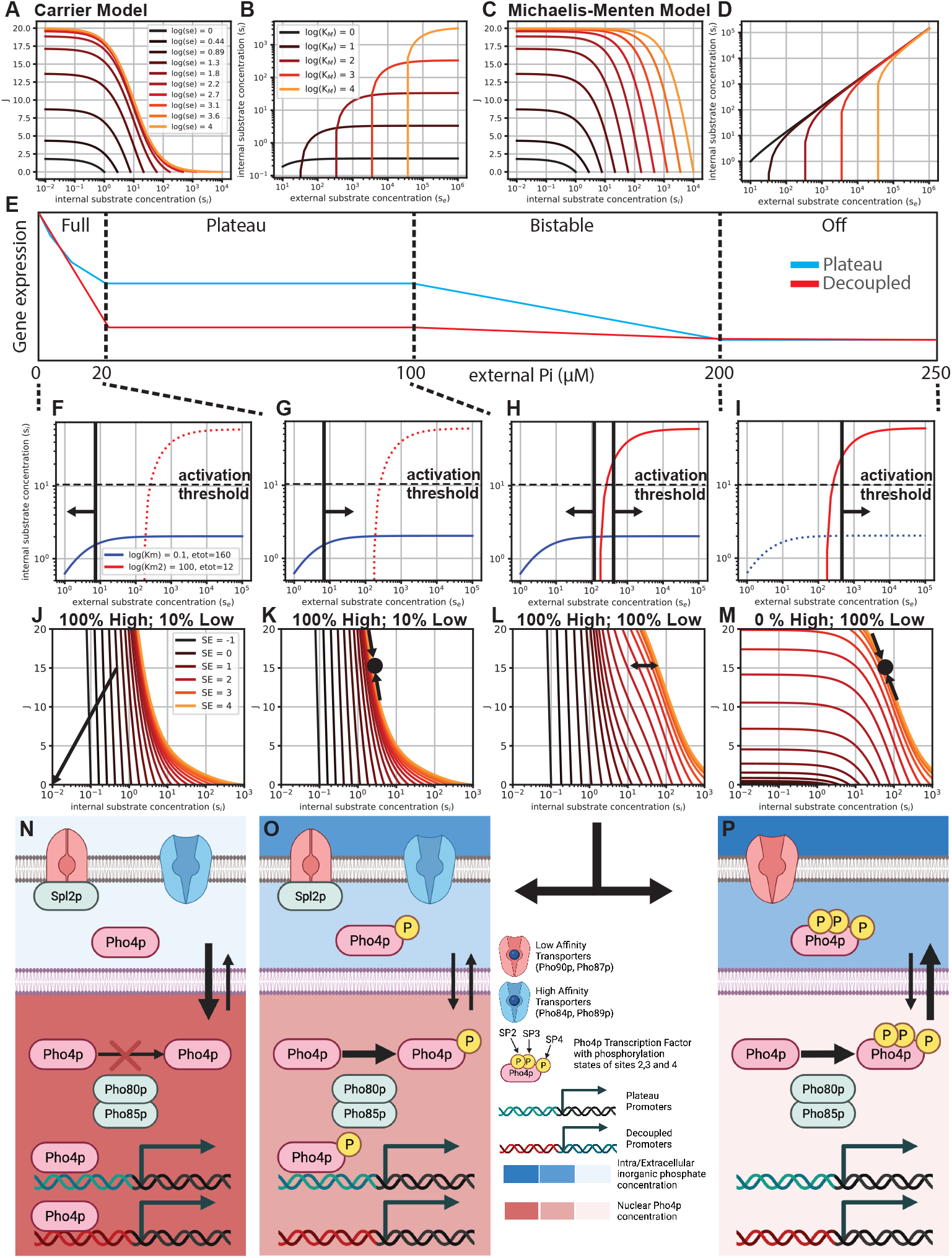
Transport-centric Pho regulon model. **A,** Carrier model steady-state import rate (J) as a function of external and internal substrate concentration [19]. Parameters were: *k*_2_ = 4; *k*_4_ = 4; *K_d_* = 10; *e_t_* = 10. **B,** Carrier model: internal substrate concentration as a function of external substrate concentration at an import rate *J* = 15. **C,** Michaelis-Menten model for comparison. Parameters were the same as for the carrier model. **D,** Michaelis-Menten model: internal substrate concentration as a function of external substrate concentration at an import rate J=15. **E,** Schematic representation of the Pho regulon output function with its three major states (fully active, plateau, off) and the transition (bistable) state demarcated by dashed vertical lines. **F-P,** Each of the states can be mapped to a specific cellular state. **I, M, P,** At high [*eP_i_*] the Pho regulon is turned off. Low-affinity transporters are active, setting a robust [*iP_i_*] that is above the activation threshold. Pho4p is fully phosphorylated and localized to the cytoplasm. No Pho regulon genes are expressed. **H, L,** When [*eP_i_*] approach the critical value set by the low-affinity *KD* (or K_M_) value a bistable state exists in which some cells remain in the off state while others enter the plateau state. **G, K, O,** When [*eP_i_*] is below the critical import threshold of the low-affinity importers, [***iP***i] falls below the Pho regulon activation threshold and Pho4p becomes partially dephosphorylated, leading to low levels of nuclear localization. This nuclear Pho4p concentration is high enough to partially activate the Pho regulon (plateau-promoters). High-affinity transporters are expressed, which establish a new, robust [*iP_i_*] below the activation threshold. Spl2p is also expressed which turns off the low-affinity transporters. **F, J, N,** When [*eP_i_*] falls below the critical import threshold defined by the high-affinity transporter *KD* (or *K_M_*) value, [*iP_i_*] falls, and tend towards zero, leading to full dephosphorylation and full nuclear localization of Pho4p, which in turn activates the entire Pho regulon (plateau and decoupled promoters).

A simple carrier model consisting of two transporters of different affinities is consistent with experimental observations of the Pho regulon, including the apparent robustness to [ePi] (Fig. 4E-P). In the off-state, when [*eP_i_*] is high, low-affinity transporters are active and set an [*iP_i_*] that lies above the activation threshold (Fig. 4I,M,P). As [*eP_i_*] approaches the critical threshold for the low-affinity transporters, a bistable region exists in which some cells remain in the off state, whereas others may enter the plateau state (Fig. 4H-L). At intermediate [*eP_i_*], [*iP_i_*] levels fall below the activation threshold, leading to partial phosphorylation and partial nuclear localization of Pho4p. This in turn activates Spl2p which inactivates existing low-affinity transporters. Concomitantly, Pho4p expresses the high-affinity transporters, which establish a new stable state of constant [*iP_i_*], that lies below the activation threshold, but above the threshold at which growth limitations occur (Fig. 4G,K,O). The cell therefore has to cope and adapt to only two [*iP_i_*], which can be relatively similar (only limited by the transfer function of the Pho80p/85p kinase system), and are constant and robust to changes in [*eP_i_*]. When [*eP_i_*] falls below the critical level for the high-affinity transporter, [*iP_i_*] will tend to zero leading to complete dephosphorylation of Pho4p and complete nuclear localization, which activates all Pho regulon genes (Fig. 4F,J,N). The model also predicts hysteresis when returning from intermediate [*eP_i_*] to high [*eP_i_*] since the high-affinity transporters are turned off and the high-affinity transporters will keep [*iP_i_*] constant. This hysteresis has been previously seen and described as commitment [20], and we also observed hysteresis in promoter activity (Supplementary Fig. S10) and Pho4p nuclear localization (Fig. 3D-E, Supplementary Fig. S19).

## Discussion

The *S. cerevisiae* Pho regulon has been studied extensively but the precise mechanism governing the regulon remained unclear. The current Pho regulon model stipulates that it is regulated by feedback loops. It furthermore is thought that *P_i_* import rates vary with [ePi] [8, 20], leading to changing [*iP_i_*] that are transduced to the master regulator Pho4p. Nuclear Pho4p concentration is therefore also thought to change as a function of external or internal *P_i_* concentration following a standard transfer function [13], although only two Pho4p localization levels had thus far been described: a fully nuclear state and a fully cytoplasmic state. It was therefore argued that affinity or kinetic differences due to phosphorylation of site 6 must play a role in the differential expression observed between *PHO5* and *PHO84* at intermediate *P_i_* levels leading to an activity rather than a concentration model, but it was also acknowledged that an unknown mechanism must exist that contributes to this differential expression [13].

By conducting a systems-level analysis of the Pho regulon we found that the Pho regulon can attain three well-defined states: a fully activated state, a plateau state, and an off state, with a bistable transition state between the off and the plateau state. The plateau state is perfectly adapted and robust to external *P_i_* concentration changes over a range of 20 - 100 *μ*M, or 20 - 200 *μ*M if the bistable region is included. We characterized nuclear localization of Pho4p over a wide range of external *P_i_* concentrations and discovered that, rather than attaining two nuclear localization states, Pho4p can attain three distinct nuclear localization states. In the plateau region Pho4p is partially localized to the nucleus, and this localization level is robust to external *P_i_* concentration.

This intermediate nuclear localization state of Pho4p readily explains the observed states of the Pho regulon and provides an appealing solution of how the Pho regulon is designed and tuned on the promoter level. The original model of Pho4p with only two nuclear localization levels (zero and full) required an “activity” model in order to explain observed differential gene expression. Alternatively, a continuum of nuclear Pho4p concentration could have explained the differential expression, but failed to predict the observed perfectly adapted and robust plateau state. Three nuclear Pho4p concentration states on the other hand readily account for the three programmatic output states of the Pho regulon and the three nuclear Pho4 concentration states can be explained via the phosphorylation states and phosphorylation preferences of sites 2,3 and 4 (Fig. 4N-P) [6]. The effect of site 6 is less clear but it is possible that although not necessary for differential expression of plateau and decoupled promoters in intermediate phosphate conditions [13], that it either plays a role in preventing aberrant expression of plateau promoters under non-starving conditions, or provides increased dynamic range between starved and intermediate expression levels by tuning the interaction strength with Pho2p.

Having three defined nuclear Pho4p concentrations of zero, intermediate, and high, also simplifies Pho regulon promoter evolution. Decoupling is used to prevent promoter activation in the plateau region, which is easily accomplished because nuclear Pho4p concentrations are sufficiently low to prevent outcompeting nucleosomes. Only when nuclear Pho4p concentrations are high can Pho4p compete with nucleosomes leading to full promoter activation. Furthermore, expression levels can be easily tuned as promoter binding site affinity can evolve to two specific Pho4p concentrations as opposed to coping with a continuum of nuclear Pho4p concentrations. This is an effective and simple approach to gene regulatory network design and is instructive for the design of complex synthetic gene regulatory networks.

The observation of three nuclear Pho4p states raises the question of how these states are established in the first place, and how a robust and perfectly adapted plateau state can exist. We excluded the possibility that the Pho regulon uses a network feedback approach to achieve perfect adaptation because we did not identify any components that act as a negative feedback nor found any negative feedback components that track [*iP_i_*] or [*eP_i_*]. It is possible that the signaling cascade leading to Pho4p phosphorylation establishes three distinct Pho4p phosphorylation states as a function of continually varying [*iP_i_*]. But we consider varying [*iP_i_*] to be inherently flawed and explored whether it might be physically possible to establish robust [*iP_i_*] instead.

A standard Michaelis-Menten transporter model predicts that internal substrate concentration tracks external substrate concentration under steady-state conditions. This seems to be a sub-optimal strategy as cells would need to function optimally over a wide range of [*iP_i_*]. A continuum of *iP_i_* levels would also necessitate complex regulatory control to achieve the observed Pho regulon program. A carrier transport model recently explained the existence of different affinity transporters [19]. We realized that the carrier model also predicts that internal substrate concentrations are constant and robust to external substrate concentrations. Therefore a simple system of two transporters with different affinities could establish two robust [*iP_i_*], which then simply need to establish two Pho4p phosphorylation states through the Pho80p/85p kinase system. These two phosphorylation states in turn determine whether Pho4p is sequestered to the cytoplasm, or nuclear localized to intermediate levels. When *iP_i_* levels fall further, Pho4p is completely dephosphorylated and fully nuclear localized. This model is compatible with all known Pho regulon phenotypes, including the observation of commitment or hysteresis that occurs when cells are returned to rich *eP_i_* conditions. It eliminates the inconsistencies of the current model, particularly in regards to the existence of the feedback loops, and predicts a stable plateau region, which has hitherto not been described. The model also predicts the establishment of homeostatic conditions. Therefore we suggest that complex gene regulatory networks such as the Pho regulon could be governed by simple control mechanisms based on transporter biophysics.

This work provides insight into how nature evolved simple, yet robust solutions to controlling sophisticated gene regulatory networks. The Pho regulon exhibits a complex response consisting of three distinct states: off, intermediate activation, and full activation. The differences between the intermediate and full activation state are complex and involve not only differences in activation levels, but also differences in the set of promoters being activated in each state. These states are in turn determined by 3 corresponding nuclear concentration levels of Pho4p. Therefore, a combination of Pho4 nuclear concentration states and promoter decoupling through nucleosome occupation of TF binding sites can fully account for this complexity. This example demonstrates that distinct and well-defined nuclear TF concentrations can be established and give rise to complex programs. It is thus likely that TF concentration is instrumental in other regulatory networks as well. Further, we posit that overall system control is not achieved by complex regulatory network architectures, but rather by simple transporter biophysics. It will be interesting to determine whether nature uses similarly simple, yet elegant solutions, to regulate other complex systems such as the glucose regulatory network, which also uses a combination of high and low affinity transporters.

## Acknowledgments

This work was supported by the European Research Council under the European Union’s Horizon 2020 research and innovation program grant 723106.

## Competing interests

None declared.

## Author contributions

S.C., M.C., and E.J.O. generated the strains used in this study. H.M.Y. designed the microfluidic devices and performed all on-chip experiments. E.J.O. wrote the image analysis pipeline. E.J.O and H.M.Y performed data analysis. S.J.M., H.M.Y., S.C., and E.J.O wrote the manuscript with input from all authors. S.J.M. conceived and directed the study.

## Notes

### Competing Interest Statement

The authors have declared no competing interest.

### Summary of Updates

This version of the document contains several changes in response to reviewer comments and suggestions.

