## Supplementary Information for "Perfect adaptation achieved by transport limitations governs the inorganic phosphate response in S. cerevisiae"

Institute of Bioengineering
School of Engineering
École Polytechnique Fédérale de Lausanne

October 24, 2022

[1]These authors contributed equally

**2 Supplementary Figures** **33**

**3 Supplementary Tables** **54**

### Chapter 1

### Materials and Methods

#### 1.1 Plasmids, Strains and Media

##### 1.1.1 Plasmid Library Construction

We used NEB 10-beta Competent *E. coli* (High Efficiency) (New England Biolabs, Cat #C3019H) for all plasmid cloning following the transformation protocol from the manufacturer. We conducted bacterial selection and growth in LB plates or LB medium at 37°C supplemented with appropriate antibiotics (chloramphenicol 34 $\mu$g mL$^{-1}$, ampicillin 100 $\mu$g mL$^{-1}$, or kanamycin 50 $\mu$g mL$^{-1}$). For the NN library (native locus, native terminator), the integration cassettes consisted of the native promoter sequence (1500 bp), yeast codon optimized enhanced green fluorescent protein (yoEGFP), and native terminator sequence (500 bp). The native ORFs were replaced by a yeast codon optimized enhanced green fluorescent protein (yoEGFP) using homologous recombination. For the LF library (*LYS2* locus, *ADH1* terminator), the integration cassettes consisted of two *LYS2* homology arms (each 500 bp), a neutral insulation sequence (200 bp), the native promoter sequence (1500 bp), yoEGFP, and ScADH1 terminator (225 bp). We designed different sgRNAs (Table S6) targeting the native loci and *LYS2* Locus and constructed corresponding sgRNA integration plasmids. These DNA sequences were generated from PCR or ordered as DNA fragments (IDT DNA). All integration cassettes were constructed via BsaI or BsmBI Goldengate/TypeIIs assembly of the DNA fragments. Goldengate reactions are prepared as following: 50 ng of backbone plasmid, 150 ng of insertion plasmid, 2 $\mu$L of 10x T4 ligase buffer (NEB), 2 $\mu$L of T4 ligase (NEB), 1 $\mu$L of BsaI or BsmBI (NEB), and adding water to a final volume of 20 $\mu$L. Thermocycler setup: (BsmBI assembly: 45°C for 2 min; BsaI assembly: 42°C for 2 min, 16°C for 5 min) x 25 cycles, 60°C for 10 min, 80°C for 20 min, hold at 4°C forever.

##### 1.1.2 Yeast Strain Construction

For yeast cloning, we used CRISPR/Cas9 assisted markerless integration by homologous recombination (Figure S1) using the lithium acetate/polyethylene glycol (PEG) method [1]. The GFP-TAG strains, including, Pho87 (tag), Pho90 (tag), and SPL2 (tag), were obtained from a commercially available yeast GFP Clone Collection (Thermo Fisher Scientific, US) [2]. Two yeast strain libraries, including NN, and LF, were constructed for this study. The sgRNA plasmids, CRISPR/Cas9 plasmid, and integration cassettes were sequence-verified and digested by EcoRV, BsmBI, and NotI for yeast integration, respectively. EcoRV digestion was set up as follows: DNA 1 $\mu$g, 10X NEBuffer r3.1 5 $\mu$L, EcoRV 1 $\mu$L, nuclease-free water to 50 $\mu$L, incubate at 37°C for 1 h and 65°C for 20 min. BsmBI digestion was set up as follows: DNA 1 $\mu$g, 10X NEBuffer r3.1 5 $\mu$L, BsmBI-v2 1 $\mu$L, nuclease-free water to 50 $\mu$L, incubate at 55°C for 1 h and 80°C for 20 min. NotI digestion was set up as follows: DNA 1 $\mu$g, 10X CutSmart buffer 5 $\mu$L, NotI-HF 1 $\mu$L, nuclease-free water to 50 $\mu$L, incubate at 37°C for 1 h and 65°C for 20 min. After DNA purification (ZYMO DNA clean and concentrator-25 kit), they were transformed and integrated into the parental strain (BY4741) by homologous recombination using the lithium acetate/polyethylene glycol (PEG) method. Briefly, the parental strain BY4741 was grown in the yeast extract peptone dextrose (YPD) medium over night at 30°C and shaking at 250 rpm. After overnight culture, the yeast cells were back diluted to OD 0.175 in 5 mL fresh YPD medium. The OD600 reached around 0.75 after incubation for 4-5 hours. We pelleted the cells by centrifuging for 10 min at 2000 rpm. While harvesting cells, we prepared the lithium acetate transformation solution and DNA mixture solution. For each transformation, we prepared 306 $\mu$L chemical solution, including 260 $\mu$L PEG3350 (50% w/v), 36 $\mu$L Lithium acetate solution, and 10 $\mu$L ssDNA (10 mg $mL^{-1}$). In the meantime, we prepared 54 $\mu$L DNA mixture solution for each transformation, including 1 $\mu$g integration cassette, 100 $\mu$g Ca9/URA3 cassette, and 200 $\mu$g sgRNA cassette. After centrifuging, we washed the yeast cells with 2.5 mL water once and with a 100 mM lithium acetate solution twice. Afterwards, the yeast cells were suspended in the DNA mixture solution (54 $\mu$L) first and then mixed with the lithium acetate transformation solution (306 $\mu$L). After incubation at 30°C for 30 min, we incubated the yeast cell transformation mixture at 42°C for a further 15 min in a water bath. We centrifuged the cells at 8000 rpm for 2 min and removed the supernatant. We re-suspended the cells in 200 $\mu$L water and plate 50 $\mu$L on the selection plates. We conducted transformant selection on synthetic complete uracil dropout (SC-URA) agar plates and cured the selected colonies on 5-FOA supplemented synthetic complete (SC 5-FOA) plates. We screened transformants for correct integration by colony PCR and verified the PCR product by sanger sequencing.

To create the mScarletI-Pho4p strain, we created a mScarletI-Pho4p plasmid with the Pho4p promoter, N-terminus tagged mScarleti-Pho4p, and the Pho4p terminator using a BsmBI assembly. The mScarleti-Pho4p plasmid, CRISPR/Cas9 plasmid, and two sgRNA plasmids (targeting the Pho4p chromosomal site), were sequence verified and digested by Not-I, BsmBI, and EcoRV for integration, respectively. After DNA purification, they were transformed into the parental strain (BY4741) using the lithium acetate/polyethylene glycol (PEG) method. We conducted the transformant selection and curing on SC-URA and SC 5-FOA plate, respectively. We screened transformants for correct integration by colony PCR and verified the PCR product by sanger sequencing.

##### 1.1.3 Media

To prepare Lysogeny Broth (LB) medium, we dissolved 25 g LB powder (MP, 3002022) in deinonized water to a final volume of 1 L. To prepare LB agar plates, we dissolved 25 g LB powder, 20 g Agar (Alfa, 10752-36) in deinonized water to a final volume of 1 L. The medium was autoclaved before adding antibiotics. To prepare YPD medium, we dissolved 50 g YPD broth (Sigma, Y1375) in deionized water to a final volume of 1 L. To prepare YPD agar plates, we dissolved 50 g YPD powder, 20 g Agar (Alfa, 10752-36) in deinonized water to a final volume of 1 L. The YPD medium was autoclaved before use. To prepare SC-URA plates, we added 1.92 g yeast synthetic dropout medium supplement (without uracil) (Y1501-20G), 6.7 g Yeast Nitrogen Base without amino acids (Y0626-1KG), and 20 g Agar in deionized water to 960 mL and after autoclaving added 40 mL 50% filter sterilized glucose solution (Corning 500 mL Vacuum Filter/Storage Bottle System, 0.22 $\mu$m Pore, Cat#431097) to a final volume of 1 L. To prepare the 5-FOA plates, we added 1.92 g yeast synthetic dropout medium supplement (without uracil) (Y1501-20G), 6.7 g Yeast Nitrogen Base without amino acids (Y0626-1KG), and 20 g Agar in deionized water to 880 mL and after autoclaving added 40 mL 50% filter sterilized glucose solution, 66 mL 760 $\mu$g mL$^{-1}$ uracil solution, 10 mL 10% 5-FOA solution (EZSolutionTM, 00386260) to a final volume of 1 L.

#### 1.2 MICROSTAR Device

##### 1.2.1 Device Design

The MICROSTAR (MICROfluidic *S. cerevisiae* Trapping ARray) device was designed in Autocad (Autodesk, U.S.A.) and is fabricated by standard multi-layer soft lithography [3] to produce a two layer device consisting of a control and a flow layer.

The MICROSTAR device consists of 256 culture areas, each area in turn contains 8 parallel culture grooves 8 $\mu$m in width or roughly 2 *S. cerevisiae* cell diameters. The device contains 2,048 cell grooves in total. Each groove is 3.8 $\mu$m in height to force cells to grow in monolayers. Each culture groove is connected to two parallel media perfusion channels. On the bottom of each groove we created a narrow sieve channel that is 1.2 $\mu$m in height, 11 $\mu$m wide, and 5.5 $\mu$m long. This allows media to perfuse the bottom as well as the top of each culture groove, but allows cell escape only from the top. This design creates a well controlled culture area proximal to the sieve channel. The MICROSTAR device contains 16 columns and 16 rows. Each column can be individually loaded with different yeast strains. The 16 rows in turn are divided into 8 individually addressable rows to provide different media conditions, generated by an upstream dilution generator.

The dilution generator [4] is designed to perform on-chip dilutions from two inlets to 8 outputs. The channel width of the dilution generator was designed to be 20$\mu$m, which greatly reduces the required dilution time. The dilution generator can either be used with a single medium inlet, in which case all DG output channel media conditions are identical and equal the media input. To generate different concentrations, two media formulations are introduced from two different inlets, which are combined in the DG to generate 8 unique output formulations. The media switching unit enables changing the inorganic phosphate concentration during the experiment.

##### 1.2.2 Mold Fabrication

The control and flow molds for the MICROSTAR chip were fabricated using conventional photolithography techniques in the center of micronanotechnology at EPFL.

The CAD file of the device was converted to CIF format for mask production. Due to PDMS shrinkage, the control and flow layer were re-scaled to 103.5 percent and 100.5 percent, respectively. Chrome masks with photoresist AZ351 coating were used for mask production. They were written on a Heidelberg Instruments VPG200 (Heidelberg Instruments Mikrotechnik GmbH, Germany).

Masks were then processed in a mask developer (Hamatech HMR900) using standard development, chrome etching, resist stripping, and final rinse dry parameters. The completed masks were then ready to use after overnight drying.

For flow layer mold fabrication, a 4-inch silicon wafer was cleaned at 500W for 7 minutes in an oxygen plasma cleaner Tepla 300 (PVA TePla, Germany). The wafer was then ready to be processed using standard SU-8 fabrication process consisting of (1) photoresist coating, (2) soft bake (SB), (3) exposure, (4) post-exposure bake (PEB), (5) development, and (6) hard bake (HB). The parameters of the process, materials, and the machines are listed in Tables S9, S10, and S11. No development after post-exposure baking was performed for the sieve layer. The photoresist for the chamber layer was directly coated on top of the sieve layer, and both layers were developed together after the chamber layer PEB. This measure prevented the sieve layer structure from negatively affecting coating of the chamber layer which otherwise would cause uneven coating. However, this measure was not applied to the dilution generator layer because: (1) photoresist became more resistant to developer after 2/3 rounds of long soft bake and post-exposure bake that made some parts undeveloped, and (2) DG layer has no overlap with the sieve and chamber layer and so the influence was minimal. After finishing all three SU-8 layers, the wafer was then patterned using a positive photoresist AZ 10XT-60 (MicroChemicals, U.S.A.). The reason for using this type of photoresist is that it is able to generate rounded structures through annealing which can be completely closed by "Quake style valves". The fabrication procedure includes (1) HMDS treatment, (2) photoresist coating, (3) baking, (3) exposure, (4) relaxation, (5) development, and (6) annealing. The details are also listed on Tables S9, S10, and S11. To round the AZ flow channels, the wafer was annealed on a hotplate at 115°C for 1 hour.

The control mold was fabricated using a standard SU-8 workflow as it consists of only a single layer. The parameters of the process, materials, and the machines are listed in Tables S9, S10, and S11. After completion of the molds, they were inspected under a microscope to ensure features were fully developed and the height of each layer was measured using a surface profiler Bruker Dektak XT (Bruker, Germany). The feature heights of the flow mold are 1.2$\mu$m (sieve), 3.8$\mu$m (chamber), 15$\mu$m (flow channel) and 40$\mu$m (dg), and the feature height on the control mold is 20$\mu$m. Both newly fabricated control and flow molds were treated with TMCS (trimethylchlorosilane) vapor in a wafer box inside a fume hood overnight before the first usage.

##### 1.2.3 Device Fabrication

Device fabrication includes (1) PDMS preparation, (2) spin coating (for flow layer), (3) 1st baking, (4) hole punching, (5) alignment, (6) final bake, and (7) bonding to glass. Before chip fabrication, both control and flow wafers were treated with TMCS vapor in a wafer box inside a fume hood for half an hour. Then, two PDMS ratios were prepared. For the control layer, 48 g of PDMS (with a base to curing agent ratio of 5:1) was prepared and mixed in a Thinky mixer ARE-250 CE (Thinky, U.S.A) for 1 min at 2,000 rpm and then de-foamed for 2 min at 2,200 rpm. The control mold was placed in a 6-inch glass petri dish coated with aluminium foil. The mixed PDMS was poured on the control mold and then degassed in a vacuum chamber for half an hour. After that, the PDMS was inspected and all air bubbles and dust particles were removed using a plastic pipette. For the flow layer, 21g PDMS (with base to curing agent ratio of 20:1) was prepared using the same procedure. The flow wafer was placed in a spin-coater G3P (Specialty Coating Systems SCS, U.S.A.) and about 8g of PDMS was poured onto it. Likewise, all air bubbles and dust particles were removed. The PDMS was then spin coated at 1,800 r.p.m. for 1 minute. The flow mold with spin-coated PDMS was then placed on a flat surface for 50 min to allow the coating to flatten. Both PDMS coated wafers were then cured for 30 minutes at 80°C. After that, the three control layers per mold were cut and peeled off the mold and the inlets of each control line were punched (OD = 889 mm) using a precision manual-punching machine (Syneo, USA). Each control layer was then aligned by hand on top of a flow layer using a Nikon stereo microscope. The aligned devices were then baked at 80°C for 90 minutes, allowing the two layers to bond together. The assembled devices were then cut and peeled off the flow mold. The flow layer inlets were then punched. Finally, the device and a No. 1 coverslip (24x60mm, BRAND 470820) were treated in an oxygen plasma machine FEMTO Model 1A (Diener, Germany) for 20s, and then bonded together. The assembled device was then baked for 2 mins at 90°C.

##### 1.2.4 Device Operation

The MICROSTAR device contains ~2,800 microvalves, which allow fluid and cells inside the device to be precisely manipulated. Chip priming (Figure S6A) is described in more detail in section 1.3.2. The 16 cell loading inlets are used to transport liquid cultures into the 16 cell culture areas of each column. Figure S6B shows the control line setting during the cell loading process. A multiplexer [3] is used to control and select a specific cell loading inlet (Figure S5). Another control line "CL1" isolates columns during cell loading to avoid cross-contamination (Figure S6C). In the same control line, two valves are placed at the top-left and bottom-right positions of each cell culture area to direct flow into the microgrooves for cell-loading (Figure S6C). Only media is able to pass through the sieve structure at the bottom of each microgroove, effectively trapping and retaining cells since their diameter is more than 3 times larger than the sieve channel height. In each column, there are 128 possible flow paths (the number of microgrooves). The length of each path is designed to be identical in order to match flow resistance and thus flow velocities. Hence, the trapping efficiency of each microgroove should be identical in the beginning of the loading process. Once cells are beginning to be trapped, the trapped cells in a microgroove increase the fluid resistance of its path causing other microgrooves that do not yet contain cells to be loaded. Once the cell loading process of a column is completed, the next column is selected and the process repeated until all columns are loaded.

Row isolation (Figure S6D,E) and media switching (Figure S6F) are the two main operations required for on-chip cell culturing. In the starvation phase, each pair of rows is perfused with a different $P_i$ concentration. In order to maintain stable $P_i$ levels supplied by the dilution generator, it is necessary to physically isolate these rows. The control line "CC1" is designed to disconnect adjacent rows to ensure that there is no mixture/dilution of media occurring in the cell trap array during on-chip cell culture (Figure S6E). Other than that, "CC1" also disconnects the two parallel channels in the chamber, cell wash channel (upper channel), and media feeding channel (lower channel), to keep the media feeding channels clean. The three media inlets and two wash channels form a media switching unit which enables changes in $P_i$ level (Figure S6F). During on-chip cell culture, media is supplied from the media switching unit, to the cell trap arrays via the dilution generator, and eventually exits at the chip outlet "CO1". The detailed microvalve operation of cell loading and on-chip cell culturing will be discussed in detail in section 1.3.3.

##### 1.2.5 Dilution Generator Characterization

To validate $P_i$ concentrations, a tracer signal was quantitated throughout the duration of a typical experiment. Measurements were obtained at the cell-trap inlets, the cell-trap outlets, the first and the last culture chamber in each row. The media used in this experiment was the same as the media used in promoter characterisation experiments (see section 1.3.1). Furthermore, another experiment was performed to study the stability of the tracer used in by tracking the signal in the

"MI2" channel, "MI3" channel, DG output supplying row 1-2, and DG output supplying row 15-16 over time.

Figure S7A and B show the tracer signals across DG outputs over time. The data in Figure S7A and B are the mean signal of the cell-trap inlets and outlets, and the chambers, respectively. Tracer signals rose when the dilution generator generated 8 different levels after the first switch and back to the basal level after the second switch. The tracer signal in each zone was stable over time. Figure S7C shows the mean normalised signal of inlet/outlet and chambers in each DG output zone.

Figure S7D shows that the signal at "MI2" and DG output 1 were identical over time. DG output 1 followed the trend of "MI2". The same result occurred in "MI3" and DG output 8. It explains that the initial fluctuation in Figure S7A and B was due to the changes in the "MI2" source but was not a systematic change indicating that on-chip dilution was stable and robust over time.

Switching rate was also measured. The last chamber in a row was captured in the first 10 min of the two switches with 10 second time resolution. Figure S7E shows that media reached the culture areas in 40 s and steady state in 1 min.

#### 1.3 MICROSTAR Experiments

##### 1.3.1 Cell and Media Preparation

Glycerol stocks (in 96-well plates or in tubes) were taken from storage at -80°C. For 96-well plate stocks, 2$\mu$L culture from the 96-well plate stock were inoculated by a 96-pin sampling replicator VP 408FS2AS (V&P Scientific, U.S.A) and then cultured in a 96-well plate with 100 $\mu$L YPD in each well overnight at 30°C in a plate shaker (Jencons Millennium 2000, Jencons Scientific, UK) at 900 rpm. Three rounds of flame sterilisation were done using ethanol to sterilise the sampling replicator before each copying process. For freezer stocks in tubes, they were sampled with a loop and transferred into 3 mL YPD and then cultured overnight at 30°C in an incubator (GYROMAX 737R, Amerex Instruments, U.S.A) at 250 rpm. The next day, 2 $\mu$L of these cultures were inoculated into tubes containing 1 mL SC media and cultured in the incubator for 5 hours, and then loaded into the microstar device.

Typically, three types of media were prepared for the on-chip experiment, (1) $P_i$-rich SC medium, (2) $P_i$-high SC medium, and (3) $P_i$-low or $P_i$-free SC medium. Inorganic phosphate from a pre-diluted stock (8.75 $\mu$M for $P_i$-high media and 1.75 $\mu$M for $P_i$-low media) and a tracer (6.3 $\mu$g/mL sulforhodamine for promoter characterisation experiments and 5.3 $\mu$g/mL fluorescein for transcription factor localization experiments) were added to the $P_i$-high medium, whereas $P_i$-low medium received only inorganic phosphate. Media were stored at 4°C and kept in an incubator for at least 2 hours at 30°C before on-chip experiments. All aforementioned medium contained chloramphenicol at a final concentration of 34 $\mu$g/mL.

##### 1.3.2 Chip Priming

First, all control inlets were connected to Tygon tubing pre-loaded with oil (Fluorinert FC 40, CAS# 51142-49-5). The lines were then pressurised at 41 kPa to out-gas prime the on-chip control lines. Tygon tubing and therefore the on-chip control lines were regulated by computer controlled solenoid valves. After that, pressure was increased to 179 kPa and all valves were inspected visually to ensure they were closed under this pressure. The flow layer was then ready for perfusion. The flow layer was then completely primed with SC medium. A tubing connected to a bottle with $P_i$-rich SC media was plugged into the media inlet "MI1"and two intermediate $P_i$ SC media were plugged into the media inlet "MI2" and "MI3" as shown in Figure S5 and S6A. All media bottles were pressurised at 31 kPa. The media wash channels "MW1" and "MW2" were then flushed by the two intermediate $P_i$ SC media from "MI2" and "MI3", respectively, by releasing the valve "MS3" and "MS4" as shown in Figure S6A(b). Then, "MS4" was closed and "MS1" was opened to let "MI1" wash both "MW1" and "MW2". All air bubbles and dead volumes of the two $P_i$-intermediate media inputs should be cleared at this step. After that, the main body of the device was ready to be perfused. "DG1" remains closed until all air in the dilution generator was cleared. Then, "DG1", all the valves controlling the cell trap array and cell loading channels were opened to perfuse the remaining parts of the device. Visual inspection under microscope with a 4x objective was performed to ascertain that no air bubbles were present in the chip. Normally, the entire process took around 20 minutes.

##### 1.3.3 Cell Loading

Before loading cells into the cell traps, "CL7" and "CA1" were closed and opens the "MX1-8" to entirely fill all the cell loading channels "CI1-16" until a droplet was formed on each "CI" to ensure there was no air bubble in the inlets. "MX1-8" were closed again afterwards. Liquid cultures of each strain (see above) were taken from the incubator and loaded into individual Tygon tubing. Tubing with cells were then connected to the 16 cell loading inlets CI1-16. After that, the device was placed into cell loading mode by closing "CL3" and "CL1". These two valves were used to physically separate columns and create actively directed flow towards the cell traps. Typically, the cell loading process starts from the first column. Tubing connecting to "CI1' was connected to a pressure source (24.1 kPa) and then the multiplexer (MX1-8) switched to open the path for the first column (both upstream and downstream). The cell loading process was tracked manually using a low magnification objective (4x). It took ∼30 seconds to trap sufficient cells into the microgrooves. The loading process stopped once the traps were filled. The same procedure was repeated sequentially for the other columns. Nominally all 16 columns were used in promoter characterisation experiments and 5 columns were used in transcription factor localisation experiments.

##### 1.3.4 On-chip Cell Culture

After cell loading, the chip was placed into cell culture mode. "CL1" was opened and then "CC1" was closed allowing media to enter the cell trap array via the dilution generator to wash away cells not located in the culture grooves and to continuously supply the trapped cells with medium. Typically, the on-chip experiments consisted of three phases, pre-starvation, starvation and recovery. This was accomplished by controlling the media inlet control unit (Figure S6F). The duration of phase 1, 2 and 3 were 6, 16 and 16 hours, respectively. Media bottle was pressurised at 31 kPa when it was used, and unpressurised otherwise. The microscope housed in an incubator chamber (ice-Cube&Box, Life Imaging Service, Switzerland) and temperature control unit which maintained the temperature at 30°C throughout on-chip operations including chip priming, cell loading, and cell-culture.

##### 1.3.5 Time-lapse Acquisition

Time-lapse imaging was fully automated using a custom written visual basic program controlling a fully automated epi-fluorescent microscope (Nikon Ti-E, Nikon Japan). The MICROSTAR device contains 256 cell trap areas. Images of each area were acquired every 20 minutes (promoter characterisation experiments) or 30 minutes (transcription factor localization experiments). The location of each area was calculated by the software and then manually adjusted. Phase contrast and fluorescent images were acquired at every position. The hardware for imaging were (1) Objective CFI Plan Apochromat Lambda D 60X Oil (Nikon, Japan), (2) LED illumination source CoolLED pE-2 (Custom interconnected Ltd., UK), (3) shutter Lambda SC (Sutter Instrument, U.S.A.), and an EMCCD camera with a 1024x1024 pixel sensor array (Ixon DU-888, Andor Technology, UK) was used to acquire images. The camera sensor temperature was set to -70°C and cooled by a liquid-based cooling system (Oasis 160, Solid State Cooling Systems, U.S.A.) at 20°C . The microscope built-in intermediate magnification switch was set to 1.5x to obtain a 90x magnification. With this magnification, the entire cell culture chamber and a portion of upper and lower flow channel were visible. Typically, an experiment lasted 38 hours. The Nikon hardware-based perfect focus system (PFS) was used to maintain a stable focus throughout the experiment. Objective oil (Nikon type F, Nikon, Japan) was replaced and refilled every 12 hours to compensate for evaporation and property changes due to incubation at 30°C. For promoter characterisation experiments, 50 ms and 400 ms exposure times were used for phase contrast and fluorescent images, respectively. For transcription factor localization experiments, 3-layer z stacks (+0.5 $\mu$m, center, and -0.5 $\mu$m) were taken, and the exposure times for phase contrast and fluorescent images were 50 ms and 4 s, respectively. FITC filter and 470 nm light excitation was used to capture yoEGFP (promoter characterisation), whereas a TxRed filter and 535 nm light excitation was used to capture mScarlet-i (transcription factor) and red tracer signal (5 s exposure time). A GFP filter was used to capture green tracer signal (5 s exposure time).

##### 1.3.6 Calibration Image Acquisition

Darkfield and flatfield calibration images were taken right after completion of the experiment. To capture darkfield images, the light path into the camera was blocked. Then, 500 images were taken using the same camera settings and exposure times as used for taking both phase contrast and fluorescent image. To capture flatfield images, 81 positions (a 9x9 array) of a blue auto-fluorescent plastic slide (92001, Chroma, U.S.A) were sampled (both phase contrast and fluorescent channel) with the same imaging setting as used in time-lapse imaging.

##### 1.3.7 Strains analysed in MICROSTAR experiments and sample size

Table S2 shows the strains analyzed in each MICROSTAR promoter activity experiment (exp) and transcription factor localisation experiment (TF). To ensure high data quality, data filtering was performed. For exp experiments, two rounds of exclusions were done based on (1) image quality control (see section 1.4.1), and (2) exclusion of chambers/rows with defects during the experiment (e.g. channel clogging and fabrication defects). For TF experiments, sample grooves were manually screened (see section 1.4.3).

Figure S14A-C and Figure S18 A-C show the number of single cells, grooves and chambers measured in all reported MICROSTAR experiments. The number of chambers and grooves measured are the number of chambers/grooves containing cells by the time point of two hours before starvation. They were detected and registered by the automated image analysis pipeline (see section 1.4.1). For promoter characterisation experiments, the measured number of single cells is the number of cells detected and analyzed by the image analysis pipeline. Figure S15 shows the number of single cells measured across $P_i$ concentrations for each strain. For TF experiments, the number of grooves and chambers were registered manually as mentioned. As there was no cell segmentation applied to TF data, the number of cells was estimated by multiplying the number of frames, the number of grooves and 8 (typical number of cell at the bottom 4 layers).

##### 1.3.8 Dilution Generator Performance

The quality of on-chip dilution was quantified by monitoring the fluorescent tracer that was added to the media (see section 1.3.1). During time lapse imaging, tracer signals were measured at the time point of (1) 10 min before switching to starvation, (2) 10 min after switching to starvation, (3) 10 min before switching to recovery, and (4) 10 min after switching to recovery. Measurements were taken at the cell-trap inlets and cell-trap outlets. Figure S7C shows that these positions reflect the concentration level in chambers. At the end of experiment, media with tracer (originally in "MI2") was supplied to the entire chip from "MI1" to acquire a reference signal of undiluted tracer.

Figure S8 shows the mean tracer signals across 8 DG output zones (two inlets and two outlets) at the four time point of each MICROSTAR experiment. Two switches were successfully performed in each experiment. The concentration profile was maintained throughout starvation phase. The small difference between the start and the end point of starvation is explained in section 1.2.5 and Figure S7D. The on-chip diluted $P_i$ concentrations were calculated using these tracer signals given the user-defined intermediate concentrations in high and low $P_i$ media sources. Figure S9 shows the calibrated $P_i$ levels of each experiment. The mean values of the start and end point were used in this paper.

#### 1.4 MICROSTAR Image Processing and Data Analysis

##### 1.4.1 Image Processing

Processing of raw microscope imagery into summarized, per-cell, per-frame features was performed using a custom, parallelized analysis pipeline written in Python. Furthermore, the pipeline depends on both CellProfiler [5], a tool for automated image analysis, and Ilastik [6], an interactive learning and segmentation toolkit.

The pipeline consists of the following files:

1. *runAnalysis.py* - contains analysis configuration and control flow. The pipeline is initiated by executing this file.

2. *rawprocessing.py* - a module containing methods related to sorting and evaluating the raw timelapse and calibration images.

3. *preprocessing.py* - a module containing methods related to processing the raw images into calibrated, aligned, single-trap timelapses.

4. *singlecellprocessing.py* - a module containing methods related to cell region of interest (ROI) segmentation and single-cell feature extraction

5. *qualityassessment.py* - a module containing methods related to assessing the quality of the imagery and analysis of each timelapse (e.g. focus, alignment, cell growth, etc.)

Before analysis commences, each experiment is given a unique name for reference, [EXPTNAME]. The nomenclature for our experiments is "TY_[DATE]," with the date given in YYMMDD format (e.g. "TY_200320"). All data and analysis results are archived within a directory given this name. After analysis is completed, the directory structure is organized as follows:

1. *[EXPTNAME]/code* - contains a complete copy of the pipeline at the moment the analysis was completed.

2. *[EXPTNAME]/raw* - contains all raw microscopy imagery as well as four spreadsheets which contain specifications of the positions of strains, timings, media conditions, any traps that should be excluded due to defects or failures, as well as any other notes associated with the execution of the experiment.

3. *[EXPTNAME]/preprocessed* - contains analysis results relating to the production of calibrated, aligned, single-trap timelapses. The resulting ImageJ hyperstacks are contained in the 'stacks' folder. The 'calibration' folder contains the flatfield and darkfield images used to calibrate each raw image. The 'metadata' folder contains metadata associated with each microscopy image, such as the camera, filter, and illumination settings. Finally, the 'movies' folder contains a movie of each single-trap timelapse.

4. *[EXPTNAME]/single_cell* - contains analysis results relating to the segmentation and feature extraction of the single cells within the trap timelapses. The 'scData' folder contains a spreadsheet for each trap listing the measured features of each cell identified in each frame of the trap timelapse. The 'quality' folder contains the results of the quality assessment in a compact, 'pickled' Python format, which can be unpacked using the 'load_stack_quality' method within 'qualityassessment.py.' The 'plots' folder contains a variety of plots which exhibit the behavior of the cells within each trap, including cell fluorescence and area over time as well as diagnostic plots arising from the quality assessment. The 'masks,' 'overlays,' and 'multilayer_masks' folders contain the results of the cell segmentation process, with 'masks' and 'multilayer_masks' containing the regions of each trap which have been identified to contain cells and multilayer cells respectively, and 'overlays' containing diagnostic images of each brighfield image with the identified cell ROI contours overlayed. Finally, the 'single_cell' folder contains the batch processing file 'Batch_data.h5' used by CellProfiler during the segmentation process.

##### Analysis Preparation

To prepare for the analysis, the user must decide the [EXPTNAME] and generate an initial directory structure consisting of '[EXPTNAME]/raw/' that contains the 'strains,' 'conditions,' and 'manualexclusion' experiment specification spreadsheets, as well as the 'microscopy' folder which contains the raw microscopy imagery. Then, the analysis is initialized via the command 'python runAnalysis.py [EXPTNAME]' which will commence the analysis on all trap positions identified in the raw data. Optionally, the user may specify additional arguments to this command to isolate the analysis to specific traps, which can be identified via their numerical position (1-256) and alphabetic trap designation (A-H). For example, 'pos105_D' for the fourth trap among the eight traps at position 105. The user can add additional arguments as desired.

#### Pipeline Configuration

The pipeline configuration process is run once at the commencement of the analysis. During this process, the user configuration supplied within the 'runAnalysis.py' file as well as the experiment specification spreadsheets contained within the 'raw' folder are read and the run-time configuration to be used by all following steps is generated. If the microscopy images within the 'raw' folder have not yet been sorted into subfolders the user will be presented with a listing of identified timelapses, positions, and channels and can then choose whether or not the imagery should be sorted based on these characteristics. This sorting step is required to continue with the remainder of the pipeline. Finally, the complete configuration is printed to the user within the terminal and the user can evaluate and choose whether or not to proceed.

#### Raw Image Processing

The raw image processing is run once during the analysis, immediately following the user authorization to proceed. During this step, the following actions are taken:

1. Establish directory structure
2. Sort raw images
3. Extract metadata from the raw and calibration
4. Ensure that the calibration images match the exposure time and camera settings of the raw images
5. Prepare darkfield (DF) and flatfield (FF) calibration images
6. Prepare image list and batch processing file for later CellProfiler-based image segmentation
7. Prepare multiprocessing workers and initiate batch processing

The directory structure is established as previously described. Raw timelapse images are sorted according to their phase of the timelapse, position, and channel, whereas the raw calibration images are sorted according to the type of calibration (darkfield or flatfield) as well as the channel and shutter exposure settings. Metadata, such as camera and optical filter settings and measurements (e.g. exposure time and sensor temperature) for each acquired frame are extracted and stored. A verification is performed to confirm that the measured calibration image conditions match those of the acquired timelapses.

Darkfield and flatfield calibration images are computed from the raw DF and FF images acquired following the acquisition of the trap timelapses. Single DF images are highly noisy, so the mean of 500 frames is computed to establish a single DF frame. Furthermore, it was observed that there were rare outlier events in which single pixels would receive a signal far outside the typical range. Even when averaged among 500 frames, these outliers had a substantial effect on the mean of the affected pixels. In order to compensate for this behavior, before the DF mean was computed, first the median and interquartile range (IQR, i.e. 75th percentile minus the 25th percentile) were computed for each pixel in the frame. Any pixels within any frames which fell outside the range of $median/pm10 \times IQR$ were removed from the computation of the mean. The FF calibration images are calculated for each channel by first subtracting the mean DF image with the same exposure time, and then calculating the median across the 81 acquired FF images. By using the median, across a large number of images acquired at different positions of the fluorescent calibration slide, we can isolate within the FF calibration image only the nonuniformities which are present across all frames, rather than those that are simply the result of dust or defects within a single frame.

Before completion of the raw image processing step, a one-time preparation step for the later single-cell segmentation process is performed in which a CellProfiler batch processing file is generated along with an indexed spreadsheet containing a list of all acquired dia images within the timelapses.

Finally, a list of jobs to be performed by the multiprocessing workers is prepared, and distributed to the worker threads which will run them in the following steps. Each worker thread runs on one core of the analysis computer. Given our hardware configuration, we utilized 38 out of the available 40 cores for workers with the remaining two being available for any overhead operations.

#### Timelapse Processing

The timelapse processing step, which prepares the timelapse images for subsequent single-cell analysis, is executed once per each of the 256 cell trap areas. During this step, the following actions are taken:

1. Calibrate raw images using DF and FF images
2. Align subsequent frames to compensate for drift
3. Split images into eight separate cell trap images
4. Produce a single ImageJ hyperstack timelapse including all channels and z-levels for each of the eight traps
5. Generate a movie of the timelapse for each of the eight traps

Each raw timelapse image is corrected for both darkfield and flatfield artifacts which are inherent to the optical system. The DF corrects for the background counts observed on any image, even within the presence of any light incident on the sensor, whereas the FF corrects for the nonuniformity of the light intensity that arises as light travels through the various apertures of the optical path of the microscope system. Furthermore, given that the FF calibration is collected in a reproducible manner, the FF can also serve as a method for standardization of fluorescence across experiments, even in the presence of variations within the excitation intensity as well as adjustments to the optical path.

Each calibrated image is calculated as $IM_{cal} = \text{int}\left(1000 \times \frac{IM_{raw}-DF}{FF}\right)$. Thus the background DF counts are subtracted, and the resulting image is then normalized by the FF. However, because the intensity of the FF image is approximately on the same order as the DF-corrected image, the resulting floating-point image is comprised of values which are on the order of unity. Many image processing operations require that the pixel values are stored as integers, including those operations used in the downstream steps of this pipeline. Thus, we multiply this floating-point image by a fixed value which is established in the configuration of the pipeline, herein set to 1000. Following this re-scaling, the result is recast as an integer.

Following calibration, an image alignment step is performed to compensate for slight drifts over time in the position of the trap area within the frame. This alignment is performed using the images at the center of the z-stack in the brightfield channel. These registration corrections are purely translational - no rotation or other affine transforms are performed. Alignments are all performed relative to the first frame of the sequence. Each frame is aligned in a two-step process, the first being a coarse alignment and the second being a greater precision alignment.

First, the coarse alignment is performed using a binary 'background mask' of the frame, wherein the cell-containing regions are excluded and only regions corresponding to the static background portions remain. This background mask is itself calculated by evaluating the signal generated by the mean of a horizontal slice of the image containing the cell traps. This signal exhibits a periodicity with bright regions corresponding to walls and cell-containing areas, and dark regions corresponding to the background between them. Given this periodicity and the known geometry of the spacing and width of the traps, the horizontal extent of each trap can be determined. To determine the vertical positioning of the traps, the mean of vertical slices of the previously identified trap-containing horizontal regions is evaluated, and a peak corresponding to the transition from the trap to the sieve at the bottom of the trap is then determined for each of the eight traps. These eight values should be in agreement, as the traps should all be at the same vertical position within the frame. Thus, we calculate the median and IQR of the eight values, remove any that exceed a distance of two IQR, and then calculate the mean of the remainders to establish the bottom of the traps. Finally, the top of the trap is established as a fixed distance above the bottom of the trap given the known geometry of the traps. After this process we have determined coordinates for the top, bottom, left, and right sides of each cell-containing region of the frame, hence the regions outside of these areas can be considered as a background which should remain static during the timelapse. The coarse alignment can be performed simply using the position of the top-left corner of the left-most trap in the background mask.

The fine alignment follow the coarse alignment, and again uses the background mask - however for this alignment the masked brightfield image is used for alignment rather than just the mask itself. Before alignment, a histogram equalization is performed on these background regions to compensate for frame-to-frame variations using the 'createCLAHE' method of the OpenCV library. Fine alignment is then performed on the resulting images using the 'findTransformECC' method of the same library. The resulting translational warps are applied to the original frames (across all channels and z-stacks) and are also recorded within the metadata of the final timelapse.

Following the timelapse alignment, the regions containing each of the eight individual traps are isolated and the resulting single-trap timelapse stacks are recorded. All metadata is included within the stack file itself, including the raw image metadata, experiment metadata (such as strain, media conditions, and experiment timings), and preprocessing metadata (calibration file locations and alignment warps). Finally, a '.avi' movie is automatically generated for each timelapse to facilitate manual experiment evaluation using an ImageJ macro.

##### Cell Region of Interest (ROI) Segmentation

The cell ROI segmentation process, in which the brightfield images are used to determine the locations of cell objects within each frame, is performed once per each of the 256 cell trap areas. During this step, the following actions are taken:

1. Collect and join into a single images the brightfield (dia) images from the eight cell traps for each frame
2. Execute the CellProfiler/Ilastik batch processing routine to produce per-cell, per-frame ROIs
3. Separate the segmented ROI results for each of the eight traps

To prepare for segmentation, the single-trap image stacks for each of the 256 trap regions are joined back together. While the segmentation process can indeed be performed on the single-trap stacks themselves, it was identified that performance is improved by segmenting 256 larger stacks each containing eight traps versus 2048 single-trap stacks.

The brightfield images at the center of the z-stack used for segmentation. Each image passes through a CellProfiler pipeline with the following steps:

1. Histogram equalization
2. Classification of pixels into probabilities for cell, membrane, background, and multilayer classes via Ilastik
3. Determine initial cell objects based on a filled watershed using the predicted membrane image
4. Measure the size, shape, and intensity of the initial cell objects
5. Filter the initial objects to remove non-cell and multilayer objects
6. Expand the identified cell objects slightly

7. Produce and record an indexed mask of the resulting cell objects

Histogram equalization is used to minimize variations from frame-to-frame or experiment-to-experiment in the brightfield intensity. The pixel classification executed using a previously-trained classifier using the 'Autocontext' workflow within Ilastik. A set of 13 histogram-equalized images from various experiments, trap areas, and experiment timings were used for training. Training consisted of manually labeling the classes of representative pixels within these images as cell, membrane, background, or multilayer. The resulting classifier can then process novel images and generate the probabilities of for each pixel to be within each of the four classes.

Following the pixel classification, a watershed operation is performed using the predicted membrane image to identify an initial set of objects. This set of objects contains the cell objects, but also contains objects which contain no cells or, occasionally, regions in which the cells have grown in a manner in which the trap is no longer maintaining a monolayer of cells. These erroneous objects are identified and filtered using several criteria as follows using the CellProfiler 'FilterObjects' module:

1. LowerQuartileIntensity of the predicted cell minus the predicted background image within the object must be greater than 0.0
2. LowerQuartileIntensity of the predicted cell minus the predicted multilayer image within the object must be greater than 0.0
3. Eccentricty of the object is less than 1.0
4. Solidity of the object is greater than 0.8
5. Compactness of the object is less than 1.5
6. Area of the object is between 100 and 10000 pixels
7. FormFactor of the object is greater than 0.6

The resulting cell objects are then expanded by two pixels to better represent the cell areas. This was necessary as a result of conservative classification of the membrane regions between cell areas which substantially improved the earlier initial object detection. These final cell objects are then indexed and recorded, along with a diagnostic image of the identified contours overlayed upon the initial brightfield image. Finally, the resulting cell ROIs for the full eight-trap image are separated into results for their constituent individual traps.

##### Cell ROI Feature Extraction

During ROI feature extraction, the characteristics of all cell ROIs are summarized for all cells in all frames. This process is run once per each of the 2048 trap timelapses.

For each cell ROI, the following statistics are measured and recorded:

1. Frame index
2. ROI index
3. Area
4. Position of the center of the ROI
5. Fluorescence mean
6. Fluorescence background
7. Fluorescence mean minus background

The fluorescence background is calculated by measuring the 5th percentile of pixel intensities within a annulus around the ROI. The ring mask is produced by the difference of a slightly dilated ROI and the original ROI.

The results of the feature extraction for each trap are recorded in a .csv. In addition to the individual ROI measurements, this file also contains the timing and media condition data corresponding to the observed frames of the trap. The file name also contains the position index of the trap area as well as the strain identifier for the contained strain. In this manner, the resulting .csv files are self-contained for downstream analysis.

##### Timelapse Quality Assessment and Plotting

During the automated quality assessment and plotting process, the timelapses are evaluated and scored on a variety of quality metrics. Additionally, diagnostics plots are created for each timelapse. This process is run ones per each of the 2048 trap timelapses.

During this process the following steps are performed:

1. Calculate focus quality
2. Calculate timelapse alignment quality
3. Generate a motion map (y-vs-t correlation kymograph of the dia channel)
4. Calculate a score to assess lack of motion (i.e. growth quality) using the motion map
5. Measure cell counts within the different regions of the trap
6. Calculate epi map (y-vs-t kymograph of the epi channel)
7. Measure the extent of any multilayer growth within the trap
8. Generate quality assessment plots
9. Compare quality scores to pre-determined quality criteria and annotate the quality performance of each trap

Focus quality is scored by evaluating the spatial sharpness of the image. To do this, the image is first normalized using the 1st and 99th percentile values in order to remove any disruptive outliers. Then, the Laplacian of the image is calculated, which produces large values in areas with high spatial gradients (i.e. sharp regions). Finally, the focus score is calculated as the difference between the 99th and 1st percentile of the Laplacian image. High scores correspond to a large difference between the tails of the distribution of the Laplacian image, meaning that there were regions of high sharpness within the image, and conversely, low values correspond to images that are blurred.

The alignment quality of the timelapse is evaluated via two scores - one which compares the correlation of each frame to it's previous frame, and another which compares to the initial frame. These 2d correlation coefficients are calculated only for background regions of the frame in order to avoid regions with frame-to-frame motion due to cell growth. Four background regions are used, corresponding to the areas above, below, to the left, and to the right of the cell trap. For each set of two frames being compared, the highest of the four region's correlation coefficients is used as the score in order to avoid possible disturbances due to cells passing through the region.

A y-vs-t correlation kymograph of the dia channel, herein termed a 'motion map,' is calculated for each timelapse. This produces a single image that facilitates evaluation of the motion of cells within the trap throughout the timelapse. To produce this image, first a mask is used so that only the interior of the trap is evaluated. Then the Pearson correlation coefficient is calculated on pixels corresponding to each horizontal slice of the trap between each frame and it's subsequent. When the correlation is high, there is a strong correspondence between the pixels in one frame to the next for this particular horizontal slice. Performing this calculation across all horizontal slices for a set of two frames produces a column of pixels which summarizes the similarity between the slices for all slices. Subsequent columns can be calculated for subsequent frames, and these columns can be assembled into a y-vs-t image. In the motion map, regions of high correlation imply that no motion is occurring between frames within a portion of the trap. On the other hand, regions of low correlation mean that substantial differences are present from one frame to the next, implying motion within that part of the trap.

The motion map can be used to produce summary statistics of the motion within the trap. A 'no-motion score' is calculated by simply calculating the proportion of the trap which exceeds a threshold correlation value of 0.8 for each frame. This per-frame score is then smoothed with using a window of five frames.

A count of cells identified within various regions of the trap is performed. These regions include the trap itself, the sieve below the trap, the area above the trap, and the area below the trap.

A y-vs-t kymograph of the epi channel is calculated, termed the 'epi map'. This follows a simplified process compared to the motion map. For the epi map, for each frame the mean is calculated over the x-axis, producing a single column of values. These columns are then assembled into a y-vs-t image. The epi map facilitates evaluation of the fluorescence both over time and as a function of vertical position within the groove by expressing this information within a single image.

The extent of any multilayer growth within the trap is evaluated using the multilayer results from the Ilastik pixel classification process which was evaluated during image segmentation. The score is simply the area proportion of the trap which was classified as multilayer.

Next, a variety of plots are generated in order to summarize these quality characteristics as well as the cell behavior measured within each timelapse. This includes density plots of the distributions of the measured cell fluorescence statistics, cell area, and a summary of the noise characteristics of the population over time. The assessed quality scores are also summarized to facilitate quality assessment.

Finally, the automatically-generated quality scores can be compared with pre-determined exclusion and warning criteria, and a summary of the pass-fail results per-trap is recorded. By incorporating these automated results alongside direct evaluation, we performed a manual screening the trap timelapses.

##### Image Analysis Computer Hardware

Image analysis was performed on a custom-built workstation with dual ten-core Intel Xeon E5-2680 v2 CPUs, a total of 128 GB of RAM, and a 2 TB SSD for storage of the active dataset. With this CPU configuration, hyperthreading enables the utilization of up to 40 logical cores. The image analysis pipeline was run with 38 worker threads, with two cores remaining for system overhead. This allows for approximately 3 GB of RAM per thread. Typical MICROSTAR experiments will have around 120 dual-channel frames captured per timelapse at each of the 256 positions. Given our hardware configuration, these approximately 30,000 frames can be fully analysed within 24 hours.

##### 1.4.2 Promoter Characterisation

The single cell data generated by the image processing pipeline was used to characterise the promoters of the 23 strains. Only the bottom four layers of cells were analysed. In Figure 2A to H, single cell data of each strain were binned according to time and concentration level. The time bin size was 1 hour. Concentrations were divided into 15 bins and the concentration level shown in the label is the mean of the interval. The line plot shows the median of each concentration bin with moving average (window size = 3). The same method applied to S12. The heatmaps in Figure 2A, C, E and G show the distribution of single cell values in each concentration bin. 50 equally sized GFP signal bins were used given the maximum and minimum value.

In Figure 2J and K, the heatmaps show the distributions of single cell data in each concentration bin for each strain. Only steady state data (10 to 15 hours of starvation) was used for this analysis. The concentration bin size is 7 $\mu$M, and 50 equally sized GFP signal bins were used given the maximum and minimum value on the y axis. For the line plot, it shows the median of each concentration bin with moving average (window size = 3). Concentrations were divided into 20 bins. The markers on the line plot is the median of corresponding bin range. The same method applied to Figure S13.

Figure S16 summarises normalised promoter activity across $P_i$ levels and plots it as a heatmap.

It is divided into four programs as shown in Figure 2I. An unsupervised hierarchical clustering was applied to the data shown in the heatmap and cluster 2 groups of promoters, named decoupled and plateau. The clustering was done by a clustering function AgglomerativeClustering in a machine learning package (scikit-learn) written in Python. The metric used to compute the distance is euclidean.

##### 1.4.3 Pho4p Nuclear Signal Analysis

To measure nuclear Pho4p signal in intermediate $P_i$ conditions proved to be challenging due the overall low abundance of Pho4p and the limited degree to which Pho4p localized to the nucleus in intermediate $P_i$ conditions. We developed a simple, effective, and intuitive method that merely quantifies pixel intensity changes in image histograms and used a simple cut-off value to classify cytoplasmic and nuclear signal. Although this method is straightforward, it required a clean dataset as the method is sensitive to the occurrence of dead or dying cells, which become brightly fluorescent. Therefore, a quality control screening step was performed. All time-lapse videos were manually annotated and only perfect microgrooves were included in the analysis. The screening criteria were: (1) the microgroove maintained good growth, (2) no dead cells formed within the ROI, and (3) no dying cells or unhealthy cells were present in the ROI. The above criteria were applied to a time window ranging from 3 hours before starvation to 3 hours before the end of experiment. Videos were cropped to 115x115 pixels (ROI) which covered the bottom ~4 cell layers (approximately 8 cells per frame) and a portion of background. Videos were binned into 10 $P_i$ concentrations (1 in full, 3 in plateau, 4 in bi-stable, and 2 in off state). 8 of the 10 concentration bins have 2-3 chip replicates and all of them have chamber and groove replicates. In total, 230 microgrooves were analyzed. In the analysis, the maximum intensity of each pixel position in the z stack was used in the analysis. Any pixel intensity higher than 3000 was filtered. Figure S19 shows the histograms of Pho4p signal across the 10 concentration bins. To plot Pho4p nuclear signal as a function of $P_i$ concentration, images from 2 hours into starvation to 2 hours before the end of starvation were analyzed. Any pixel intensity higher than 850 was defined as nuclear signal and below 850 considered to be cytoplasmic. The nuclear signal of each concentration bin is the mean nuclear signal of microgrooves in each bin (sum of nuclear signal divided by no. of sample grooves in the bin). The result is reported in Figure 2F with a moving average (window size =2).

#### 1.5 Micro-Chemostat-Array (MCA)

##### 1.5.1 Device Design

The microchemostat array (MCA) device used in this study is based on a previous work [7] and includes design modifications to adapt to the requirements of this study. The device design is shown in Figure S3. The MCA contains 1,152 cell culture chambers (48 columns x 24 rows) which support on-chip cell culture with continuous media supply. Unlike the MICROSTAR chip, cells are deposited directly onto the coverslip by live-cell spotting using a microarrayer (Arrayit, U.S.A.). Cell-arraying greatly increases the number of strains that can be analyzed on a single chip. The original MCA was able to measure 384 strains under three different media conditions. In this study, we required higher throughput in terms of conditions (multiple $P_i$ concentrations). We therefore divided the MCA into 12 zones and each zone was connected to an outlet of a dilution generator to measure 96 strains under 12 $P_i$ concentrations (normally 10 intermediate $P_i$ levels, one 0 $P_i$, and one $P_i$-rich).

We also modified the cell culture chamber design. The original chamber design is a wide compartment which allows cells to grow freely in any direction. To better define and constrict the direction in which cells will be pushed due to growth we included two vertical walls in the imaging ROI in each chamber (see Figure S4A). These walls guide the cells and minimize lateral movement.

##### 1.5.2 Mold Fabrication

The mold fabrication workflow is identical to the procedure described above for the MICROSTAR device. However, different types of SU-8 (GM series, Gersteltec, Switzerland) were used and therefore the fabrication parameters differed (see Table S12, S13 and S14). The major difference is that these types of photoresist require ramp heating and cooling in baking steps instead of a step temperature.

##### 1.5.3 Device Fabrication

The chip fabrication method is identical to the procedure described above for the MICROSTAR device.However, the device was bonded onto epoxy coated coverslip instead of plasma bonding.

##### 1.5.4 Cell Culturing

The glycerol stocks stored at -80°C were used to inoculate a 96-well plate with 100 $\mu$L YPD in each well. The plates were then placed in a plate shaker for two days at 30°C. This long incubation period rendered the cells more robust to physical arraying.

##### 1.5.5 Cell Arraying

We use a commercial microarrayer (Qarray2, Genetix, UK) equipped with 4 arraying pins (SMP3, ArrayIt Corp, Sunnyvale, CA) to transfer cells from the 96-well plate onto an epoxy coated coverslip. The duration of spotting was around 50 min. The spotting pins were cleaned and dried after each spot deposition using water and compressed air. The humidity in the spotting chamber was maintained at 75%. After spotting, the MCA device was aligned to the spotted coverslip under a stereo microscope.

#### 1.6 MCA Experiments

##### 1.6.1 Chip Priming

All control lines were connected to tubing pre-loaded with oil (Fluorinert FC 40, CAS# 51142-49-5) and pressurised at 41.3 kPa to out-gas prime the control lines. Pressure was then increased to 151 kPa and each on-chip microvalve was inspected to ensure that they were fully closed. The above procedure does not apply to the pressure element located in the roof of each microchemostat. This element was primed before device-to-coverslip alignment. If this element was primed with cells underneath, the pressure caused cells to enter the sieve channel or break through the chamber outlet valve. The button valve remained un-pressurised until just before 90x time lapse imaging started.

MCA has the same media switch unit as MICROSTAR. The priming step is also the same as shown in Figure S6A. After that, media was allowed to enter the dilution generator and prime it while valve "DG1" remained closed. Then, "DG1" was opened allowing media to fill the upper flow channels. Upper "CP2" and lower channel control "CP1" remained closed to direct flow towards the cell culture chambers in order to ensure that cells remained in the chamber and avoid cross-contamination (Figure S4D(a) and (b)). After that, the pressure of chamber outlet valve "CC1" was gradually decreased from 151 kPa to 20.6 kPa. Media then started to perfuse the cell culture chamber and cells (Figure S4D(c)). Once the cell culture chamber was filled, media would enter the sieve channels and then enter the lower flow channel. At that moment, the lower flow channel control "CP1" was opened to prime the lower flow channel and the chamber outlet valve was closed to prevent cell exit (Figure S4D(d)). The chip priming procedure was completed when all air bubbles were successfully removed by out-gas priming.

##### 1.6.2 On-chip Cell Culture

The next step is initial growth of the cell spot to fill the cell culture chamber. Normally, live cell spots take 15-20 hours to recover from spotting. During this period, time lapse imaging was performed with a 4x objective. These images are used for manual quality annotation. Once the cells filled up the chambers, the chamber outlet valve was opened to allow excessive cells to exit via the upper flow channel, and the button valve was pressurised at 13.7 kPa to keep cells growing in a mono-layer (Figure S4E(a) and (b)). Similar to MICROSTAR experiments, a typical experiment consists of three phases, (1) pre-starvation, (2) starvation, and (3) recovery (Figure S4E(b), (c) and (d)). The duration of these phases were 6, 16, and 24 hours. The media in negative control zone remained the same throughout the experiment.

##### 1.6.3 Time-lapse Acquisition

The hardware, software and imaging parameters/conditions are identical to the procedure described above for the MICROSTAR device.

#### 1.7 MCA Image Processing and Data Analysis

##### 1.7.1 Image Processing

The image analysis process for the MCA experiments largely follows the procedure described above for the MICROSTAR devices.

During the image alignment portion of the preprocessing phase, the background identification process cannot be performed in the same manner, as the MICROSTAR approach relies on identifying the periodic spacing of the cell traps. Instead, for the MCA images, the background region determination is performed by first calculating an adaptive binary threshold on the dia image followed by identification of the large region of background along the bottom of the frame and along a vertical column in the middle of the frame between the two chambers. The performance of this mask generation is susceptible to disruption by cells which have made their way into the channel below the chambers, as well as occasional dust or debris which makes identification of the central column difficult. Thus, if the mask was not able to be identified as desired (that is, the central column was identified with a width of less than 25 pixels), then the background mask will be recalculated with a new bias term included on the binary threshold. This process will repeat with new bias values until either a successful mask is generated or until a maximum number of bias values have been attempted, in which case the previous frame's warp will be used and a warning will be logged. In the rare event that the background extraction fails on the reference frame (i.e. first frame), then the alignment of the of chambers in that position will fail to occur.

During segmentation, a new Ilastik autoclassification pixel classifier was trained on and for MCA images.

The full suite of quality assessment analyses were not adapted for MCA image analysis. Only plots summarizing the population fluorescence and area characteristics were generated.

#### 1.8 Transport Models

The carrier model was as previously described [8]:

$$v = e_t \frac{\frac{k_{cat}}{K_M}(s_e - s_i)}{1 + \frac{s_e}{K_M} + \frac{s_i}{K_M} + \alpha \frac{s_e s_i}{K_M^2}} \tag{1.1}$$

with

$$k_{cat} = \frac{k_2 k_4}{k_2 + k_4} \tag{1.2}$$

$$K_M = 2K_D \frac{k_4}{k_2 + k_4} \tag{1.3}$$

$$\alpha = 4 \frac{k_2 k_4}{(k_2 + k_4)^2} \tag{1.4}$$

Solving for $s_i$:

$$s_i = \frac{s_e - \frac{v K_M}{e_t k_{cat}} - \frac{v s_e}{e_t k_{cat}}}{1 + \frac{v}{e_t k_{cat}} + \alpha \frac{v s_e}{e_t k_{cat} K_M}} \tag{1.5}$$

The Michaelis-Menten model was also as described previously [8]:

$$v_{MM} = e_t \frac{\frac{k_{cat}}{K_M}(s_e - s_i)}{1 + \frac{s_e}{K_M} + \frac{s_i}{K_M}} \tag{1.6}$$

and solving for $s_i$:

$$s_{i_{MM}} = \frac{\frac{v}{e_t} + \frac{v s_e}{e_t K_M} - \frac{k_{cat} s_e}{K_M}}{\frac{-k_{cat}}{K_M} + \frac{-v}{e_t K_M}} \tag{1.7}$$

We used the following parameters for all plots: $k_2 = 4$, $k_4 = 4$ and otherwise only adjusted $K_D$ or $e_t$ values as indicated in the figure captions.

### Chapter 2

### Supplementary Figures

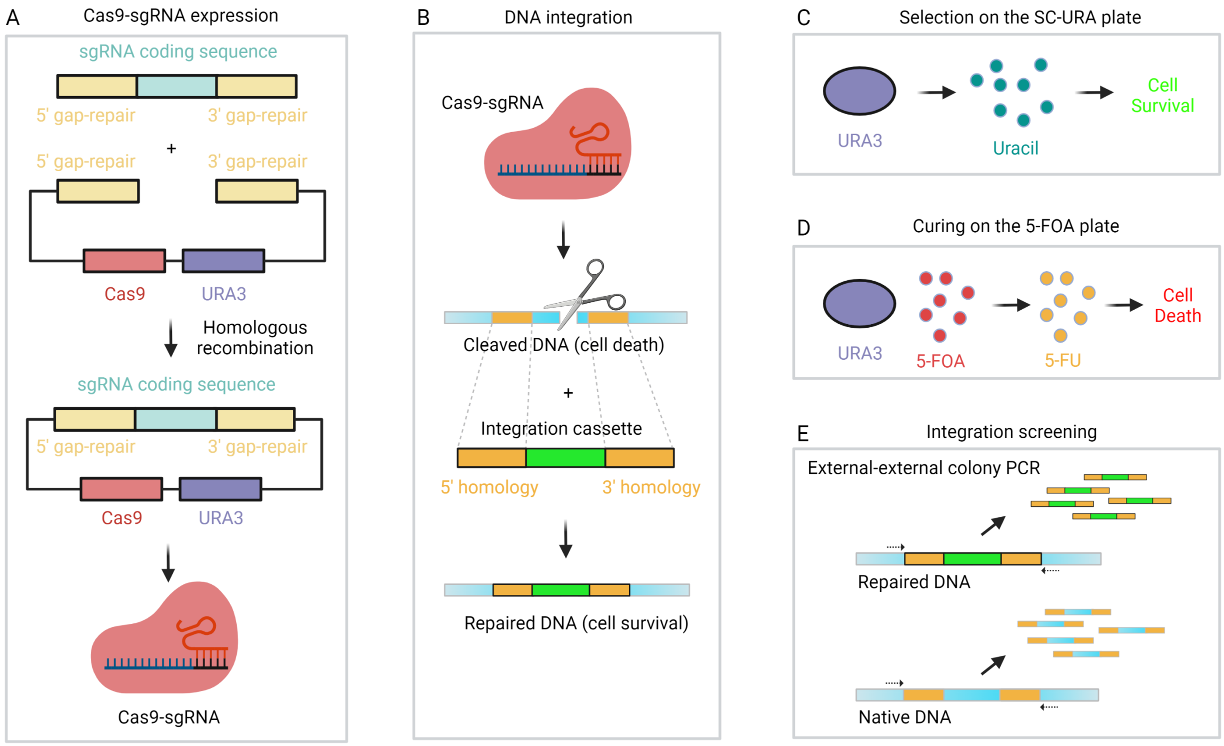

Figure S1: **Cas9-assisted cloning by homologous recombination.** **A,** The sgRNA coding sequence cassette and Cas9-*URA3* cassette were digested with EcoRV and BsmBI, respectively, for yeast transformation. After transformation, they combined to form a plasmid and expressed the Cas9-sgRNA complex to target the integration loci. **B,** The DNA insertion cassette was digested with NotI for yeast transformation. The native genomic DNA was cleaved by the Cas9-sgRNA complex to create a double stranded break, which could lead to cell death. The integration cassette was inserted into the target loci via Cas9-assisted homologous recombination and the genomic DNA was repaired. **C,** The resultant transformants were selected on SC-URA plates. Yeast cells containing Cas9-*URA3* plasmid could produce uracil and thus survive on the SC-URA plate. **D,** The colonies on the SC-URA plates were cured on 5-FOA plates to remove the Cas9-*URA3* plasmid under selection pressure. 5-FOA can be converted into 5-FU when URA3 is expressed leading to cell death. **E,** Colonies on the 5-FOA plates were screened by external-external colony PCR and the PCR products were verified by Sanger sequencing.

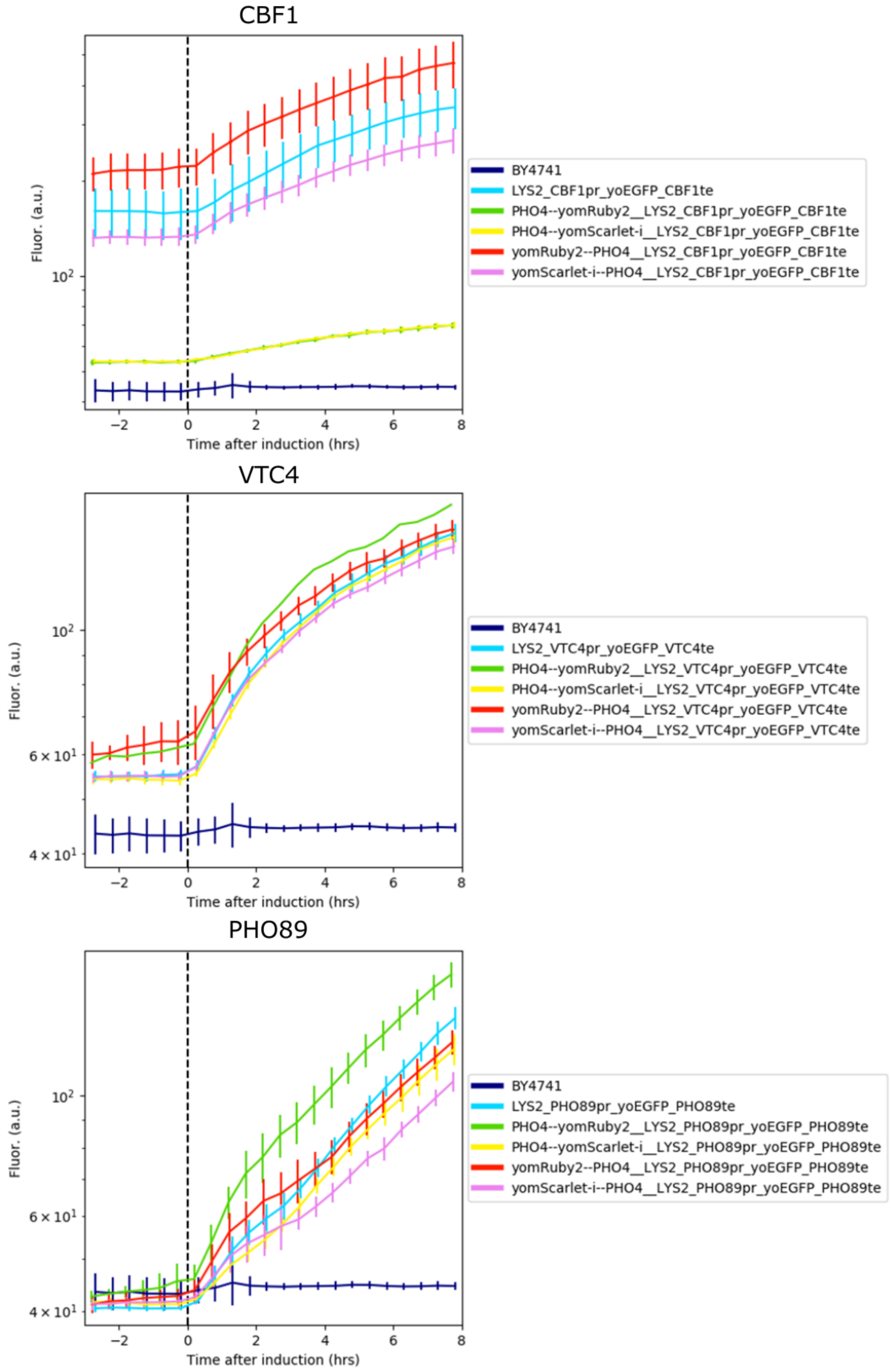

Figure S2: **Assessment of fluorescently tagged Pho4p variants.** Different fluorescent Pho4p fusion variants were tested for their ability to activate 3 target promoters: *CBF1*, *VTC4*, and *PHO89* and compared to wild type Pho4p.

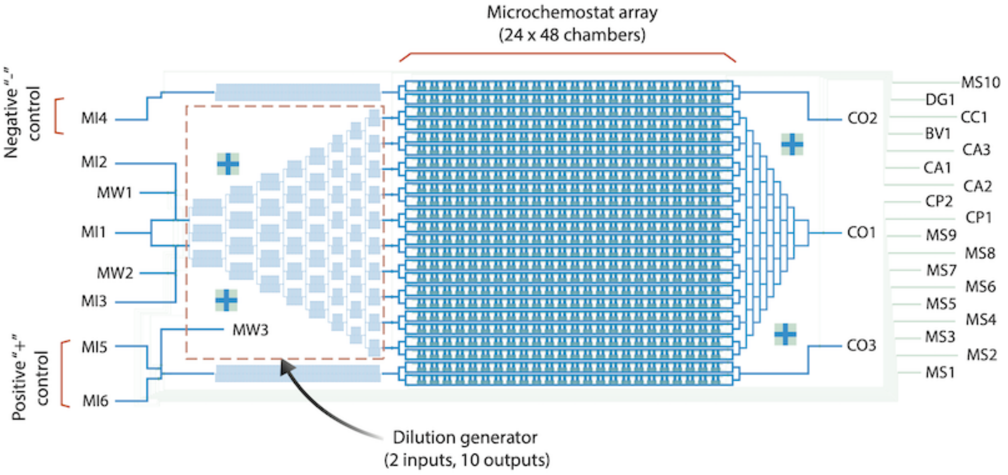

Flow layer

| | |
|---|---|
| MI1 | A media inlet of DG and typically feeding Pi-rich media. |
| MI2 | A media inlet of DG and typically feeding Pi-high media. |
| MI3 | A media inlet of DG and typically feeding Pi-low/Pi-free media. |
| MW1 | A media outlet for "MI1" or "MI2" and is used during media switch to clear dead volume and air bubble. |
| MW2 | A media outlet for "MI1" or "MI3" and is used during media switch to clear dead volume and air bubble. |
| MI4 | A media inlet of "-" control and typically feeding Pi-rich media. |
| MI5 | A media inlet of "+" control and typically feeding Pi-free media. |
| MI6 | A media inlet of "+" control and typically feeding Pi-rich media. |
| MW3 | A media outlet for "MI5" or "MI6" and is used during media switch to clear dead volume and air bubble. |
| CO1 | The exit for cells and used media during on-chip cell culture in DG zone. |
| CO2 | The exit for cells and used media during on-chip cell culture in "-" zone. |
| CO3 | The exit for cells and used media during on-chip cell culture in "+" zone. |

Control layer

| | |
|---|---|
| MS1 | Control "MI1" and is typically used in combination with "MS 1-5" during media switch. |
| MS2 | Control the inlet of DG zone and is typically used in combination with "MS 1-5" during media switch. |
| MS3 | Control all "MW" and is typically used in combination with "MS 1-5" during media switch. |
| MS4 | Control "MI2" and "MI3", and is typically used in combination with "MS 1-5" during media switch. |
| MS5 | Control "MI2", "MI3" and all "MW", and is typically used in combination with "MS 1-5" during media switch. |
| MS6 | Control "MW3" and is typically used in combination with "MS 6-9" during media switch. |
| MS7 | Control "MI5" and is typically used in combination with "MS 6-9" during media switch. |
| MS8 | Control "MI6" and is typically used in combination with "MS 6-9" during media switch. |
| MS9 | Control the inlet of "+" control and is typically used in combination with "MS 6-9" during media switch. |
| MS10 | Control "MI4" and hence the inlet of "-" control. |
| CP1 | Control the lower flow channel in the chamber array. |
| CP2 | Control the upper flow channel in the chamber array. |
| DG1 | Control the channels connecting DG and MCA, and typically closed to clear air bubbles in the DG during chip priming. |
| CA1 | Control the media and cell exit (CO1), and typically closed to clear air bubbles in the cell trap array during chip priming. |
| CA2 | Control the media and cell exit (CO2), and typically closed to clear air bubbles in the cell trap array during chip priming. |
| CA3 | Control the media and cell exit (CO3), and typically closed to clear air bubbles in the cell trap array during chip priming. |
| BV1 | Gently press the cell culture chamber to guide cell grow in monolayer. |
| CC1 | Control the outlet of chambers to prevent cross-contamination during chip priming and initial growth. |

Figure S3: **Microchemostat array (MCA) chip design.** The MCA is a two layer PDMS device. The blue layer is the flow layer, and the green layer is the control layer. It contains a 24x48 cell culture chamber array. A two-input to 10-output microfluidic dilution generator is integrated into the array. The function of the microvalves, inlets, and outlets are listed.

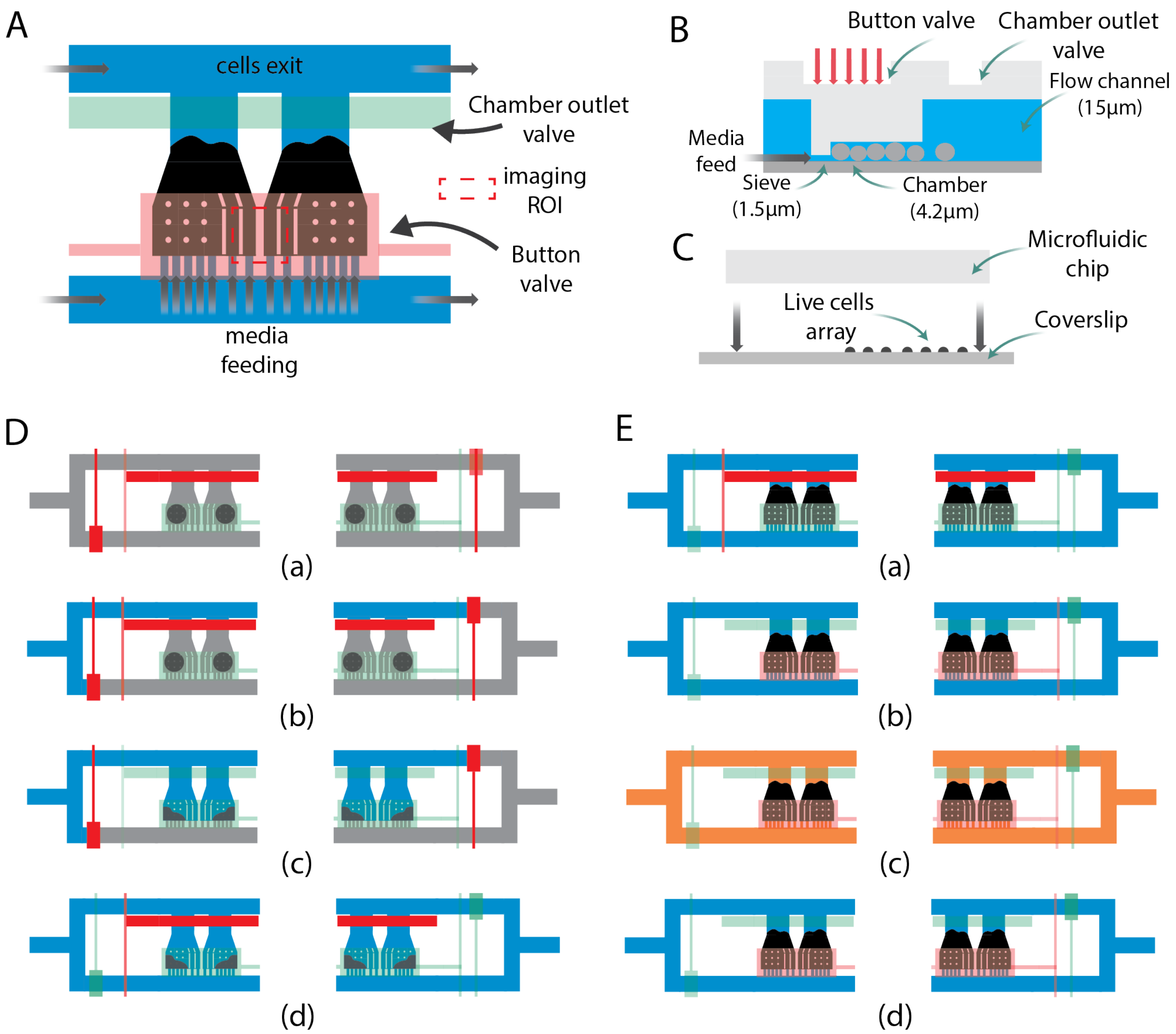

Figure S4: **MCA chamber design and experimental workflow. A.** The MCA chamber design. **B.** The cross-sectional view of an MCA chamber. The red arrows indicate pneumatic pressure. **C.** The alignment process of the MCA chip and coverslip with live cell array. **D.** MCA chip priming process and microavalve settings. **E.** MCA initial growth and media switch process. The outlet valve is closed during initial growth and opened during long-term cell culture. The button valve gently presses the chambers during long-term cell culture.

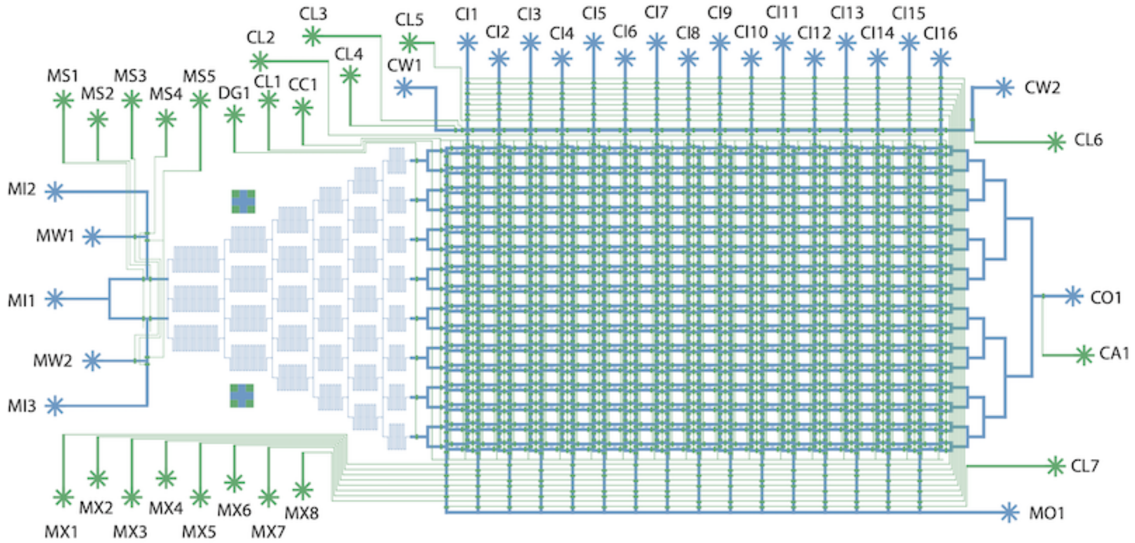

**Flow layer**

| | |
|---|---|
| MI1 | A media inlet and typically feeding Pi-rich media. |
| MI2 | A media inlet and typically feeding Pi-high media. |
| MI3 | A media inlet and typically feeding Pi-low/Pi-free media. |
| MW1 | A media outlet and is used during media switch to clear dead volume and air bubble. |
| MW2 | A media outlet and is used during media switch to clear dead volume and air bubble. |
| CO1 | An exit for cells and used media during on-chip cell culture. |
| MO1 | An exit for used media during cell loading process. |
| CI1-16 | 16 cell loading inlets. |
| CW1 | A media inlet/outlet for optional washing operation |
| CW2 | A media inlet/outlet for optional washing operation |

**Control layer**

| | |
|---|---|
| MS1 | Control "MI1" and is typically used in combination with other "MS" during media switch. |
| MS2 | Control all "MI" and is typically used in combination with other "MS" during media switch. |
| MS3 | Control all "MW" and is typically used in combination with other "MS" during media switch. |
| MS4 | Control "MI2" and "MI3", and is typically used in combination with other "MS" during media switch. |
| MS5 | Control "MI2", "MI3" and all "MW", and is typically used in combination with other "MS" during media switch. |
| DG1 | Control the accessibility of media into the cell trap array and typically closed to clear air bubbles in the DG during chip priming. |
| CL1 | Disconnect the columns of cell trap array and is typically used during cell loading. |
| CL2 | Disconnect the cell trap array and "CI1-16", and is typically used during cell loading. |
| CC1 | Disconnect the rows of cell trap array and is typically used during on-chip cell culture. |
| CL3 | Disconnect "CI1-16" and is typically used during cell loading. |
| CL4 | Control "CW1" and is typically used during media switch. |
| CL5 | Control "CI1-16" and is typically used during cell loading. |
| CL6 | Control "CW2" and is typically used during media switch. |
| CA1 | Control the media and cell exit (CO1), and typically closed to clear air bubbles in the cell trap array during chip priming. |
| CL7 | Disconnect the cell trap array and "MO1", and is typically used during chip priming. |
| MX1-8 | Control the accessibility of 16 cell loading paths using a combinatorial operation. |

Figure S5: **MICROSTAR chip design.** The MICROSTAR chip consists of two PDMS layers, control layer (in green) and flow layer (in blue). Each control and flow inlet is named and its function is indicated.

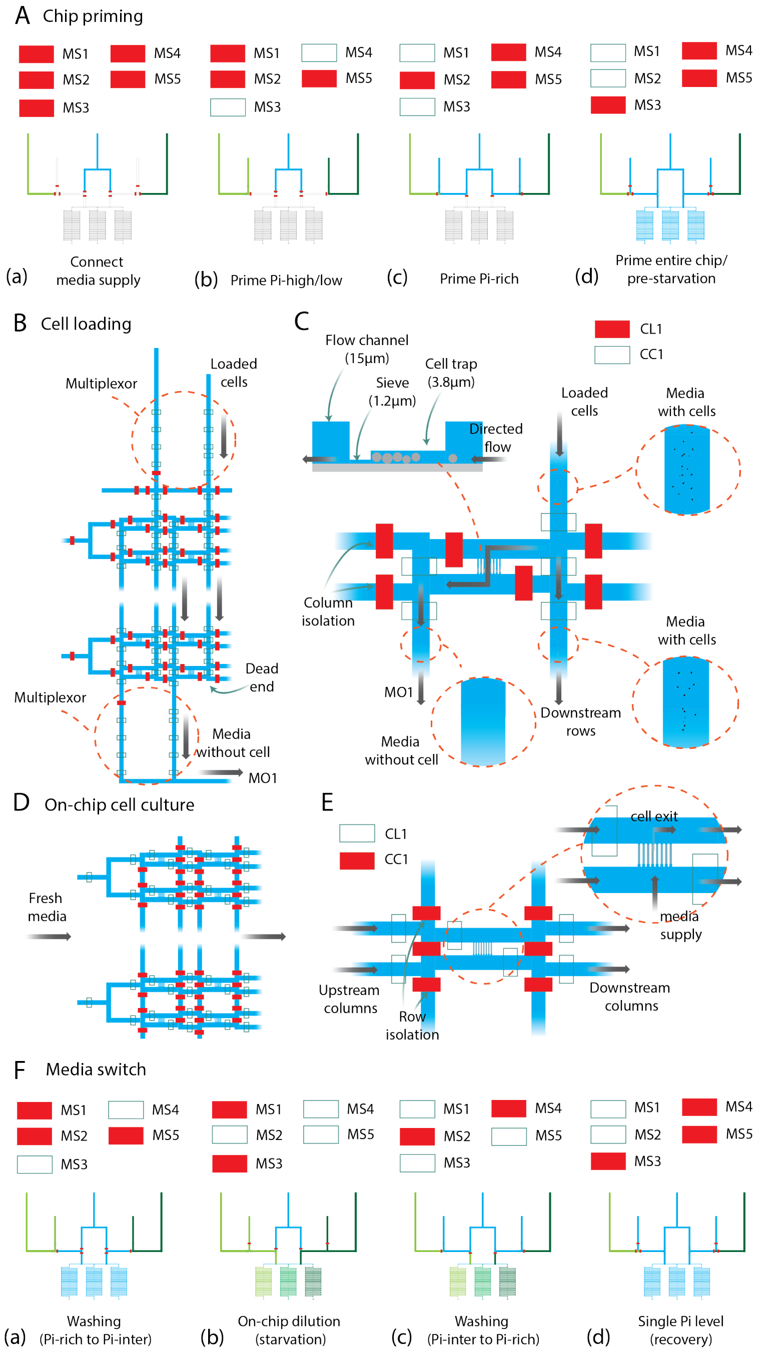

Figure S6: **MICROSTAR experimental workflow. A.** Microvalve settings for the chip priming process. These settings apply to both the MCA and MICROSTAR. **B.** Microvalve settings for the cell loading process. **C.** A close-up schematic of a MICROSTAR chamber during the cell loading process. A cross-sectional view is also shown. **D.** Microvalve settings for the cell culture process. **E.** A close-up schematic of a MICROSTAR chamber during the cell culture process. **F.** Microvalve settings for media switching (applies to both the MCA and MICROSTAR devices).

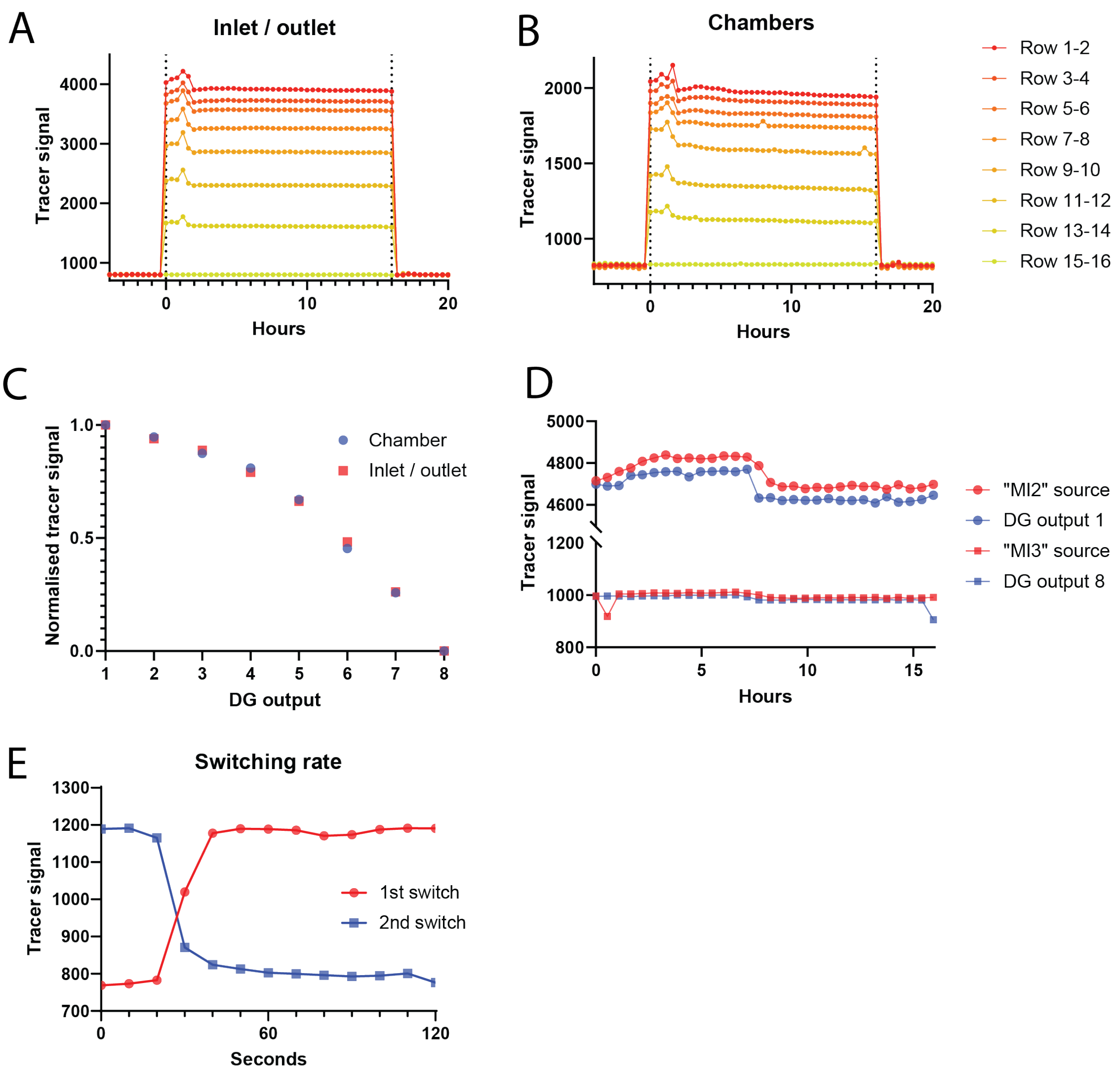

Figure S7: **Dilution generation validation and characterisation. A.** Tracer signals measured at the inlets and outlets of the eight media zones in the MICROSTAR array over time. **B.** Tracer signals measured at the chambers in the eight media zones in the MICROSTAR array over time. **C.** Normalised mean tracer signal of the inlets and outlets and chamber measured at the eight media zones. **D.** Tracer signals measured at the two media inlets for the dilution generator, the inlets of the first media zone (DG output1), and the inlets of the eighth media zone (DG output8). **E.** Media switching rate measured in a MICROSTAR chamber. The red line indicates the switch for "starvation", and the blue line indicate the switch for "recovery".

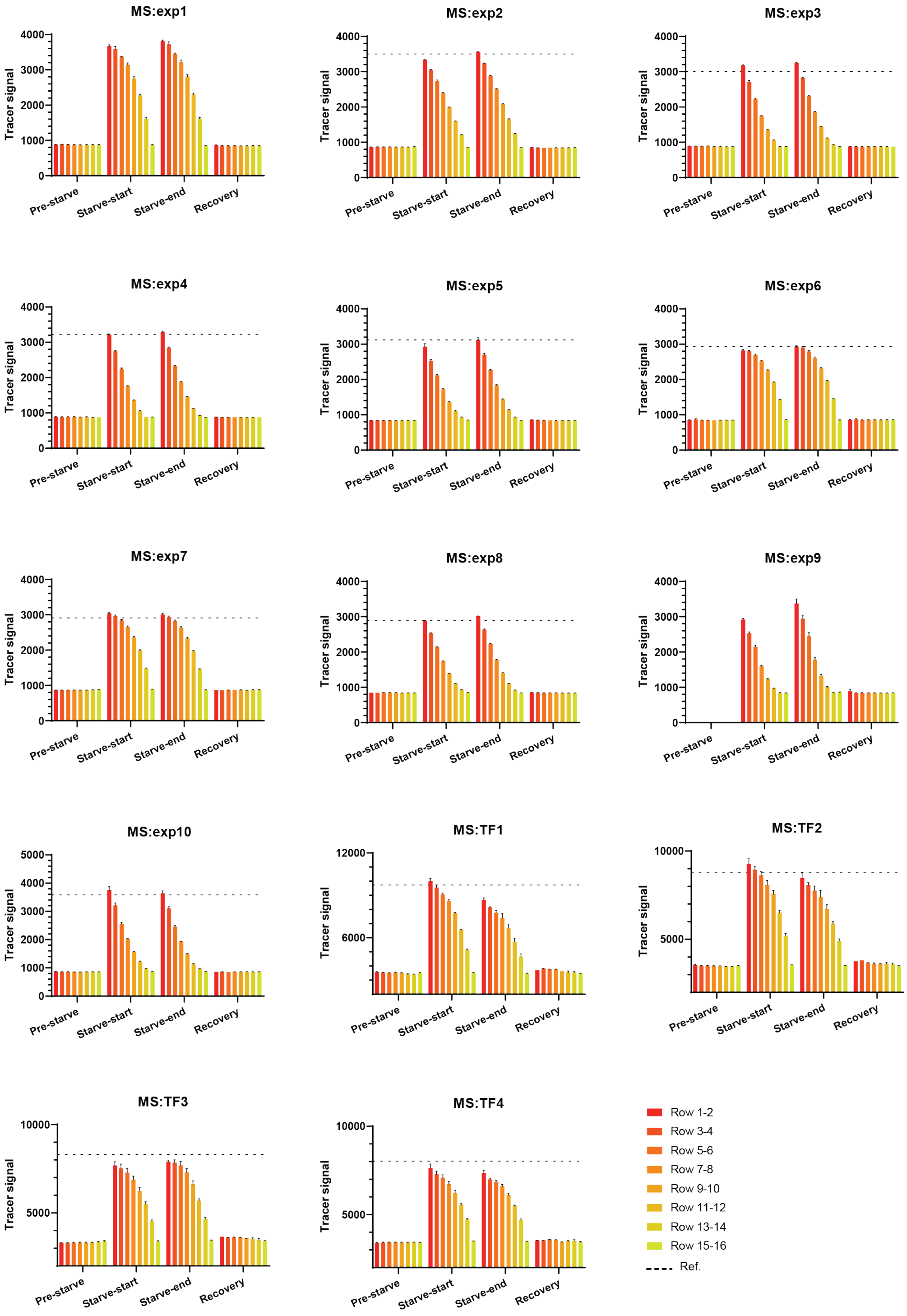

Figure S8: **Tracer signals across rows at four time points in MICROSTAR experiments.**

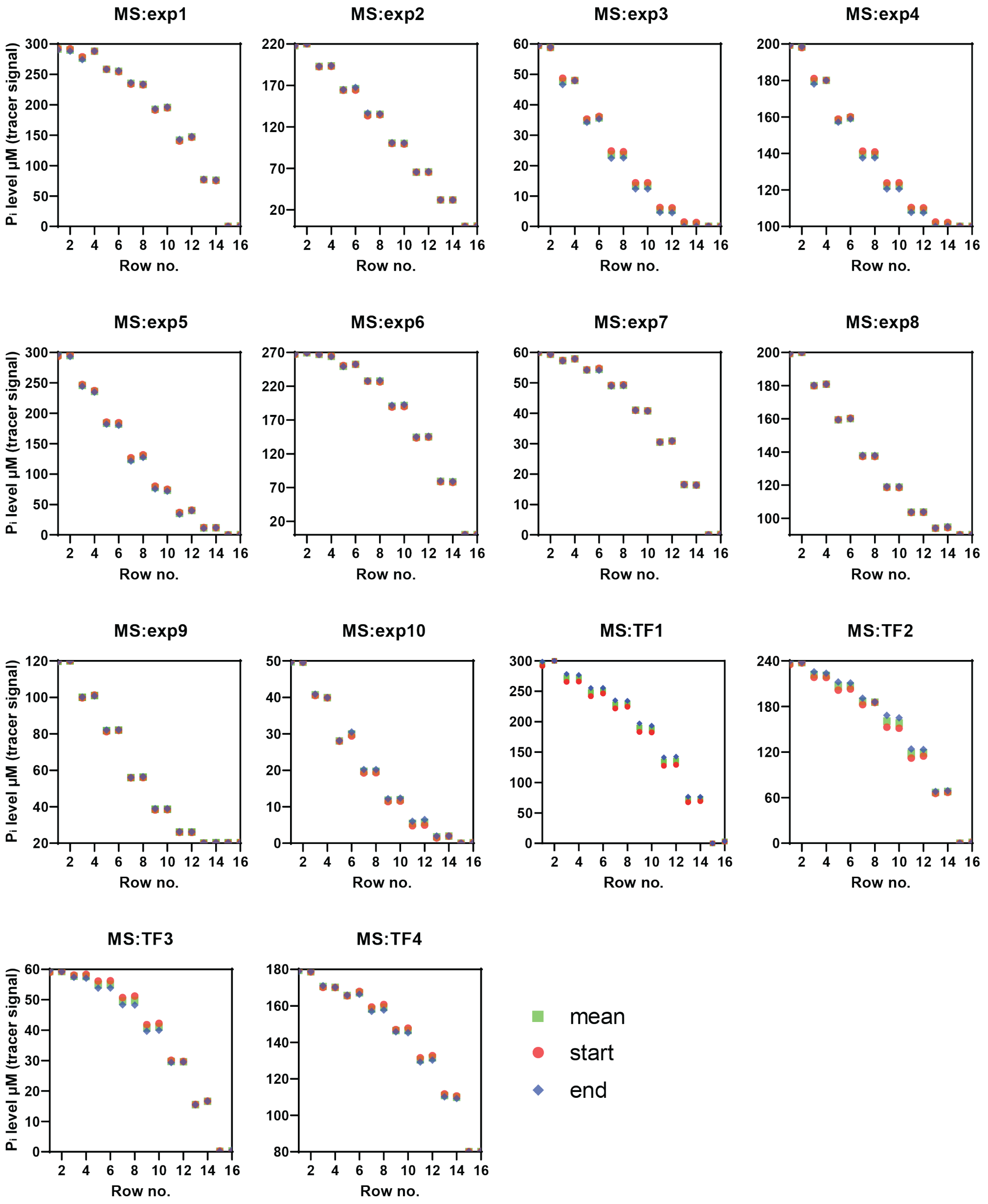

Figure S9: **Dilution generator calibrations for MICROSTAR experiments.** "Exp" indicates promoter characterisation experiments and "TF" denotes transcription factor localization experiments.

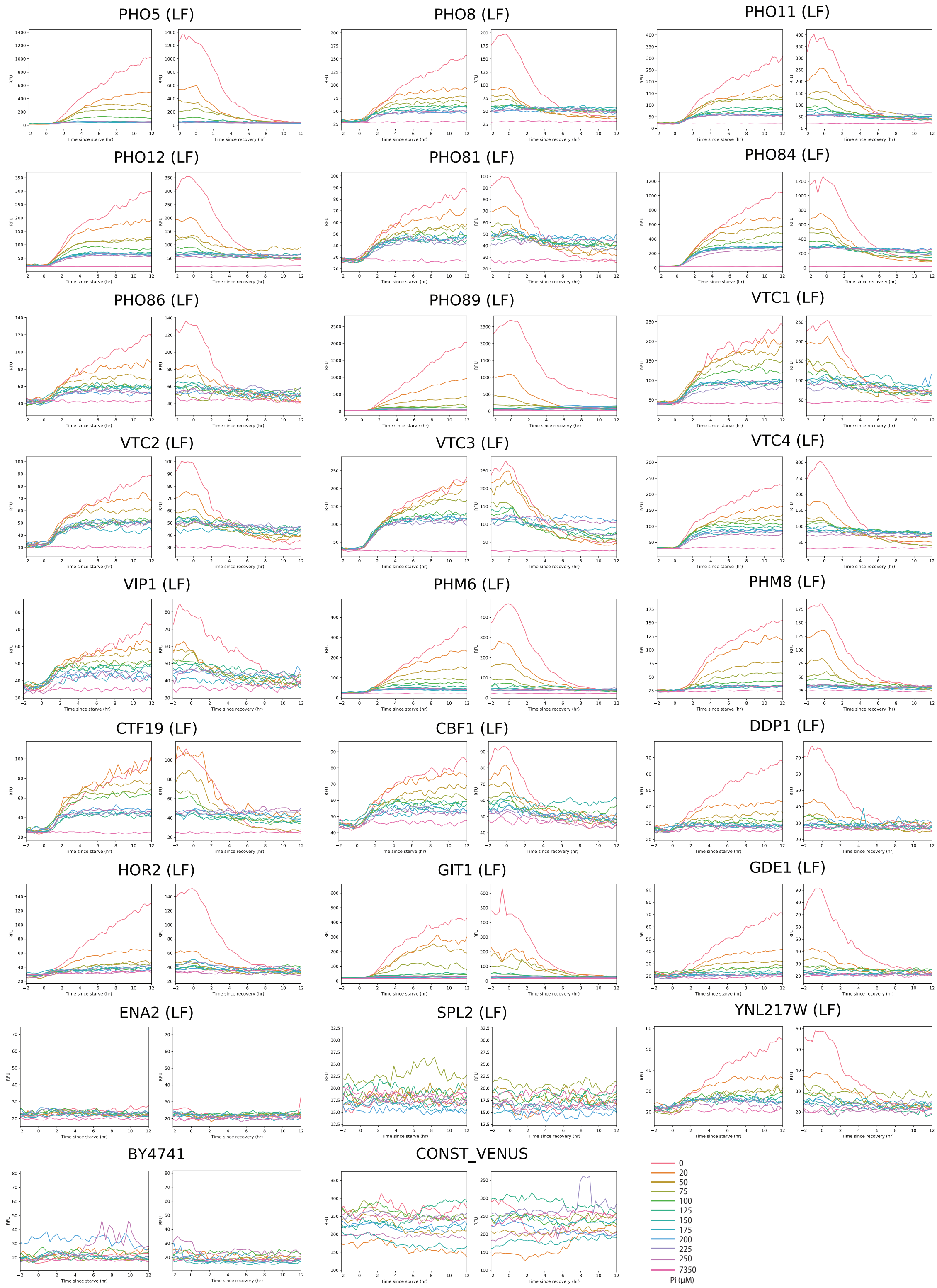

Figure S10: **Promoter expression over time at different $P_i$ concentrations for LF library strains measured on MCA.**

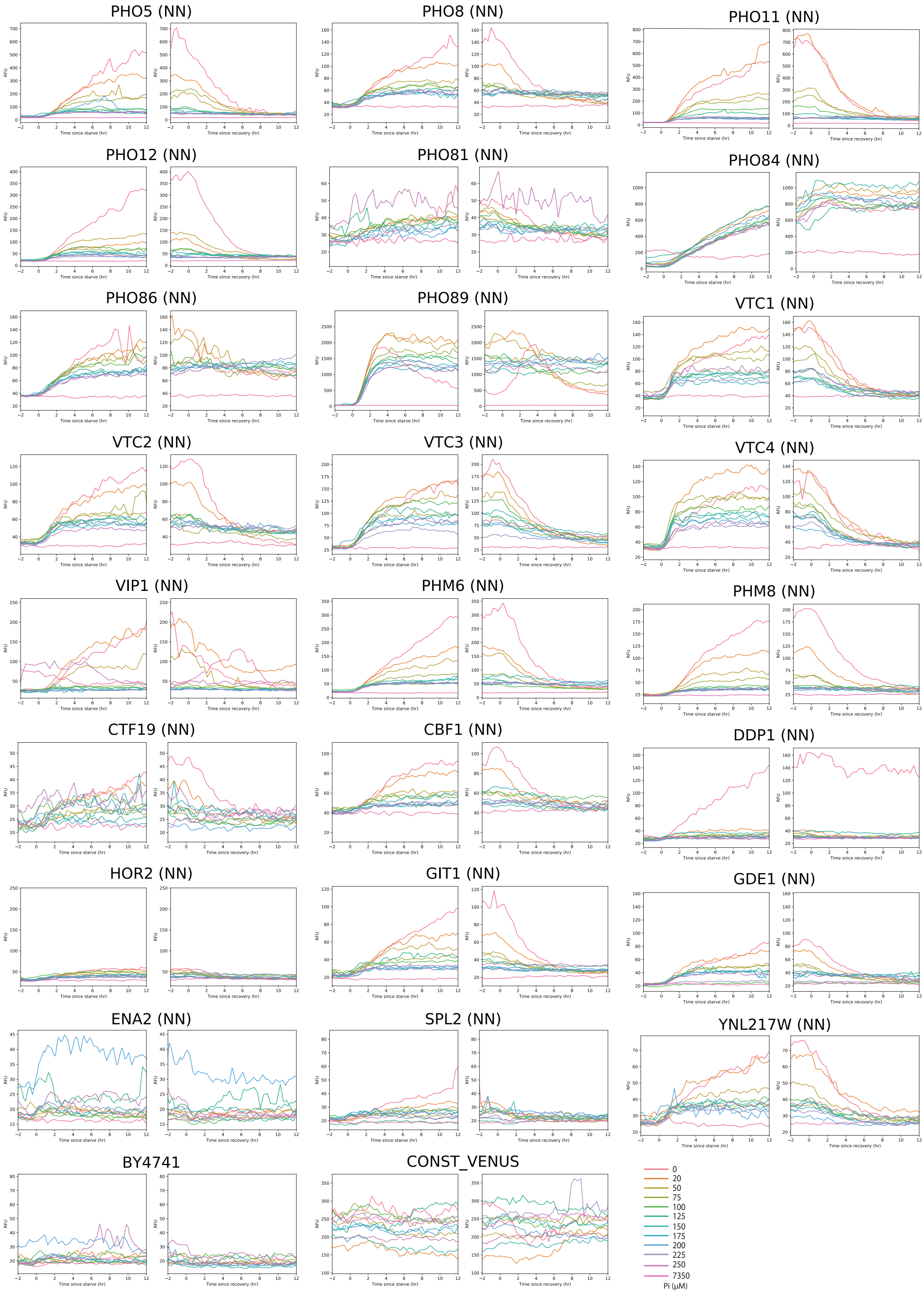

Figure S11: **Promoter expression over time at different $P_i$ concentrations for NN library strains measured on MCA.**

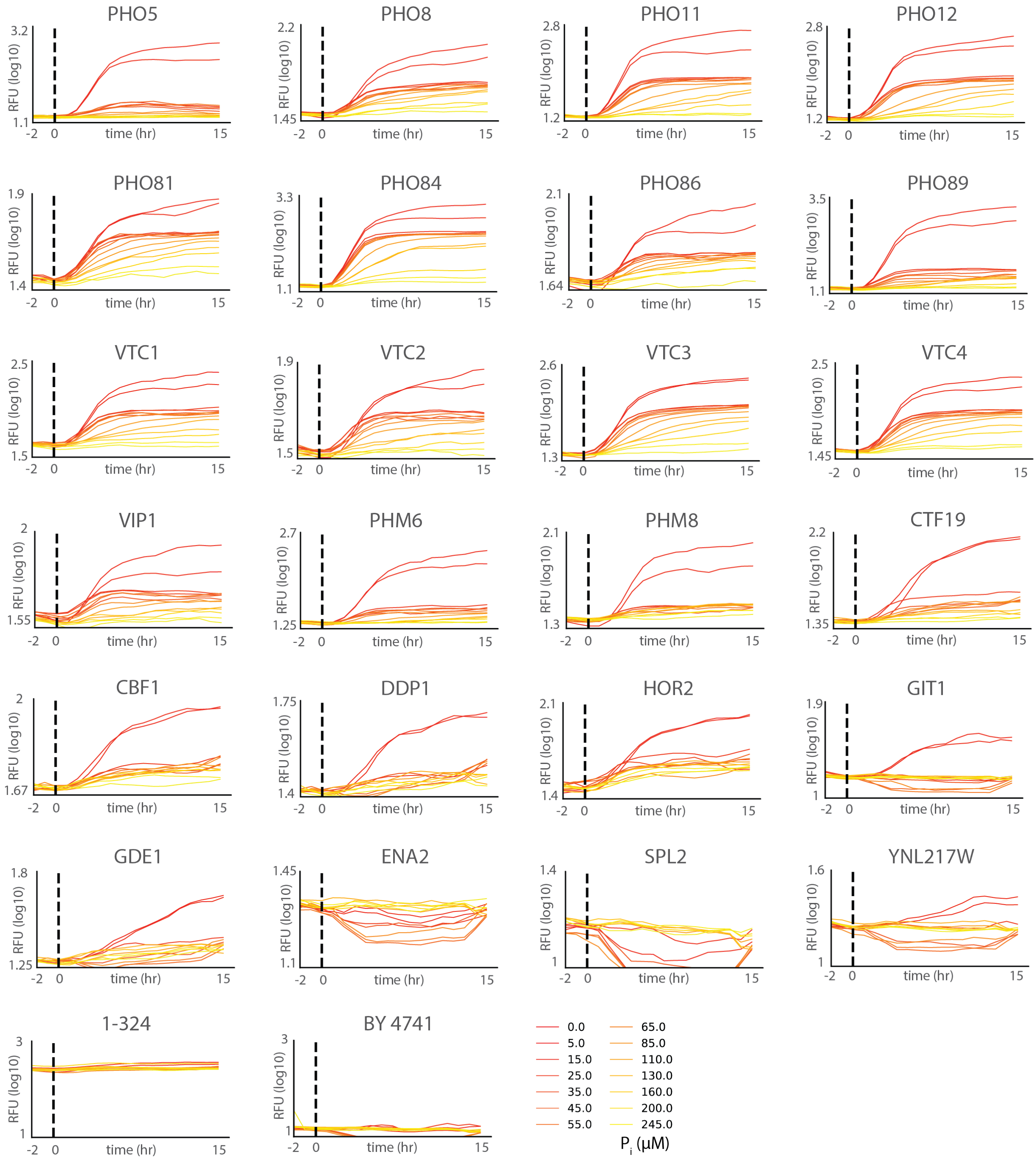

Figure S12: **Promoter expression over time at different $P_i$ concentrations for all strains measured on MICROSTAR.**

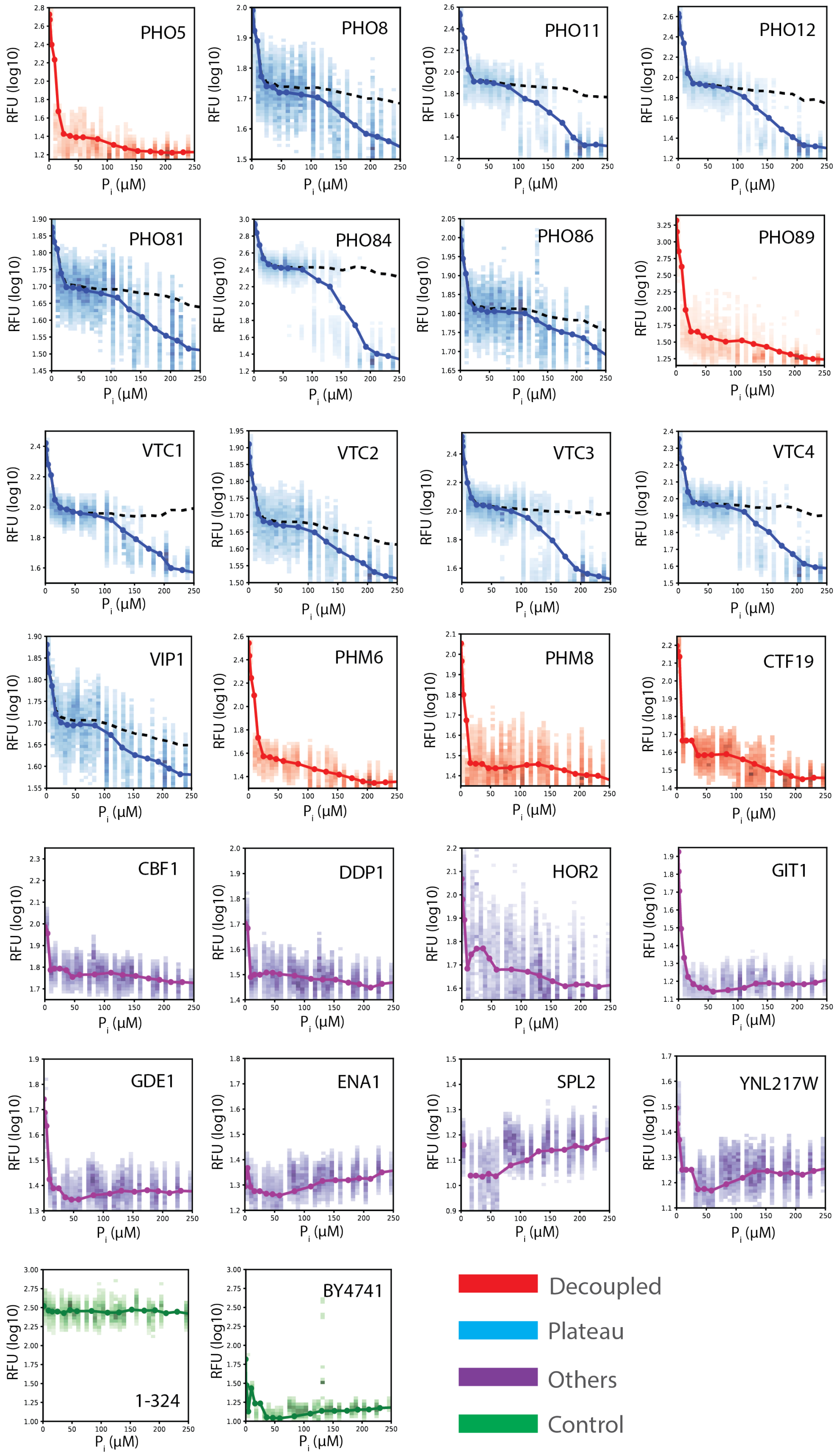

Figure S13: **Promoter expression versus $P_i$ for all strains measured on the MICROSTAR device.**

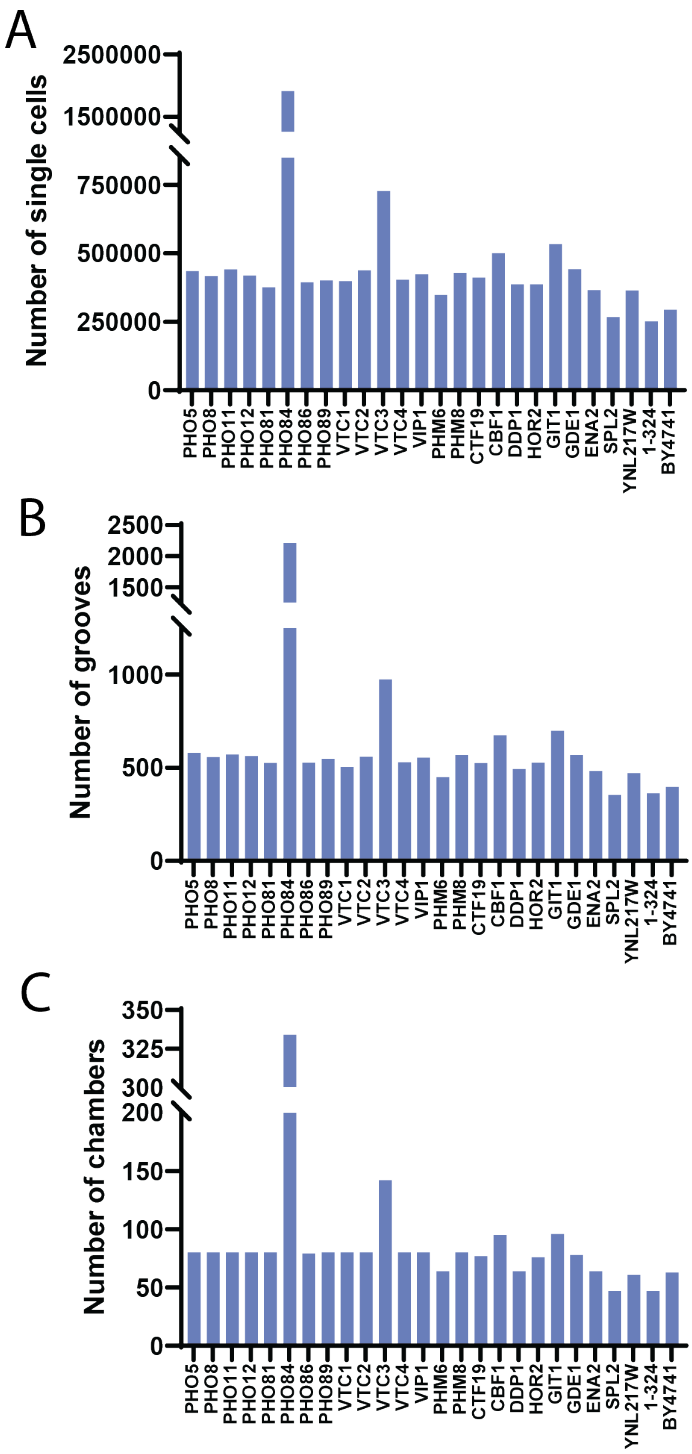

Figure S14: **MICROSTAR promoter expression sample sizes. A,** Number of single cells, **B,** number of grooves, and **C,** number of chambers measured for each strain in MICROSTAR promoter expression experiments.

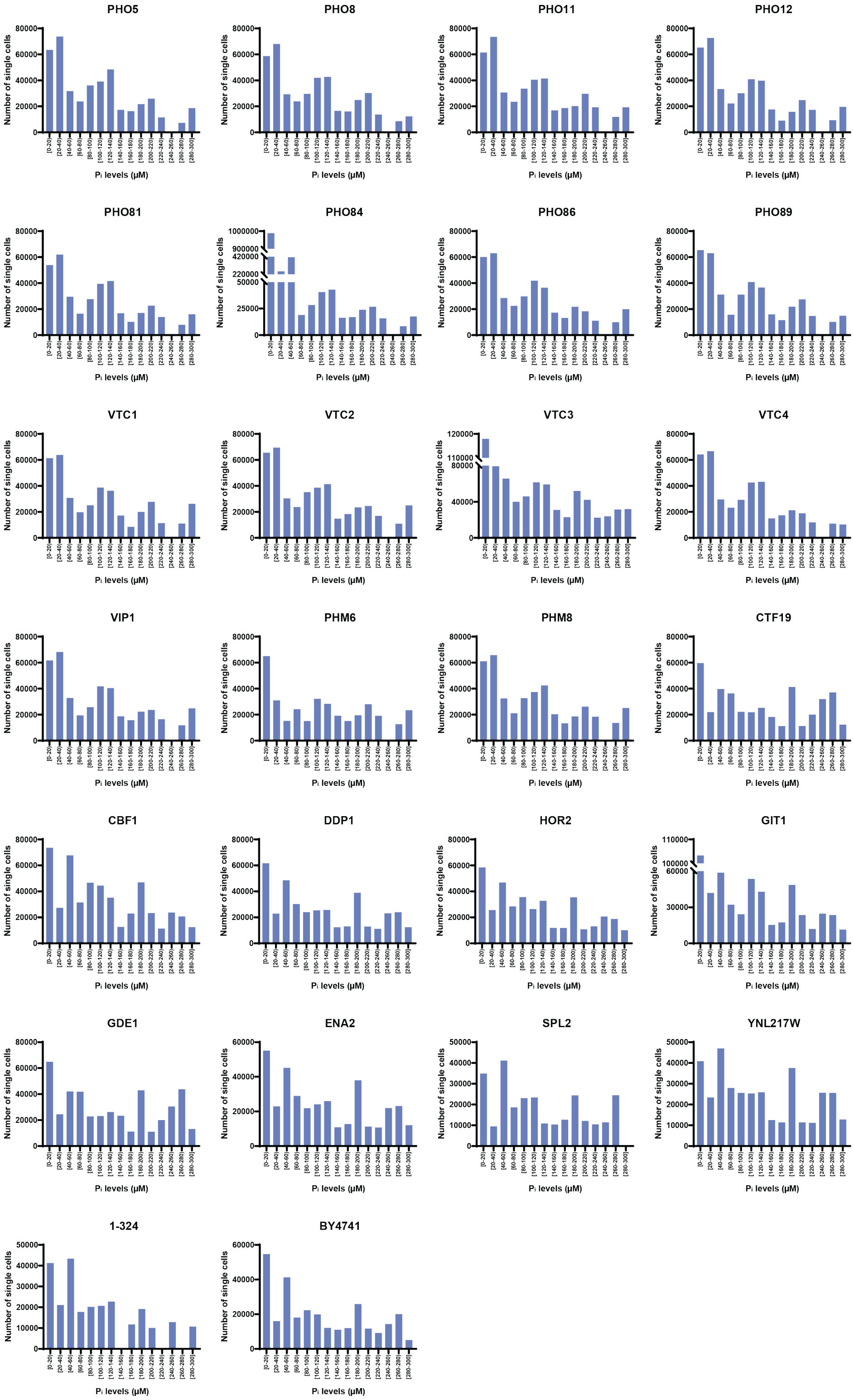

Figure S15: **Number of single cells measured across $P_i$ concentrations for each strain in MICROSTAR promoter expression experiments.**

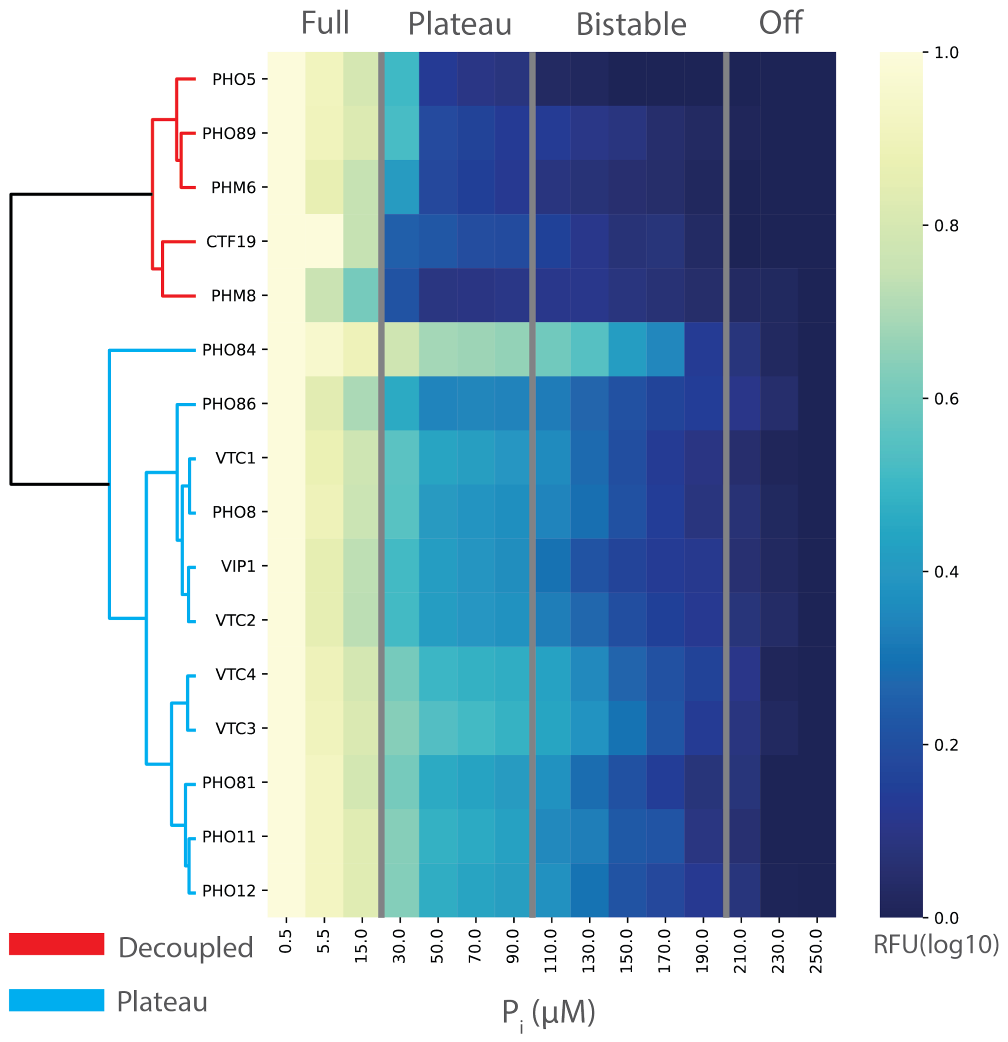

Figure S16: **Normalised promoter expression across $P_i$ levels and hierarchical clustering.** Normalised promoter activity levels of PHO promoters are plotted in a heatmap. The activation level is normalised by its maximum and minimum levels. The hierarchical tree on the left shows the clustering result. The promoters in red are clustered as decoupled promoters, and the promoters in blue are clustered as plateau promoters.

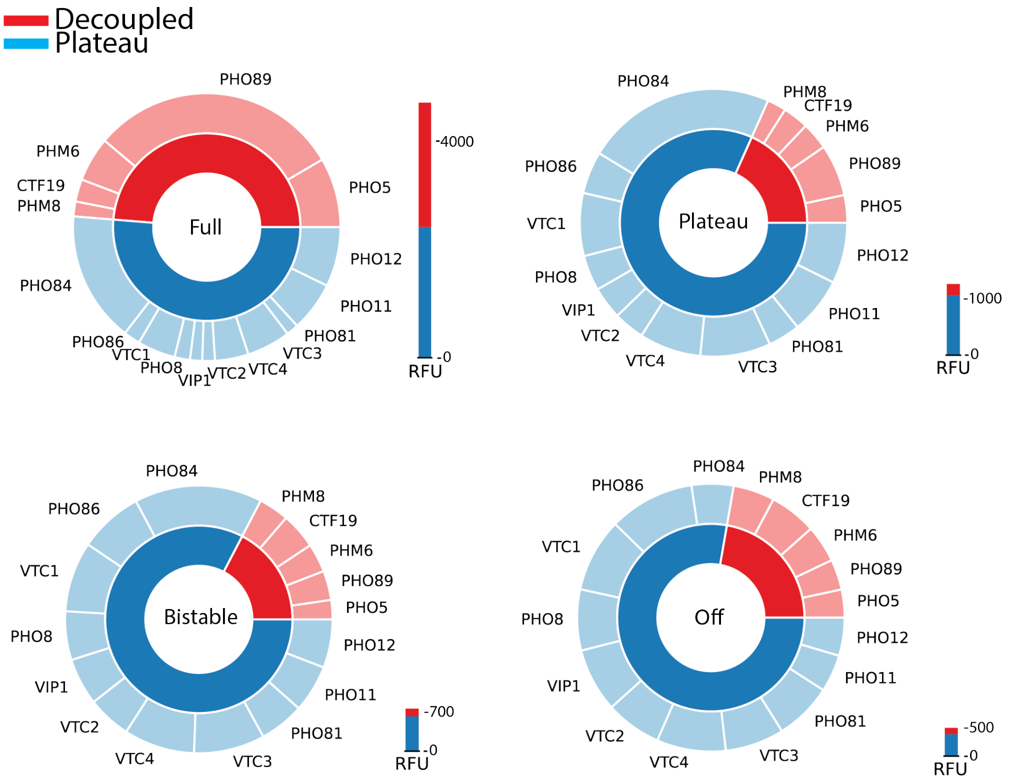

Figure S17: **Promoter activation levels during the different expression programs of the pho-regulon.** The relative activation levels of the PHO promoters in the four programs are shown as pie charts. The bar charts indicate the absolute level of activation. Promoters are grouped in either ”decoupled” or ”plateau” represented as red and blue, respectively.

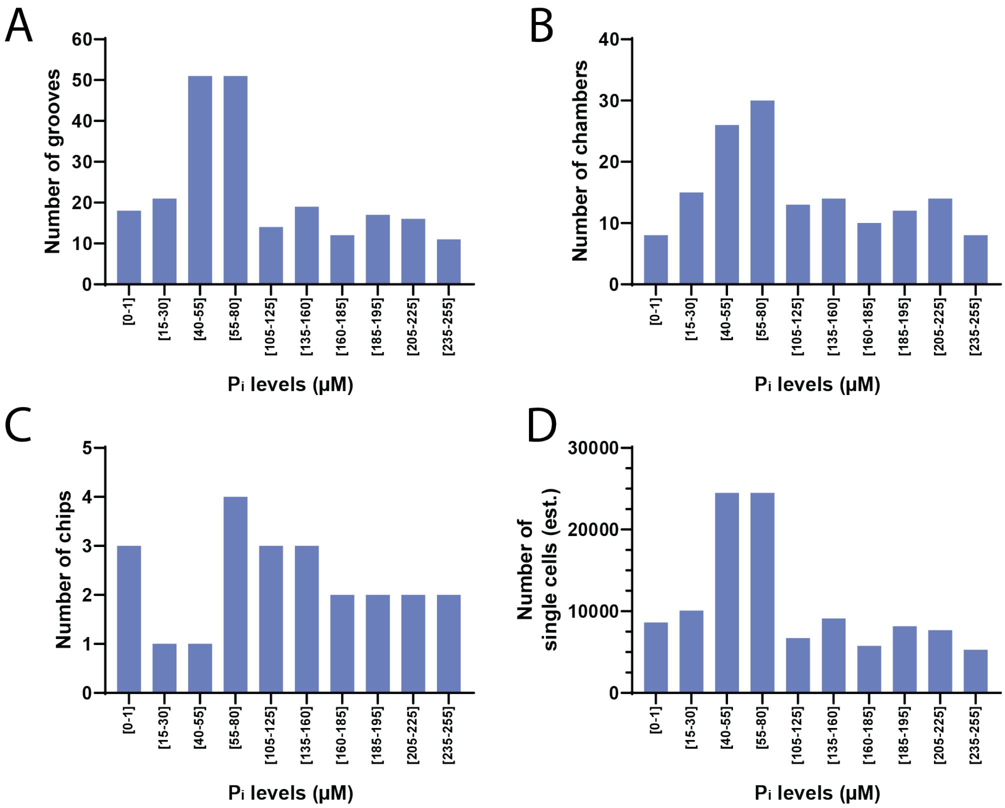

Figure S18: **Pho4p nuclear localization analysis sample size. A,** Number of grooves, **B,** number of chambers, **C,** number of chips, and **D,** estimated number of single cells measured in each $P_i$ concentration bin for Pho4p nuclear localization analysis.

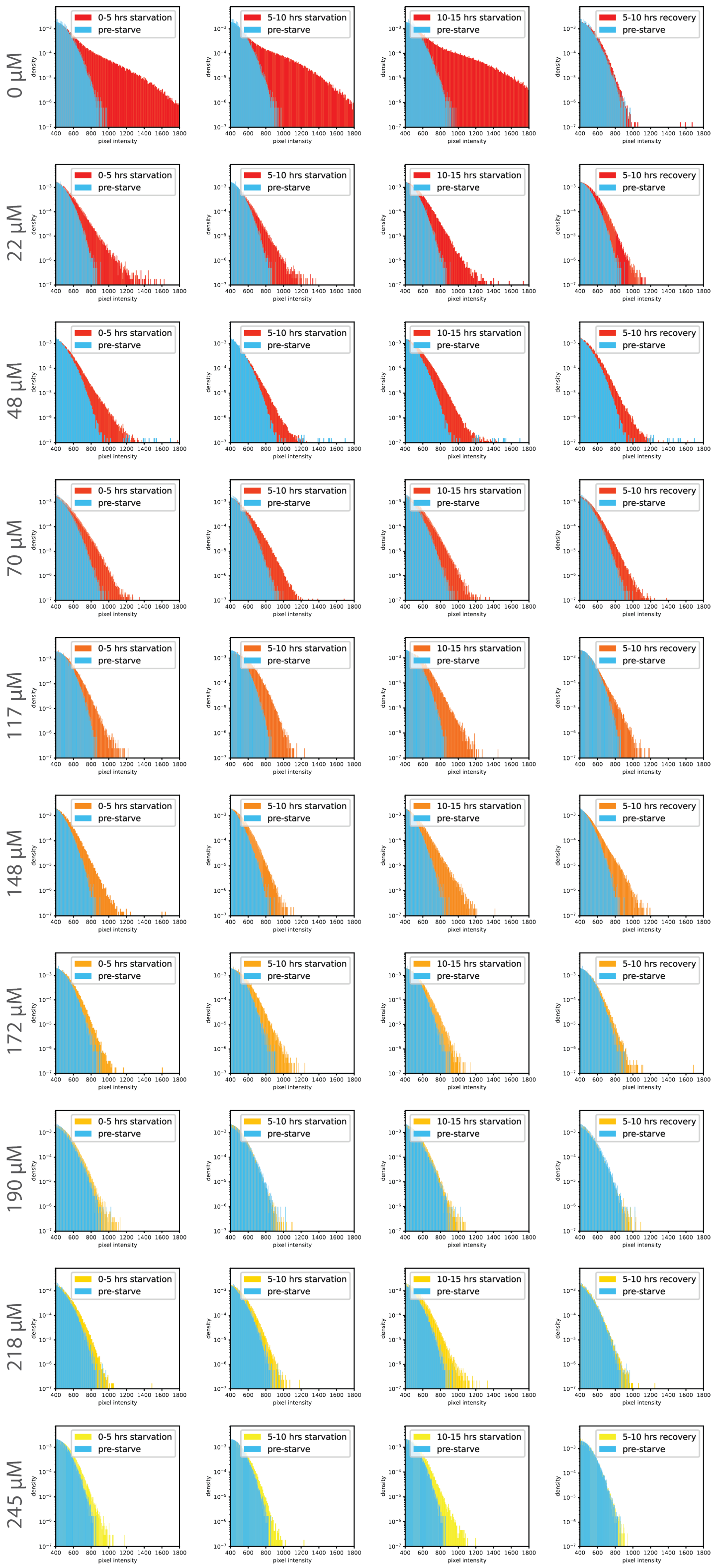

Figure S19: **Histograms of Pho4p signal at different $P_i$ levels.**

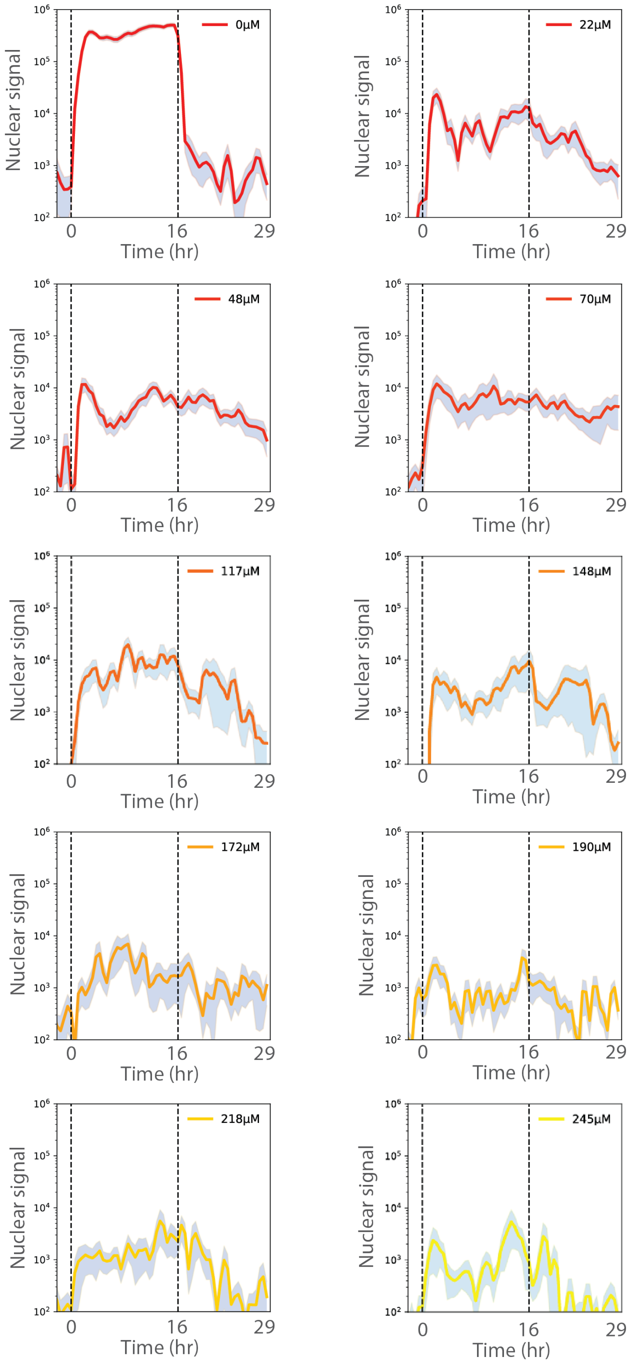

Figure S20: **Nuclear signal showing standard error (shaded region).**

### Chapter 3

### Supplementary Tables

Table S1: Genes in the phosphate gene regulatory network.

| Genes included in this study | | |
|---|---|---|
| **Gene** | **Location** | **Function** |
| *PHO4* | Chromosome VI | Basic helix-loop-helix (bHLH) transcription factor of the myc-family; activates transcription cooperatively with Pho2p in response to phosphate starvation; binding to 'CACGTG' motif is regulated by chromatin restriction, competitive binding of Cbf1p to the same DNA binding motif and cooperation with Pho2p; function is regulated by phosphorylation at multiple sites and by phosphate availability. |
| *PHO5* | Chromosome II | Repressible acid phosphatase; 1 of 3 repressible acid phosphatases that also mediates extracellular nucleotide-derived phosphate hydrolysis; secretory pathway derived cell surface glycoprotein; induced by phosphate starvation and coordinately regulated by Pho4p and PHO2. |
| *PHO89* | Chromosome II | Plasma membrane Na+/Pi cotransporter; active in early growth phase; similar to phosphate transporters of Neurospora crassa; transcription regulated by inorganic phosphate concentrations and Pho4pp; mutations in related human transporter genes hPit1 and hPit2 are associated with hyperphosphatemia-induced calcification of vascular tissue and familial idiopathic basal ganglia calcification. |
| *PHM6* | Chromosome IV | Phosphate metabolism protein 6. Expression is regulated by phosphate levels. |
| Pho84 | Chromosome XIII | High-affinity inorganic phosphate (Pi) transporter; also low-affinity manganese transporter; regulated by Pho4pp and Spt7p; mutation confers resistance to arsenate; exit from the ER during maturation requires Pho86p; cells overexpressing Pho84p accumulate heavy metals but do not develop symptoms of metal toxicity. |

| | | |
|---|---|---|
| *PHO11* | Chromosome I | One of three repressible acid phosphatases; glycoprotein that is transported to the cell surface by the secretory pathway; induced by phosphate starvation and coordinately regulated by Pho4p and PHO2; PHO11 has a paralog, PHO12, that arose from a segmental duplication. |
| *PHO12* | Chromosome VIII | One of three repressible acid phosphatases; glycoprotein that is transported to the cell surface by the secretory pathway; pregulated by phosphate starvation; PHO12 has a paralog, PHO11, that arose from a segmental duplication. |
| *PHO8* | Chromosome IV | Repressible vacuolar alkaline phosphatase; controls polyphosphate content; regulated by levels of Pi and by Pho4pp, Pho9p, Pho80p, Pho81p and Pho85p; dephosphorylates phosphotyrosyl peptides; contributes to NAD+ metabolism by producing nicotinamide riboside from NMN. |
| *PHO86* | Chromosome X | Endoplasmic reticulum (ER) resident protein; required for ER exit of the high-affinity phosphate transporter Pho84p, specifically required for packaging of Pho84p into COPII vesicles; protein abundance increases in response to DNA replication stress. |
| *VTC1* | Chromosome V | Regulatory subunit of the vacuolar transporter chaperone (VTC) complex; VTC complex is involved in membrane trafficking, vacuolar polyphosphate accumulation, microautophagy and non-autophagic vacuolar fusion; also has mRNA binding activity; protein abundance increases in response to DNA replication stress. |
| *VTC2* | Chromosome VI | Regulatory subunit of the vacuolar transporter chaperone (VTC) complex; involved in membrane trafficking, vacuolar polyphosphate accumulation, microautophagy and non-autophagic vacuolar fusion; VTC2 has a paralog, VTC3, that arose from the whole genome duplication. |

| | | |
|---|---|---|
| *VTC3* | Chromosome XVI | Regulatory subunit of the vacuolar transporter chaperone (VTC) complex; involved in membrane trafficking, vacuolar polyphosphate accumulation, microautophagy and non-autophagic vacuolar fusion; VTC3 has a paralog, VTC2, that arose from the whole genome duplication. |
| *VTC4* | Chromosome X | Vacuolar membrane polyphosphate polymerase; subunit of the vacuolar transporter chaperone (VTC) complex involved in synthesis and transfer of polyP to the vacuole; regulates membrane trafficking; role in non-autophagic vacuolar fusion; protein abundance increases in response to DNA replication stress. |
| *VIP1* | Chromosome XII | Bifunctional inositol pyrophosphate kinase and phosphatase; contains an N-terminal PP-InsP kinase domain that phosphorylates inositol hexakisphosphate and heptakisphosphate, and a C-terminal phosphatase domain that hydrolyzes both 1PP-InsP5 and 5PP-InsP5; IP7 levels decrease during starvation, suggesting a role for PP-InsP enzymes in Pi homeostasis; may regulate the dimorphic switch and the function of the cortical actin cytoskeleton. |
| *PHM8* | Chromosome V | Lysophosphatidic acid (LPA) phosphatase, nucleotidase; principle and physiological nucleotidase working on GMP, UMP and CMP; involved in LPA hydrolysis in response to phosphate starvation and ribose salvage pathway; phosphatase activity is soluble and Mg2+ dependent; expression is induced by low phosphate levels and by inactivation of Pho85p; repressed by Gcn4p under normal conditions; PHM8 has a paralog, SDT1, that arose from the whole genome duplication. |

| | | |
|---|---|---|
| *HOR2* | Chromosome V | DL-glycerol-3-phosphate phosphatase involved in glycerol biosynthesis; also known as glycerol-1-phosphatase; induced in response to hyperosmotic or oxidative stress, and during diauxic shift; HOR2 has a paralog, GPP1, that arose from the whole genome duplication. |
| *CTF19* | Chromosome XVI | Outer kinetochore protein, needed for accurate chromosome segregation; component of kinetochore sub-complex COMA (Ctf19p, Okp1p, Mcm21p, Ame1p) that functions as platform for kinetochore assembly; required for spindle assembly checkpoint; minimizes potentially deleterious centromere-proximal crossovers by preventing meiotic DNA break formation proximal to centromere; homolog of human centromere constitutive-associated network (CCAN) subunit CENP-P and fission yeast fta2. |
| *CBF1* | Chromosome X | Basic helix-loop-helix (bHLH) protein; forms homodimer to bind E-box consensus sequence CACGTG present at MET gene promoters and centromere DNA element I (CDEI); affects nucleosome positioning at this motif; associates with other transcription factors such as Met4p and Isw1p to mediate transcriptional activation or repression; associates with kinetochore proteins, required for chromosome segregation; protein abundance increases in response to DNA replication stress. |
| *ENA1* | Chromosome IV | P-type ATPase sodium pump; involved in Na+ and Li+ efflux to allow salt tolerance. |
| *ENA2* | Chromosome IV | P-type ATPase sodium pump; involved in Na+ efflux to allow salt tolerance; likely not involved in Li+ efflux. |
| *GDE1* | Chromosome XVI | Glycerophosphocholine (GroPCho) phosphodiesterase; hydrolyzes GroPCho to choline and glycerolphosphate, for use as a phosphate source and as a precursor for phosphocholine synthesis; may interact with ribosomes. |

| | | |
|---|---|---|
| *DDP1* | Chromosome XV | Diadenosine and diphosphoinositol polyphosphate phosphatase; hydrolyzes diphosphorylated inositol polyphosphates and diadenosine polyphosphates; high specificity for diadenosine hexa- and pentaphosphates; contains endopolyphosphatase activity with a high affinity for polyphosphates, an activity also observed for its human DIPP homologs; possesses mRNA decapping activity; nudix hydrolase family member; protein abundance increases in response to DNA replication stress. |
| *GIT1* | Chromosome III | Plasma membrane permease; mediates uptake of glycerophosphoinositol and glycerophosphocholine as sources of the nutrients inositol and phosphate; expression and transport rate are regulated by phosphate and inositol availability. |
| *SPL2* | Chromosome VIII | Protein with similarity to cyclin-dependent kinase inhibitors; downregulates low-affinity phosphate transport during phosphate limitation by targeting Pho87p to the vacuole; upstream region harbors putative hypoxia response element (HRE) cluster; overproduction suppresses a plc1 null mutation; promoter shows an increase in Snf2p occupancy after heat shock; GFP-fusion protein localizes to the cytoplasm. |
| *YJL119C* | Chromosome X | Dubious open reading frame; unlikely to encode a functional protein, based on available experimental and comparative sequence data. |
| *YAR070C* | Chromosome I | Dubious open reading frame; unlikely to encode a protein, based on available experimental and comparative sequence data; YAR070C has a paralog, YHR214C-B, that arose from a segmental duplication. |

| | | |
|---|---|---|
| *YNL217W* | Chromosome XIV | Zn2+-dependent endopolyphosphatase; required with PPN1 to mobilize polyphosphate stores in response to phosphate starvation; member of the PPP-superfamily of metalloproteases; localizes to the vacuolar lumen via the MVB pathway; null mutant is highly sensitive to azaserine and resistant to sodium-O-vandate. |
| Reference: *Saccharomyces* Genome Database (SGD), https://www.yeastgenome.org/ | | |

Table S2: List of strains analyzed in each MICROSTAR experiment

| | exp1 | exp2 | exp3 | exp4 | exp5 | exp6 | exp7 | exp8 | exp9 | exp10 |
|---|---|---|---|---|---|---|---|---|---|---|
| PHO5 | ■ | ■ | ■ | ■ | | | | | ■ | |
| PHO8 | ■ | ■ | ■ | ■ | | | | | ■ | |
| PHO11 | ■ | ■ | ■ | ■ | | | | | ■ | |
| PHO12 | ■ | ■ | ■ | ■ | | | | | ■ | |
| PHO81 | ■ | ■ | ■ | ■ | | | | | ■ | |
| PHO84 | ■ | ■ | ■ | ■ | | | | | ■ | ■ |
| PHO86 | ■ | ■ | ■ | ■ | | | | | ■ | |
| PHO89 | ■ | ■ | ■ | ■ | | | | | ■ | |
| VTC1 | ■ | ■ | ■ | ■ | | | | | ■ | |
| VTC2 | ■ | ■ | ■ | ■ | | | | | ■ | |
| VTC3 | ■ | ■ | ■ | ■ | ■ | ■ | ■ | ■ | ■ | |
| VTC4 | ■ | ■ | ■ | ■ | | | | | ■ | |
| VIP1 | ■ | ■ | ■ | ■ | | | | | ■ | |
| PHM6 | ■ | ■ | ■ | ■ | | | | | | |
| PHM8 | ■ | ■ | ■ | ■ | | | | | ■ | |
| CTF19 | | | | | ■ | ■ | ■ | ■ | | |
| CBF1 | | | | | ■ | ■ | ■ | ■ | | |
| DDP1 | | | | | ■ | ■ | ■ | | ■ | |
| HOR2 | | | | | ■ | ■ | ■ | ■ | | |
| GIT1 | | | ■ | ■ | ■ | ■ | ■ | ■ | | |
| GDE1 | | | | | ■ | ■ | ■ | ■ | | |
| ENA2 | | | | | ■ | ■ | ■ | ■ | | |
| SPL2 | | | | | ■ | ■ | ■ | ■ | | |
| YNL217W | | | | | ■ | ■ | ■ | ■ | | |
| 1-324 | | | | | ■ | | ■ | ■ | | |
| BY4741 | | | | | ■ | ■ | ■ | ■ | | |

Table S3: Media, chemicals, enzymes, etc. used in this study.

| **Reagent or Kit** | **Source** | **Identifier** |
|---|---|---|
| YPD broth | Sigma Aldrich | Cat# Y1375-1KG |
| Glucose | Sigma Aldrich | Cat# G8270-1KG |
| LB powder | AppliChem GmbH | Cat# A0954 |
| Agar | Sigma Aldrich | Cat# 05039-500G |
| Chloramphenicol | Sigma Aldrich | Cat# C0378-5G |
| Ampicillin | Roche | Cat# 10835242001 |
| Kanamycin Sulfate | Life Technologies | Cat# 11815024 |
| Yeast Synthetic Drop-out Medium Supplements | Sigma Aldrich | Cat# Y1501-20G |
| Yeast Nitrogen Base Without Amino Acids | Sigma Aldrich | Cat# Y0626-1KG |
| Uracil | Sigma Aldrich | Cat# U1128-25G |
| EZSolution 5-Fluoroorotic acid monohydrate (5-FOA) | Biovision | Cat# 2797-10 |
| YNB with ammonium sulfate, powder | MP Biomedicals | Cat# MP4027-812 |
| Complete Supplement Mixture (CSM), powder | MP Biomedicals | Cat# MPB-114500022 |
| Adenine hemisulfate salt | Sigma Aldrich | Cat# A3159-25G |
| L-Tryptophan | Sigma Aldrich | Cat# 1083740100 |
| Monobasic potassium phosphate | Sigma Aldrich | Cat# P5655-100G |
| Sodium chloride | Sigma Aldrich | Cat# S9888-25G |
| Potassium chloride | Sigma Aldrich | Cat# P5405-250G |
| Phire Plant Direct PCR Master Mix | Molecular Dimensions | Cat# 133083 |
| NeXtalStock PEG 3,350 (200) - 200 ml | Sigma Aldrich | Cat# 05039-500G |
| QIAprep Spin Miniprep Kit (250) | QIAGEN | Cat# 27106 |
| Wizard SV Gel and PCR Clean-Up System | Promega Corp | Cat# A9281 |
| QIAquick Gel Extraction Kit (50) | QIAGEN | Cat# 28704 |
| BsaI-HF | New England Biolabs Inc. | Cat# R3733L |
| BsmBI-v2 | New England Biolabs Inc. | Cat# R0739L |
| EcoRV | New England Biolabs Inc. | Cat# R0195L |
| T4 DNA ligase | New England Biolabs Inc. | Cat# M0202L |
| NotI-HF (High Fidelity) | New England Biolabs Inc. | Cat# R3189L |
| NEB 10-beta Competent E. coli (High Efficiency) | New England Biolabs Inc. | Cat# C3019H |
| Midori Green Advance (1 ml) | Nippon Genetics Europe | Cat# MG04 |
| UltraPure Salmon Sperm DNA Solution | Life Technologies | Cat# 15632011 |

| | | |
|---|---|---|
| 500 mL Vacuum Filter/Storage Bottle System | Corning | Cat# 431097 |

Table S4: Plasmids used or generated in this study.

| Plasmid ID | Strain Name | Source | Identifier |
|---|---|---|---|
| pYTK001-096 | Yeast MoClo Toolkit | John Dueber lab | Addgene# 1000000061 |
| pTY001 | LF_PHO5_yoEGFP_tADH1 | This work | N/A |
| pTY002 | LF_PHO89_yoEGFP_tADH1 | This work | N/A |
| pTY003 | LF_PHM6_yoEGFP_tADH1 | This work | N/A |
| pTY004 | LF_PHO84_yoEGFP_tADH1 | This work | N/A |
| pTY005 | LF_PHO11_yoEGFP_tADH1 | This work | N/A |
| pTY006 | LF_PHO12_yoEGFP_tADH1 | This work | N/A |
| pTY007 | LF_VTC1_yoEGFP_tADH1 | This work | N/A |
| pTY008 | LF_VTC3_yoEGFP_tADH1 | This work | N/A |
| pTY009 | LF_VTC4_yoEGFP_tADH1 | This work | N/A |
| pTY010 | LF_VTC2_yoEGFP_tADH1 | This work | N/A |
| pTY011 | LF_PHO8_yoEGFP_tADH1 | This work | N/A |
| pTY012 | LF_PHO86_yoEGFP_tADH1 | This work | N/A |
| pTY013 | LF_VIP1_yoEGFP_tADH1 | This work | N/A |
| pTY014 | LF_PHO81_yoEGFP_tADH1 | This work | N/A |
| pTY015 | LF_HOR2_yoEGFP_tADH1 | This work | N/A |
| pTY016 | LF_PHM8_yoEGFP_tADH1 | This work | N/A |
| pTY017 | LF_CTF19_yoEGFP_tADH1 | This work | N/A |
| pTY018 | LF_SPL2_yoEGFP_tADH1 | This work | N/A |
| pTY019 | LF_CBF1_yoEGFP_tADH1 | This work | N/A |
| pTY020 | LF_DDP1_yoEGFP_tADH1 | This work | N/A |
| pTY021 | LF_ENA1_yoEGFP_tADH1 | This work | N/A |
| pTY022 | LF_ENA2_yoEGFP_tADH1 | This work | N/A |
| pTY023 | LF_GDE1_yoEGFP_tADH1 | This work | N/A |
| pTY024 | LF_GIT1_yoEGFP_tADH1 | This work | N/A |
| pTY025 | LF_YAR070C_yoEGFP_tADH1 | This work | N/A |
| pTY026 | LF_YJL119C_yoEGFP_tADH1 | This work | N/A |
| pTY027 | LF_YNL217W_yoEGFP_tADH1 | This work | N/A |
| pTY028 | NN_PHO5_yoEGFP_tPHO5 | This work | N/A |
| pTY029 | NN_PHO89_yoEGFP_tPHO89 | This work | N/A |
| pTY030 | NN_PHM6_yoEGFP_tPHM6 | This work | N/A |

| | | | |
|---|---|---|---|
| pTY031 | NN_PHO84_yoEGFP_tPHO84 | This work | N/A |
| pTY032 | NN_PHO11_yoEGFP_tPHO11 | This work | N/A |
| pTY033 | NN_PHO12_yoEGFP_tPHO12 | This work | N/A |
| pTY034 | NN_VTC1_yoEGFP_tVTC1 | This work | N/A |
| pTY035 | NN_VTC3_yoEGFP_tVTC3 | This work | N/A |
| pTY036 | NN_VTC4_yoEGFP_tVTC4 | This work | N/A |
| pTY037 | NN_VTC2_yoEGFP_tVTC2 | This work | N/A |
| pTY038 | NN_PHO8_yoEGFP_tPHO8 | This work | N/A |
| pTY039 | NN_PHO86_yoEGFP_tPHO86 | This work | N/A |
| pTY040 | NN_VIP1_yoEGFP_tVIP1 | This work | N/A |
| pTY041 | NN_PHO81_yoEGFP_tPHO81 | This work | N/A |
| pTY042 | NN_HOR2_yoEGFP_tHOR2 | This work | N/A |
| pTY043 | NN_PHM8_yoEGFP_tPHM8 | This work | N/A |
| pTY044 | NN_CTF19_yoEGFP_tCTF19 | This work | N/A |
| pTY045 | NN_SPL2_yoEGFP_tSPL2 | This work | N/A |
| pTY046 | NN_CBF1_yoEGFP_tCBF1 | This work | N/A |
| pTY047 | NN_DDP1_yoEGFP_tDDP1 | This work | N/A |
| pTY048 | NN_ENA1_yoEGFP_tENA1 | This work | N/A |
| pTY049 | NN_ENA2_yoEGFP_tENA2 | This work | N/A |
| pTY050 | NN_GDE1_yoEGFP_tGDE1 | This work | N/A |
| pTY051 | NN_GIT1_yoEGFP_tGIT1 | This work | N/A |
| pTY052 | NN_YAR070C_yoEGFP_tYAR070C | This work | N/A |
| pTY053 | NN_YJL119C_yoEGFP_tYJL119C | This work | N/A |
| pTY054 | NN_YNL217W_yoEGFP_tYNL217W | This work | N/A |
| pTY055 | LYS2 sgRNA1 | This work | N/A |
| pTY056 | LYS2 sgRNA2 | This work | N/A |
| pTY057 | pWS158 [o] | Ellis lab | Addgene# 90517 |
| pTY058 | pWS082 - sgRNA Entry Vector | Ellis lab | Addgene# 90516 |
| pTY059 | VTC3 (YPL019C) sgRNA1 | This work | N/A |
| pTY060 | VTC3 (YPL019C) sgRNA2 | This work | N/A |
| pTY061 | CTF19 (YPL018W) sgRNA1 | This work | N/A |
| pTY062 | CTF19 (YPL018W) sgRNA2 | This work | N/A |
| pTY063 | PHO84 (YML123C) sgRNA2 | This work | N/A |

| | | | |
|---|---|---|---|
| pTY064 | PHO84 (YML123C) sgRNA1 | This work | N/A |
| pTY065 | VIP1 (YLR410W) sgRNA1 | This work | N/A |
| pTY066 | VIP1 (YLR410W) sgRNA2 | This work | N/A |
| pTY067 | CBF1 (YJR060W) sgRNA2 | This work | N/A |
| pTY068 | CBF1 (YJR060W) sgRNA1 | This work | N/A |
| pTY069 | YJL119C (YJL119C) sgRNA1 | This work | N/A |
| pTY070 | YJL119C (YJL119C) sgRNA2 | This work | N/A |
| pTY071 | VTC4 (YJL012C) sgRNA2 | This work | N/A |
| pTY072 | VTC4 (YJL012C) sgRNA1 | This work | N/A |
| pTY073 | SPL2 (YHR136C) sgRNA1 | This work | N/A |
| pTY074 | SPL2 (YHR136C) sgRNA2 | This work | N/A |
| pTY075 | PHO5 (YBR093C) sgRNA1 | This work | N/A |
| pTY076 | PHO5 (YBR093C) sgRNA2 | This work | N/A |
| pTY077 | PHO81 (YGR233C) sgRNA1 | This work | N/A |
| pTY078 | PHO81 (YGR233C) sgRNA2 | This work | N/A |
| pTY079 | VTC2 (YFL004W) sgRNA1 | This work | N/A |
| pTY080 | VTC2 (YFL004W) sgRNA2 | This work | N/A |
| pTY081 | PHO8 (YDR481C) sgRNA1 | This work | N/A |
| pTY082 | PHO8 (YDR481C) sgRNA2 | This work | N/A |
| pTY083 | PHM8 (YER037W) sgRNA1 | This work | N/A |
| pTY084 | PHM8 (YER037W) sgRNA2 | This work | N/A |
| pTY085 | GIT1 (YCR098C) sgRNA2 | This work | N/A |
| pTY086 | GIT1 (YCR098C) sgRNA1 | This work | N/A |
| pTY087 | PHO89 (YBR296C) sgRNA1 | This work | N/A |
| pTY088 | PHO89 (YBR296C) sgRNA2 | This work | N/A |
| pTY089 | YAR071W (PHO11) sgRNA (OT) | This work | N/A |
| pTY090 | YDR039C (ENA2) sgRNA (OT) | This work | N/A |
| pTY091 | YAR070C sgRNA (OT) | This work | N/A |
| pTY092 | YHR215W (PHO12) sgRNA | This work | N/A |
| pTY093 | YDR040C (ENA1) sgRNA | This work | N/A |
| pTY094 | YDR281C (PHM6) sgRNA | This work | N/A |
| pTY095 | YER062C (HOR2) sgRNA | This work | N/A |
| pTY096 | YER072W (VTC1) sgRNA | This work | N/A |

| | | | |
|---|---|---|---|
| pTY097 | YJL117W (PHO86) sgRNA | This work | N/A |
| pTY098 | YOR163W (DDP1) sgRNA | This work | N/A |
| pTY099 | YNL217W sgRNA | This work | N/A |
| pTY100 | YPL110C (GDE1) sgRNA | This work | N/A |
| pTY101 | Pho4p knockout sgRNA1 | This work | N/A |
| pTY102 | Pho4p knockout sgRNA2 | This work | N/A |
| pTY103 | mScarlet-i Pho4p N-tagged | This work | N/A |
| pTY104 | Pho4p mScarlet-i C-tagged | This work | N/A |

Table S5: Yeast strains used or generated in this study.

| Strain ID | Strain Name | Cassette | Source | Identifier |
|---|---|---|---|---|
| sTY001 | BY4741 | N/A | HorizonDiscovery | Cat# YSC1048 |
| sTY002 | mScarleti-Pho4p | N/A | This work | N/A |
| sTY003 | Yeast GFP library, Pho87-GFP | N/A | ThermoFisher | Cat# 95700 |
| sTY004 | Yeast GFP library, Pho90-GFP | N/A | ThermoFisher | Cat# 95700 |
| sTY005 | Yeast GFP library, SPL2-GFP | N/A | ThermoFisher | Cat# 95700 |
| sTY006 | LF library, PHO5_yoEGFP_tADH1 | pTY001 | This work | N/A |
| sTY007 | LF library, PHO89_yoEGFP_tADH1 | pTY002 | This work | N/A |
| sTY008 | LF library, PHM6_yoEGFP_tADH1 | pTY003 | This work | N/A |
| sTY009 | LF library, PHO84_yoEGFP_tADH1 | pTY004 | This work | N/A |
| sTY010 | LF library, PHO11_yoEGFP_tADH1 | pTY005 | This work | N/A |
| sTY011 | LF library, PHO12_yoEGFP_tADH1 | pTY006 | This work | N/A |
| sTY012 | LF library, VTC1_yoEGFP_tADH1 | pTY007 | This work | N/A |
| sTY013 | LF library, VTC3_yoEGFP_tADH1 | pTY008 | This work | N/A |
| sTY014 | LF library, VTC4_yoEGFP_tADH1 | pTY009 | This work | N/A |
| sTY015 | LF library, VTC2_yoEGFP_tADH1 | pTY010 | This work | N/A |
| sTY016 | LF library, PHO8_yoEGFP_tADH1 | pTY011 | This work | N/A |
| sTY017 | LF library, PHO86_yoEGFP_tADH1 | pTY012 | This work | N/A |
| sTY018 | LF library, VIP1_yoEGFP_tADH1 | pTY013 | This work | N/A |
| sTY019 | LF library, PHO81_yoEGFP_tADH1 | pTY014 | This work | N/A |
| sTY020 | LF library, HOR2_yoEGFP_tADH1 | pTY015 | This work | N/A |
| sTY021 | LF library, PHM8_yoEGFP_tADH1 | pTY016 | This work | N/A |
| sTY022 | LF library, CTF19_yoEGFP_tADH1 | pTY017 | This work | N/A |
| sTY023 | LF library, SPL2_yoEGFP_tADH1 | pTY018 | This work | N/A |
| sTY024 | LF library, CBF1_yoEGFP_tADH1 | pTY019 | This work | N/A |
| sTY025 | LF library, DDP1_yoEGFP_tADH1 | pTY020 | This work | N/A |
| sTY026 | LF library, ENA1_yoEGFP_tADH1 | pTY021 | This work | N/A |
| sTY027 | LF library, ENA2_yoEGFP_tADH1 | pTY022 | This work | N/A |
| sTY028 | LF library, GDE1_yoEGFP_tADH1 | pTY023 | This work | N/A |
| sTY029 | LF library, GIT1_yoEGFP_tADH1 | pTY024 | This work | N/A |
| sTY030 | LF library, YAR070C_yoEGFP_tADH1 | pTY025 | This work | N/A |
| sTY031 | LF library, YJL119C_yoEGFP_tADH1 | pTY026 | This work | N/A |

| | | | | |
|---|---|---|---|---|
| sTY032 | LF library, YNL217W_yoEGFP_tADH1 | pTY027 | This work | N/A |
| sTY033 | NN library, PHO5_yoEGFP_tPHO5 | pTY028 | This work | N/A |
| sTY034 | NN library, PHO89_yoEGFP_tPHO89 | pTY029 | This work | N/A |
| sTY035 | NN library, PHM6_yoEGFP_tPHM6 | pTY030 | This work | N/A |
| sTY036 | NN library, PHO84_yoEGFP_tPHO84 | pTY031 | This work | N/A |
| sTY037 | NN library, PHO11_yoEGFP_tPHO11 | pTY032 | This work | N/A |
| sTY038 | NN library, PHO12_yoEGFP_tPHO12 | pTY033 | This work | N/A |
| sTY039 | NN library, VTC1_yoEGFP_tVTC1 | pTY034 | This work | N/A |
| sTY040 | NN library, VTC3_yoEGFP_tVTC3 | pTY035 | This work | N/A |
| sTY041 | NN library, VTC4_yoEGFP_tVTC4 | pTY036 | This work | N/A |
| sTY042 | NN library, VTC2_yoEGFP_tVTC2 | pTY037 | This work | N/A |
| sTY043 | NN library, PHO8_yoEGFP_tPHO8 | pTY038 | This work | N/A |
| sTY044 | NN library, PHO86_yoEGFP_tPHO86 | pTY039 | This work | N/A |
| sTY045 | NN library, VIP1_yoEGFP_tVIP1 | pTY040 | This work | N/A |
| sTY046 | NN library, PHO81_yoEGFP_tPHO81 | pTY041 | This work | N/A |
| sTY047 | NN library, HOR2_yoEGFP_tHOR2 | pTY042 | This work | N/A |
| sTY048 | NN library, PHM8_yoEGFP_tPHM8 | pTY043 | This work | N/A |
| sTY049 | NN library, CTF19_yoEGFP_tCTF19 | pTY044 | This work | N/A |
| sTY050 | NN library, SPL2_yoEGFP_tSPL2 | pTY045 | This work | N/A |
| sTY051 | NN library, CBF1_yoEGFP_tCBF1 | pTY046 | This work | N/A |
| sTY052 | NN library, DDP1_yoEGFP_tDDP1 | pTY047 | This work | N/A |
| sTY053 | NN library, ENA1_yoEGFP_tENA1 | pTY048 | This work | N/A |
| sTY054 | NN library, ENA2_yoEGFP_tENA2 | pTY049 | This work | N/A |
| sTY055 | NN library, GDE1_yoEGFP_tGDE1 | pTY050 | This work | N/A |
| sTY056 | NN library, GIT1_yoEGFP_tGIT1 | pTY051 | This work | N/A |
| sTY057 | NN library, YAR070C_yoEGFP_tYAR070C | pTY052 | This work | N/A |
| sTY058 | NN library, YJL119C_yoEGFP_tYJL119C | pTY053 | This work | N/A |
| sTY059 | NN library, YNL217W_yoEGFP_tYNL217W | pTY054 | This work | N/A |

Table S6: sgRNA sequences.

| **Target loci** | **Landing pad sequence** |
|---|---|
| Pho4p sgRNA1 | ACTGGCGTCTTTAATCCCCG |
| Pho4p sgRNA2 | CAACTGCTACAATCAAGCCG |
| LYS2 sgRNA1 | GATAAATTCACAATGCTGAG |
| LYS2 sgRNA2 | CTAATAACAATCAACCCACG |
| VTC3 sgRNA1 | CATCAGAAAGAACCCTCAGG |
| VTC3 sgRNA2 | AGAAAGGGGTACTCAAAAGG |
| CTF19 sgRNA1 | ACGCGCACTGAAGCTACAGG |
| CTF19 sgRNA2 | ATGGGCAGAAAACAGTTACG |
| PHO84 sgRNA1 | ACTACCGTGCCAGTAAACGT |
| PHO84 sgRNA2 | GCTTGGGGTCAAATCTCCGG |
| VIP1 sgRNA1 | TGATCATCTTGAATACCGGG |
| VIP1 sgRNA2 | ATGTAAGACCCCTCCGTACG |
| CBF1 sgRNA1 | CGAGGAGCAGAACTACAGCG |
| CBF1 sgRNA2 | AGACGTCATTCCCTCCAGGG |
| YJL119C sgRNA1 | ATACAGTTGATACAGAACGT |
| YJL119C sgRNA2 | CACCACACAGGTTTTACGTG |
| VTC4 sgRNA1 | TAGGTCAAGAACCTCCAGAG |
| VTC4 sgRNA2 | CGAAGATAACGACTTCGATG |
| SPL2 sgRNA1 | TGTGACTGCGATGTTCTACG |
| SPL2 sgRNA2 | GGGTATTTACAGAGAGCCAG |
| PHO5 sgRNA1 | GTCTTAGCCAGACTGACAGT |
| PHO5 sgRNA2 | GTTACAAGCACTCAAAGTGT |
| PHO81 sgRNA1 | GTGCATTCAAAACATCAACG |
| PHO81 sgRNA2 | ACTATTGATACGTTATGGGG |
| VTC2 sgRNA1 | TCCTAAGCAAGCATATGAGG |
| VTC2 sgRNA2 | ACATTGGCGTAGACCAACAC |
| PHO8 sgRNA1 | CAAACCAATACAGACCACAG |
| PHO8 sgRNA2 | AAGTCTACTACTCTCCCCAG |
| PHM8 sgRNA1 | CCACTACGACAGACCCATCG |
| PHM8 sgRNA2 | TCCGGATCCGGTAATTTGGG |
| GIT1 sgRNA1 | ATAGAAAACACCTCTAACAG |

| | |
|---|---|
| GIT1 sgRNA2 | TAATAAAATATGATGCCGAG |
| PHO89 sgRNA1 | TAGTAAGCATTAAAACAGCG |
| PHO89 sgRNA2 | CATTGGTGGGATATCATCAG |
| PHO11 sgRNA | CCAAAAGTATCATGACAACA |
| ENA2 sgRNA | CACAGCGAAAGAGATCGCCA |
| YAR070C sgRNA | ACTTTGCACAGGAATCGTGT |
| PHO12 sgRNA | TGTGAAATAAATGACTTCTA |
| ENA1 sgRNA | CGTCCACAGATTGAAAACAG |
| PHM6 sgRNA | TACATCATCGATGCACCTCG |
| HOR2 sgRNA | ACCGTCGACGTCGAACAAAG |
| VTC1 sgRNA | AATTCAGTAAACCTACACCA |
| PHO86 sgRNA | TCTGAAAATCGACTACTAGG |
| DDP1 sgRNA | TGAAGACATGAGACCCCCTA |
| YNL217W sgRNA | AAACTGGTGGAGACAACCGA |
| GDE1 sgRNA | TGATTCGGCATCTCCGAACG |

Table S7: Oligonucleotides used for PCR or sequencing validation.

| **Oligonucleotides** | **Name** | **Source** |
|---|---|---|
| **LF lib (LYS2 locus) validation** | | |
| 5'-TATGCTGGATCTGGTAGAGTGGAG-3' | LYS2 Seq Fw | This work |
| 5'-TGCGTCAAGGGCTGAAAAGAC-3' | LYS2 Seq Rev | This work |
| **NN lib (Native locus) validation** | | |
| 5'-TGCCCACACATCTAATCAAACG-3' | CBF1 Native Fw | This work |
| 5'-TAATTCCCTCTTTTATGCTTTAGTATCG-3' | CBF1 Native Rev | This work |
| 5'-AATGGGCCCTCTCGATATTG-3' | CTF19 Native Fw | This work |
| 5'-CTCATATTAAACCCATAGGAGACG-3' | CTF19 Native Rev | This work |
| 5'-CCATTAGTAATATGGCATGGAACC-3' | DDP1 Native Fw | This work |
| 5'-TAATTCCCTCTTTTATGCTTTAGTATCG-3' | DDP1 Native Rev | This work |
| 5'-GTTTGTTAGGGCAGGGATGTAG-3' | ENA1 Native Fw | This work |
| 5'-GAAGGAAAATAAAATTCTTTCCTCTTTC-3' | ENA1 Native Rev | This work |
| 5'-CGAACTTTTCACAGGTTCTGAAC-3' | ENA2 Native Fw | This work |
| 5'-CTTTGCCTCGAGAGATAATATGGAG-3' | ENA2 Native Rev | This work |
| 5'-AACATGGTGTTCCAAAGCAC-3' | GDE1 Native Fw | This work |
| 5'-GAAAATCTTGAAAGATCTGGGTATCG-3' | GDE1 Native Rev | This work |
| 5'-GAAAGCAGAGAATCAAAAGAAGC-3' | GIT1 Native Fw | This work |
| 5'-AACATGGTGTTCCAAAGCAC-3' | GIT1 Native Rev | This work |
| 5'-GTTTGGCCTTCTCAAGTATTTTGG-3' | HOR2 Native Fw | This work |
| 5'-CTCTGGAATCTTCCCCACTG-3' | HOR2 Native Rev | This work |
| 5'-CAAATGTGCATTAGCGTGTAATG-3' | PHM6 Native Fw | This work |
| 5'-GGATGCGTTGACGCTAATG-3' | PHM6 Native Rev | This work |
| 5'-AGGTGAACGAAGAAAAAAAAAAAG-3' | PHM8 Native Fw | This work |
| 5'-ATGATGATGTTAGAGAATTCTTTCTG-3' | PHM8 Native Rev | This work |
| 5'-CATTCATCGTGGGTGTCTAATAAAG-3' | PHO11 Native Fw | This work |
| 5'-CTTGAATAAAAAGCAATGTAGAGAACAG-3' | PHO11 Native Rev | This work |
| 5'-GTGGGTGTCTAATAAAGTTTTAATGAC-3' | PHO12 Native Fw | This work |
| 5'-CCAATAATTTTGAATAAAAGAAGCAC-3' | PHO12 Native Rev | This work |
| 5'-ACGTATTTGGAAGTCATCTTATGTG-3' | Pho5 Native Fw | This work |
| 5'-ACCTGCATTAACTAAAGCAGCG-3' | Pho5 Native Rev | This work |
| 5'-GCAGACCACAGGGTAGTCAAC-3' | PHO8 Native Fw | This work |

| | | |
|---|---|---|
| 5’-AGGAAGAAGTTGGCTGGTAGATC-3’ | PHO8 Native Rev | This work |
| 5’-AAACGCGCTAATTGCATCAG-3’ | PHO81 Native Fw | This work |
| 5’-GATGGACACCTTTCCATATTG-3’ | PHO81 Native Rev | This work |
| 5’-TGCCCACACATCTAATCAAACG-3’ | PHO84 Native Fw | This work |
| 5’-TAATTCCCTCTTTTATGCTTTAGTATCG-3’ | PHO84 Native Rev | This work |
| 5’-GAGGAGAGACAACCCTGTTCTC-3’ | PHO86 Native Fw | This work |
| 5’-AGATGGTTCCGATGGTTGC-3’ | PHO86 Native Rev | This work |
| 5’-AGACCTTTTTTTTCTTTTTCTGC-3’ | Pho89 Native Fw | This work |
| 5’-GATAACTGTAAGTCAAAAGGCCATG-3’ | Pho89 Native Rev | This work |
| 5’-ATGTTTCCTACCCCAATGATGGTTCG-3’ | SPL2 Native Fw | This work |
| 5’-AGTTGGCGGCGGTCGTGG-3’ | SPL2 Native Rev | This work |
| 5’-TATTCGCATCATATATCTGAAATGTTTTAC-3’ | VIP1 Native Fw | This work |
| 5’-TTTAATAAGGCAATAATATTAGGTATGTagatatac-3’ | VIP1 Native Rev | This work |
| 5’-GGCAGAGGAGAGTTATCACTCC-3’ | VTC1 Native Fw | This work |
| 5’-CGAGCCTCTTTAACCTAATGC-3’ | VTC1 Native Rev | This work |
| 5’-ACGAGGTTCCCAGTTTCCC-3’ | VTC2 Native Fw | This work |
| 5’-ATAGAGAATAGTGGAATAAGGGTGG-3’ | VTC2 Native Rev | This work |
| 5’-GTTACATTACCGGCCACATAGTAG-3’ | VTC3 Native Fw | This work |
| 5’-TCTTTTTTTATAAATACCACTAAAACTGG-3’ | VTC3 Native Rev | This work |
| 5’-TAATTGTCATACGCATTGTTATACC-3’ | VTC4 Native Fw | This work |
| 5’-TTCCAGATTGTTTGGAATTACC-3’ | VTC4 Native Rev | This work |
| 5’-CTGAGCTTGAATCTATCGAAATG-3’ | YAR070C Native Fw | This work |
| 5’-GAAATTATATCCAACAGAAAGCTCAG-3’ | YAR070C Native Rev | This work |
| 5’-GAAAAAAAGATGTTCTTTGGTCG-3’ | YJL119C Native Fw | This work |
| 5’-CTATAATTGGTTTGACCATTTTTTC-3’ | YJL119C Native Rev | This work |
| 5’-GGAGGAGCTGGGTTATCATAAAG-3’ | YNL217W Native Fw | This work |
| 5’-CAGAACCACTGTCAACAACGAC-3’ | YNL217W Native Rev | This work |
| **GFP-tag lib (Native locus) validation** | | |
| 5’-TGCCCACACATCTAATCAAACG-3’ | CBF1pr Seq Fw | This work |
| 5’-GGAACTAGTAACAAAATACGACACG-3’ | CTF19pr Seq Fw | This work |
| 5’-GCGGCTTAATTCTCAAACGTC-3’ | DDP1pr Seq Fw | This work |
| 5’-GGTCTAGAAAAGGCTGCTCC-3’ | ENA1pr Seq Fw | This work |
| 5’-GGGCTAGGTCTAGAAAAGGC-3’ | ENA2pr Seq Fw | This work |

| | | |
|---|---|---|
| 5'-GGAGAGCTTGGGATTTACACG-3' | GDE1pr Seq Fw | This work |
| 5'-CGGTACCATTATCATCTCTCAACC-3' | HOR2pr Seq Fw | This work |
| 5'-GGATGTTCTGCCCTATAATGTTACG-3' | PHM6pr Seq Fw | This work |
| 5'-CAGAACGCCTAATCGAATCGTATTATC-3' | PHM8pr Seq Fw | This work |
| 5'-GGTATTGCCAAGAGATTAAACAAGG-3' | PHO11pr Seq Fw | This work |
| 5'-GGTATTGCCAAGAGATTAAACAAGG-3' | PHO12pr Seq Fw | This work |
| 5'-TGAATACGACACAACCTACTTGG-3' | PHO5pr Seq Fw | This work |
| 5'-AGAATTGATCATGCTGGTCACC-3' | PHO8pr Seq Fw | This work |
| 5'-GACTTCGGTAAAATTGATGGAAGC-3' | PHO81pr Seq Fw | This work |
| 5'-TGTTCTAACCATTGGGCTGC-3' | PHO86pr Seq Fw | This work |
| 5'-GCCCTTATTGTTCTGGTTAGTAATCTC-3' | PHO89pr Seq Fw | This work |
| 5'-GCAATTACAACTACAACGCCG-3' | SPL2pr Seq Fw | This work |
| 5'-CCCAAATCTTCTTCTCAATTCGATG-3' | VIP1pr Seq Fw | This work |
| 5'-TGGCAAGAGCGATTAAGTATAGC-3' | VTC1pr Seq Fw | This work |
| 5'-GGTATCTCAAGAGGAAAATGAGCG-3' | VTC2pr Seq Fw | This work |
| 5'-AGAATACTCGCCACGTTATTGC-3' | VTC3pr Seq Fw | This work |
| 5'-CAGGAAACCTCCACTACCAAC-3' | VTC4pr Seq Fw | This work |
| 5'-TGCTGGGTGACTTTATTCACAAG-3' | YNL217Wpr Seq Fw | This work |
| 5'-CACTCTTGAAAAAGTCATGCCG-3' | GFP Seq Rev | This work |

Table S8: Media and plate recipes.

| **Ingredient** | **Amount** |
|---|---|
| **Synthetic complete, phosphate-rich media (per liter)** | |
| Glucose | 20 g |
| YNB with ammonium sulfate, without phosphates | 5.6 g |
| Complete Supplement Mixture | 790 mg |
| Adenine hemisulfate salt | 163.5 mg |
| L-Tryptophan | 50 mg |
| Monobasic potassium phosphate | 1 g |
| Sodium chloride | 0.1 g |
| **Synthetic complete, phosphate-free media (per liter)** | |
| Glucose | 20 g |
| YNB with ammonium sulfate, without phosphates | 5.6 g |
| Complete Supplement Mixture | 790 mg |
| Adenine hemisulfate salt | 163.5 mg |
| L-Tryptophan | 50 mg |
| Potassium chloride | 547.8 mg |
| Sodium chloride | 0.1 g |
| **Synthetic complete, uracil dropout plate (per liter)** | |
| Glucose | 40 g |
| Yeast synthetic dropout supplement, without uracil | 1.92 g |
| YNB without Amino Acids | 6.7 g |
| Agar | 40 g |
| Note: 20 mL per plate | |
| **Synthetic complete, uracil dropout plate (per liter)** | |
| Glucose | 40 g |
| Yeast synthetic dropout supplement, without uracil | 1.92 g |
| YNB without Amino Acids | 6.7 g |
| Agar | 40 g |
| Uracil (760 $\mu$g/mL) | 66 mL |
| 5-FOA (10%) | 10 mL |
| Note: 20 mL per plate | |

Table S9: MICROSTAR mold fabrication: coating and soft bake parameters

| | Spin coating | | | | Soft bake | | | |
|---|---|---|---|---|---|---|---|---|
| # | **Layer** | **Resist** | **Speed** | **Time** | **PB temp.** | **PB time** | **SB temp.** | **SB time** |
| **f1** | sieve | SU8-2 | 2500 rpm | 30s | 65°C | 1min | 95°C | 1min |
| **f2** | chamber | SU8-2 | 1000 rpm | 30s | 65°C | 1min | 95°C | 3min |
| **f3** | dg | SU8-3025 | 1875rpm | 40s | none | none | 95°C | 13.3min |
| **f4** | flow | AZ 10XT-60 | 1000rpm | 40s | none | none | 105°C | 6.5min |
| **c1** | control | SU8-3025 | 4144rpm | 40s | none | none | 95°C | 10min |

Table S10: MICROSTAR mold fabrication: exposure and post exposure bake parameters

| | Exposure | | | | Post exposure bake | | | |
|---|---|---|---|---|---|---|---|---|
| # | **Layer** | **Power** | **Time** | **Dose** | **PB temp.** | **PB time** | **PEB temp.** | **PEB time** |
| **f1** | sieve | 20mW/cm2 | 3.2s | 64mJ/cm2 | 65°C | 1min | 95°C | 1min |
| **f2** | chamber | 20mW/cm2 | 6.4s | 128mJ/cm2 | 65°C | 1min | 95°C | 1min |
| **f3** | dg | 20mW/cm2 | 10.8s | 216mJ/cm2 | 65°C | 1min | 95°C | 4.3min |
| **f4** | flow | 20mW/cm2 | 15x10s | 3000mJ/cm2 | none | none | none | none |
| **c1** | control | 20mW/cm2 | 7.5s | 150mJ/cm2 | 65°C | 1 | 95°C | 3min |

Table S11: MICROSTAR mold fabrication: development and hard bake parameters.

| | Exposure | | | | | Hard bake | | |
|---|---|---|---|---|---|---|---|---|
| # | **Layer** | **Developer** | **Time** | **Rinser** | **Rinsing time** | **HB temp.** | **HB time** | **Final height** |
| **f1** | sieve | none | none | none | none | none | none | 1.2 $\mu$m |
| **f2** | chamber | PGMEA | 1min | 2-propanol | 1min | 135 | 2 hrs | 3.8 $\mu$m |
| **f3** | dg | PGMEA | 2x3.5min | 2-propanol | 1min | 135 | 2 hrs | 40 $\mu$m |
| **f4** | flow | AZ400K 1:3.5 | 12min | DI water | 1min | 115 | 1 hrs | 15 $\mu$m |
| **c1** | control | PGMEA | 2x2.5min | 2-propanol | 1min | 135 | 2 hrs | 20 $\mu$m |

Table S12: MCA mold fabrication: coating and soft bake parameters

| # | Spin coating | | | | Soft bake | | | | | |
|---|---|---|---|---|---|---|---|---|---|---|
| | **Layer** | **Resist** | **Speed** | **Time** | **Start temp.** | **Ramp time** | **Step temp.** | **Step time** | **End temp.** | **Ramp time** |
| **f1** | sieve | GM1040 | 3616 rpm | 40s | 30°C | 20min | 130°C | 5min | 30°C | 20min |
| **f2** | chamber | GM1050 | 1803 rpm | 40s | 30°C | 25min | 130°C | 5min | 30°C | 25min |
| **f3** | dg | GM1070 | 1933rpm | 40s | 30°C | 50min | 130°C | 5min | 30°C | 50min |
| **f4** | flow | AZ 10XT-60 | 1000rpm | 40s | none | none | 105°C | 6.5min | none | none |
| **c1** | control | GM1060 | 1793rpm | 40s | 30°C | 30min | 130°C | 5min | 30°C | 30min |

Table S13: MCA mold fabrication: exposure and post exposure bake parameters

| # | Exposure | | | | Post exposure bake | | | | | |
|---|---|---|---|---|---|---|---|---|---|---|
| | **Layer** | **Resist** | **Time** | **Dose** | **Start temp.** | **Ramp time** | **Step temp.** | **Step time** | **End temp.** | **Ramp time** |
| **f1** | sieve | 20mW/cm2 | 3s | 60mJ/cm2 | 30°C | 20min | 90°C | 15min | 30°C | 90min |
| **f2** | chamber | 20mW/cm2 | 6.4s | 128mJ/cm2 | 30°C | 20min | 90°C | 20min | 30°C | 90min |
| **f3** | dg | 20mW/cm2 | 16.6s | 332mJ/cm2 | 30°C | 40min | 90°C | 40min | 30°C | 90min |
| **f4** | flow | 20mW/cm2 | 15x10s | 3000mJ/cm2 | 30°C | none | none | none | none | none |
| **c1** | control | 20mW/cm2 | 9.4s | 188mJ/cm2 | 30°C | 30min | 90°C | 30min | 30°C | 90min |

Table S14: MCA mold fabrication: development and hard bake parameters

| # | Development | | | | | Hard bake | | |
|---|---|---|---|---|---|---|---|---|
| | **Layer** | **Developer** | **Time** | **Rinser** | **Rinsing time** | **HB temp.** | **HB time** | **Final height** |
| **f1** | sieve | none | none | none | none | none | none | 1.5 $\mu$m |
| **f2** | chamber | PGMEA | 1+0.5min | 2-propanol | 1min | 135 | 2 hrs | 4.2 $\mu$m |
| **f3** | dg | PGMEA | 2+1min | 2-propanol | 1min | 135 | 2 hrs | 40 $\mu$m |
| **f4** | flow | AZ400K 1:3.5 | 12min | DI water | 1min | 115 | 1 hrs | 15 $\mu$m |
| **c1** | control | PGMEA | 2x1min | 2-propanol | 1min | 135 | 2 hrs | 12 $\mu$m |

[6] Stuart Berg, Dominik Kutra, Thorben Kroeger, Christoph N. Straehle, Bernhard X. Kausler, Carsten Haubold, Martin Schiegg, Janez Ales, Thorsten Beier, Markus Rudy, Kemal Eren, Jaime I. Cervantes, Buote Xu, Fynn Beuttenmueller, Adrian Wolny, Chong Zhang, Ullrich Koethe, Fred A. Hamprecht, and Anna Kreshuk. ilastik: interactive machine learning for (bio)image analysis. *Nature Methods*, September 2019.

[7] Nicolas Denervaud, Johannes Becker, Ricard Delgado-Gonzalo, Pascal Damay, Arun S. Rajkumar, Michael Unser, David Shore, Felix Naef, and Sebastian J. Maerkl. A chemostat array enables the spatio-temporal analysis of the yeast proteome. *Proceedings of the National Academy of Sciences*, 110(39):15842–15847, 2013.

[8] Evert Bosdriesz, Meike T. Wortel, Jurgen R. Haanstra, Marijke J. Wagner, Pilar de la Torre Cortés, and Bas Teusink. Low affinity uniporter carrier proteins can increase net substrate uptake rate by reducing efflux. *Scientific Reports*, 8(1):5576, 2018.
